# Bayesian reassessment of the epigenetic architecture of complex traits

**DOI:** 10.1101/450288

**Authors:** Daniel Trejo Banos, Daniel L. McCartney, Tom Battram, Gibran Hemani, Rosie M. Walker, Stewart W. Morris, Qian Zhang, David J. Porteous, Allan F. McRae, Naomi R. Wray, Peter M. Visscher, Chris S. Haley, Kathryn L. Evans, Ian J. Deary, Andrew M. McIntosh, Riccardo E. Marioni, Matthew R. Robinson

## Abstract

Epigenetic DNA modification is partly under genetic control, and occurs in response to a wide range of environmental exposures. Linking epigenetic marks to clinical outcomes may provide greater insight into underlying molecular processes of disease, assist in the identification of therapeutic targets, and improve risk prediction. Here, we present a statistical approach, based on Bayesian inference, that estimates associations between disease risk and all measured epigenetic probes jointly, automatically controlling for both data structure (including cell-count effects, relatedness, and experimental batch effects) and correlations among probes. We benchmark our approach in simulation study, finding improved estimation of probe associations across a wide range of scenarios over existing approaches. Our method estimates the total proportion of disease risk captured by epigenetic probe variation, and when we applied it to measures of body mass index (BMI) and cigarette consumption behaviour in 5,101 individuals, we find that 66.7% (95% CI 60.0-72.8) of the variation in BMI and 67.7% (95% CI 58.4-76.9) of the variation in cigarette consumption can be captured by methylation array data from whole blood, independent of the variation explained by single nucleotide polymorphism markers. We find novel associations, with smoking behaviour associated with a methylation probe at the MNDA gene with >95% posterior inclusion probability, which is a myeloid cell nuclear differentiation antigen gene previously implicated as a biomarker for inflammation and non-Hodgkin lymphoma risk. We conduct unique genome-wide enrichment analyses, identifying blood cholesterol, lipid transport and sterol metabolism pathways for BMI, and response to xenobiotic stimulus and negative regulation of RNA polymerase II promoter transcription for smoking, all with >95% posterior inclusion probability of having methylation probes with associations >1.5 times larger than the average. Finally, we improve phenotypic prediction in two independent cohorts by 28.7% and 10.2% for BMI and smoking respectively over a LASSO model. These results imply that probe measures may capture large amounts of variance because they are likely a consequence of the phenotype rather than a cause. As a result, ‘omics’ data may enable accurate characterization of disease progression and identification of individuals who are on a path to disease. Our approach facilitates better understanding of the underlying epigenetic architecture of complex common disease and is applicable to any kind of genomics data.

## 2 Introduction

Data characterizing gene expression, protein structure, or epigenetic modifications such as DNA methylation, histone marks and nucleosome positioning are becoming increasingly available. Epigenetic marks reflect a wide range of environmental exposures and genetic influences, are critical for regulating gene and non-coding RNA expression [1], and have been shown to be associated either as a cause or consequence with disease [2]. The identification of clinically relevant epigenetic loci can provide insight into the molecular underpinning of disease [3], leading to identification of biologically relevant therapeutic targets [4] and potentially epigenetic-guided clinical decision making [5].

Most studies testing for association between ‘omics’ data and complex traits utilize methodology from genome-wide associations studies, meaning that probe effects are tested one at a time [6]. This methodology does not account for correlations among probes and leads to model overfitting, poor effect size estimation, and poor calibration of prediction due to omitted variable bias [7]. Additionally, data structure such as intra-sample cellular heterogeneity, sample relatedness, population stratification, or experimental design effects are a major challenge [8] and result in more cross-chromosome correlation than genetic data. This structure, in conjunction with the fact that cases and controls typically differ in their cell-type composition, can result in spurious associations and many statistical algorithms have been proposed to tackle these potential biases [7, 9, 10]. However, all current statistical approaches rely upon corrections for structure that require a choice of either a suitable reference profile of representative cell types, or a limited number of pre-selected variables computed from the methylation data (e.g. Surrogate Variable Analysis [9], RefFreeEWAS [11], EWASher [12], or ReFACTor [13]), with the underlying assumption that all confounders are reflected by a sparse set of latent covariates and methylation sites.

Here, we present an alternative approach, based on Bayesian inference, that: (i) estimates probe effects on an outcome jointly and conditionally on each other whilst controlling for other covariates such as sex and age, avoiding model over-fitting and controlling for both data structure (including cell-count effects) and the correlations among probes; (ii) does not require any knowledge of cell-type composition or any selection of proxy confounder variables (i.e., accounts for both known and unknown confounders); (iii) estimates the total proportion of disease risk accounted for by the probe effects (cumulative proportion of variance explained); (iv) estimates probe effects conditional on other sources of data such as single nucleotide polymorphism data, enabling a determination of the unique contribution of different data types; (v) gives an in-depth understanding of the genome-wide range of probe effects on the phenotype, in terms of the likely number of independent effects and their variance explained; (vi) enables unique genome-wide enrichment analyses, describing the variance explained and number of trait-associated probes of each annotation; and (vii) provides improved estimation of biomarker effects, which could be used for disease risk assessment. The approach is similar to, but more flexible than linear mixed model analyses recently proposed in genome-wide association studies [14], and we demonstrate properties (i) through (vii) with theory, simulation and then empirical analysis of body-mass index (BMI) and smoking behaviour for over 5000 individuals with methylation probe measures [15].

## 3 Results

### 3.1 Methods overview

Our approach assumes that the observed phenotype **y** is reflected by a linear combination of genetic effects (*β_G_*) estimated from single nucleotide polymorphism (SNP) data, epigenetic effects (*β_cpg_*) estimated from probes on an array, along with age and sex specific effects (*α, γ*), such that:

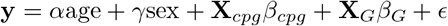

with the effects *β_G_*, *β_cpg_*, *α* and *γ* being estimated in a Bayesian statistical model. We assign to each type of effect a prior distribution, which is a mixture of Gaussian distributions and a discrete "spike" at zero. This allows for non-identifiable effects to be excluded from the model, while the rest are estimated jointly (and conditionally on each other given our algorithm). Controlling for all different covariates and their effects while estimating each individual effect, better alleviates problems related to correlations and structure in the marker data as we show in our simulation study (**fig.1**, also see Methods). This is because each probe estimate is made conditionally after accounting for the SNP markers and the other ‘omics’ probes in the model, which capture, and thus account for, genetic relationships in the data [16] and capture structural effects (such as cell-count effects, experimental batch effects, or population structure).

**Figure 1.**
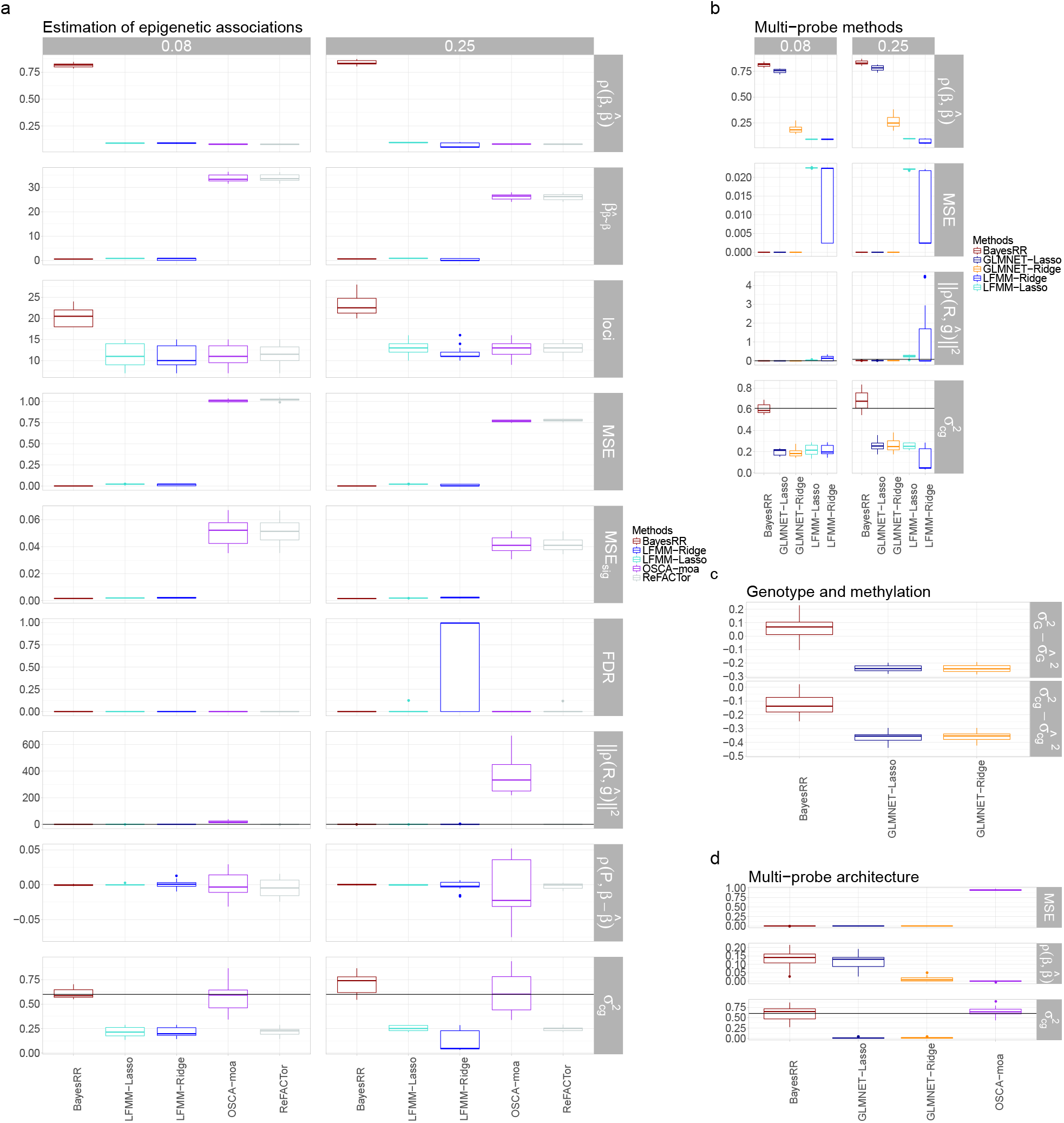
Simulation study. **a**. Estimation of phenotype-epigenetic associations using five recent approaches (see Methods), where probes are associated with cell-type proportion variation and the norm of the correlation vector between the phenotype and the cell-type proportions have two different values either 0.08 or 0.25. Row panels provide results for different metrics of performance: the correlation between true effects and estimates (*ρ*(*β, β̂*)), the slope of a regression of the estimates on the true effects (*β_β̂_*~_*β*_), the number of genome-wide significant probes identified (loci), the mean square error (MSE), the MSE of the genome-wide significant probes (*MSE_sig_*), the false discovery rate (FDR), the norm of the correlation vector between a individual-level predictor made from the probe effects and the cell-type proportions (||*ρ*(**R**, **ĝ**)||), the correlation between the first principal component of the probe data and the difference between the estimated and true effect (||*ρ*(**P**, **R**)||) and the phenotypic variance attributable to the probes 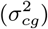. Black lines give the true value across panels. **b**. Comparison of BayesRR with just the methods which fit probes jointly (multi-probe methods) either accounting for latent factors (LFMM-Lasso and LFMM-Ridge) or not (GLMNET-Lasso and GLMNET-Ridge). **c**. Simulation results of methylation marker effects for a phenotype influenced by both 100 methylation probes and 1000 SNP markers, showing the difference between the true and the estimated phenotypic variance explained by genetic markers 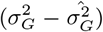 and epigenetic probes 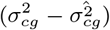. **D** Comparisons of approaches that do not fit latent factors within the model when the underlying epigenetic architecture is less sparse (phenotype is influenced by 1000 probes, rather than only 100).

As described above, we fit both SNP and epigenetic probe data together, as it allows a determination of the unique contribution of each data source. However, our software implementation of this modelling framework, (BayesRR, which is freely available, see URLs) is entirely flexible. Any number of data sources can be modelled together, each with separate mixtures, making it applicable to any kind of genetic or epigenetic data. Furthermore, if only epigenetic probe data are available, estimates of the probe effects can still be obtained jointly, avoiding model over-fitting and controlling for both data structure (including cell-count effects) and correlations among probes (**fig.1**, also see Methods). Additionally, other major covariates could be included, including genetic loci of large effect such as HLA in immunodisease or APOE4 variant in Alzheimer disease, or latent factors can still be fit alongside alongside the probe data. This flexibility is important as data sets will likely be variable in their structure and the degree to which different ‘omics’ measures are correlated.

### 3.2 Simulation study

We simulated methylation data for 2000 individuals at 103,638 probes. We reproduced cell-type proportion variation present in real methylation probe data, using a recently proposed simulation model [13]. Our first simulation scenario, was a "sparse" setting, where a phenotype is determined by 100 differentially methylated probes (see Methods in **sec.5.1.**), which cumulatively explained 60% of the phenotypic variance in the trait. We focus in the main text on two scenarios where probes are associated with cell-type proportion variation and the norm of the correlation vector between the phenotype and the cell-type proportions is either 0.08 or 0.25. These scenarios reflect different degrees of confounding between phenotype and cell-type proportions. We then conduct additional simulations with a wide a range of settings, varying the cell-type proportions, the proportions of differentially methylated probes, the variance of differentially methylated probes, and the variance of the measurement noise and we present these within the Supplementary Material (see Methods and fig.S1 to fig.S4).

We benchmarked our BayesRR approach against four recently proposed methods: single-probe least-squares regression which estimates probe associations one-by-one whilst correcting for sparse latent factors to control for cell proportion confounding (ReFACTor [13]), single-probe mixed linear model association test which estimates probe associations one-by-one conditional on a relationship matrix estimated from the probe data (OSCA-moa [14]), a multi-probe ridge regression which estimates all probe associations jointly and conditionally on latent factors (LFMM2-ridge [17]), and a multimarker LASSO model which estimates all probe associations jointly and conditionally on latent factors (LFMM2-lasso [17]). Our BayesRR approach outperforms these approaches as it estimates phenotype-probe associations more accurately with higher correlation of the estimated effects with the true simulated values and with lower mean-square error (MSE, **fig.1a**). This results in almost twice the number of methylome-wide significant discoveries at >95% posterior inclusion probability within this data, whilst controlling for cell-type proportion confounding and maintaining a false discovery rate of much less than 5% (**fig.1a**). BayesRR controls for cell proportion confounding without a requirement for the addition of latent factors within the model, as evidenced by: (i) the accurate effect size estimates with reduced MSE; (ii) no inflation of the norm of the correlation vector between a individual-level predictor made from the probe effects and the cell-type proportions, and (iii) no correlation between the first principal component of the probe data and the difference between the estimated and true effect, despite significant cell-type proportion confounding within the simulated data (**fig.1a**).

This is further evidenced by comparing LASSO and ridge regression with latent factors implemented in LFMM [17], to LASSO and ridge regression without latent factors as implemented in glmnet [18], where we find that that power is increased, phenotype-probe associations are better estimated, and cell-type confounding is controlled by the models that do not fit latent factors (**fig.1b**). This is because there are probes in this situation setting which both influence the phenotype and are associated with cell-type proportions, and thus by removing variation associated with leading latent factors of the data, power to detect these probes and estimate their effects accurately is reduced. In this setting, approaches that estimate phenotype-probe association one-by-one such as ReFACTor and OSCA-moa, do not control for correlations in probe effects across the genome, resulting in increased MSE and erroneous correlations between probe effect estimates and cell-type proportion confounding (**fig.1a**). Having shown that multi-probe methods remove the necessity for latent factor correction and that BayesRR performs better than other multi-probe approaches (**fig.1a** and **b**), we then conduct our remaining benchmarking of BayesRR against LASSO and ridge regression without latent factor correction as implemented in glmnet, finding the exact same increased performance of BayesRR irrespective of the variance of the cell-type proportions, the proportions of differentially methylated probes, the variance of differentially methylated probes, and the variance of the measurement noise (see supplementary fig.S1 to fig.S4).

BayesRR also provides accurate estimation of the total proportion of phenotypic variance explained by the probes, represented by the panel 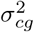 in **fig.1a** across different scenarios. With the exception of a mixed linear model as implemented in restricted effects maximum likelihood within OSCA [14] (OSCA-moa), all other approaches only enable estimation of the proportion of variance attributable probes identified as genome-wide significant and thus do not provide an estimate of the total association between the phenotype and the probe data (**fig.1a**). We examined whether methylation probe effects can be estimated conditionally on the SNP marker effects to determine the unique contribution of each type of marker. We simulated correlated genetic and epigenetic effects, with 100 epigenetic effects drawn from a normal distribution 𝒩(0, 0.5/100), and a combination of 100 genetic effects drawn from a normal distribution 𝒩 (0, 0.2/100) and 900 smaller genetic effects drawn from 𝒩 (0, 0.01/900). We find that BayesRR can better distinguish between the variance explained by genetic markers and methylation probes, as compared to LASSO or ridge regression implemented in glmnet, but with higher variability in the error of estimates as compared to when only estimating phenotypic variance associated with methylation probe (**fig.1c** and fig.S4).

We then compared BayesRR to other multi-probe approaches that do not fit latent factors within the model across two levels of sparsity, the first where a phenotype is influenced by 100 probes (**fig.1a**) and the second where a phenotype is influenced by 1000 probes (**fig.1d**). Both the mixed linear model and BayesRR provide unbiased estimates of the proportion of phenotypic variance captured by the probes, with the error variance of each approach dependent upon the underlying effect size distribution (**fig.1a** and **d**). Again however, estimated effects from the other multi-probe approaches showed reduced correlation with the true effects and higher MSE as compared to BayesRR, demonstrating that BayesRR will provide improved performance in both sparse and non-sparse regimes (**fig.1d**).

In the following empirical analysis we also illustrate the additional advantages of working with a posterior distribution to elucidate the architecture of epigenetic effects whilst accounting for the uncertainty of our model given the phenotypic data.

### 3.3 Application to body mass index and smoking

We applied our BayesRR approach to two lifestyle factors, smoking and body mass index, that are correlated with numerous health outcomes across the lifecourse. Previous studies have shown that smoking produces a strong alteration in methylation levels which at the same time, are related to the etiology of smoking-related disease [19], and BMI has also been associated with methylation levels and adipose-related traits [20]. Here, we present results from a converged set of four models for each trait, each model having different starting values, applied to 5,101 individuals of the Generation Scotland cohort (see Methods in **sec.5.5.1**).

Our approach facilitates the estimation of the total phenotypic variation attributable to the SNP markers and methylation probes (**fig.2** and summary in **table S1** and **table S2**). For BMI, 66.74% (95% CI 59.99-72.80) of the phenotypic variance of BMI was captured by methylation probes, with 65.47% (95% CI 53.75-77.28) of this attributable to 143.66 (95% CI 116.25-168.10) probes that each explain <1% of the phenotypic variance. The remaining third of the phenotypic variance captured by methylation probes, was attributable to 10 probes with 95% inclusion probability (IP), which cumulatively explain 17.11% (95% CI 13.18-21.16) of the phenotypic variance of BMI **fig.2b**. This suggests that most epigenetic probe effects for BMI are relatively small, but larger than SNP marker effects, which cumulatively capture an additional 19.89%(CI 13.69-25.57) of the phenotypic variance **fig.2a**. In total, the variance captured by both methylation probes and SNP markers was estimated as 86.63% (95% CI 73.68-98.37). We repeated the analysis but excluded close relatives and modelled only the methylation probe effects, finding 69.55% (95% CI 57.35-78.32) of the phenotypic variance in the remaining 2614 unrelated individuals was associated with methylation probe variation. This implies that our estimates of the variance captured by methylation probes, are independent of the variance attributable to SNP markers and do not reflect family effects.

**Figure 2.**
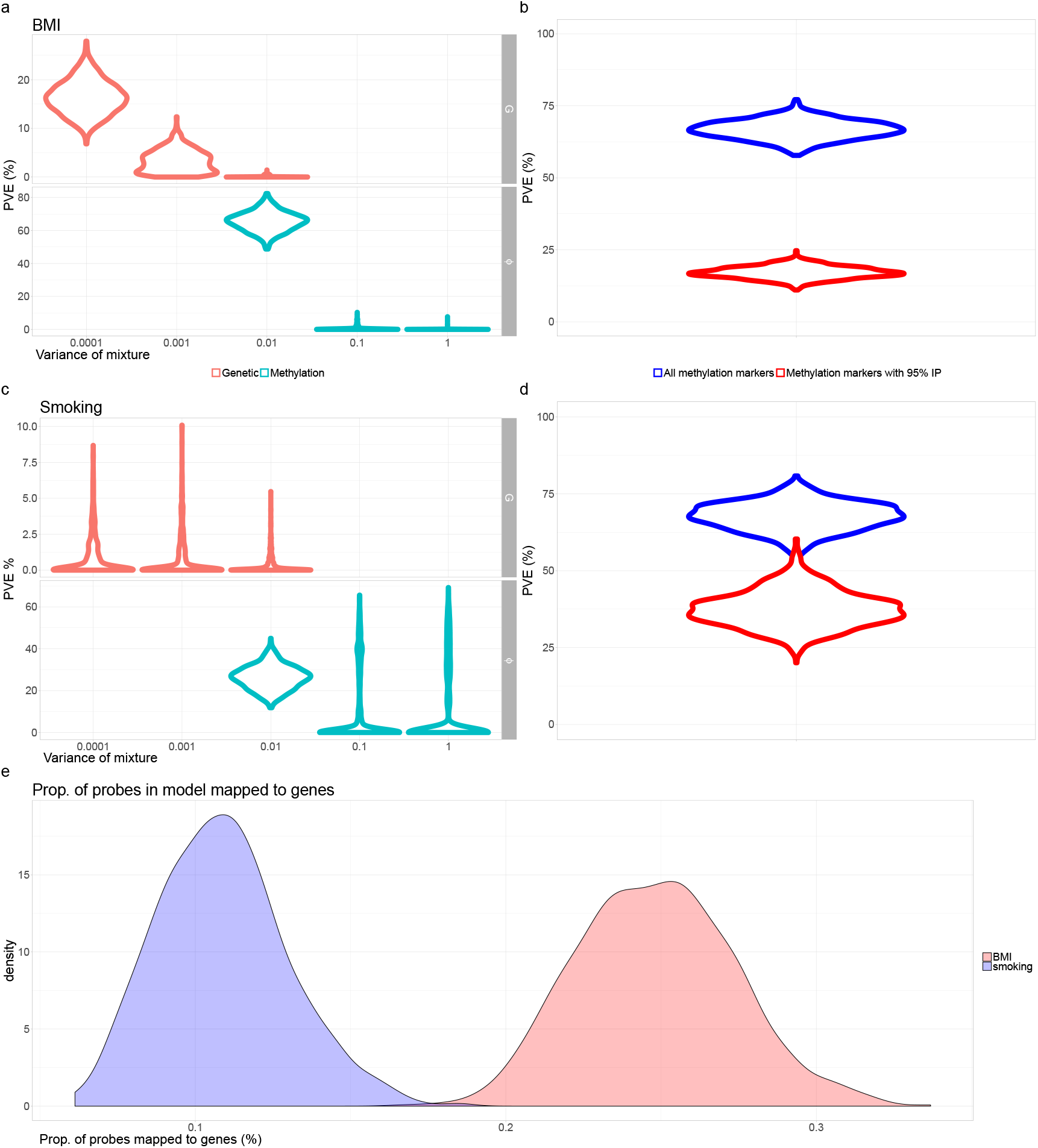
Biomarker architecture. **a**. Phenotypic variance of BMI explained by the three mixtures for single nucleotide polymorphism markers (SNP; Genetic) and methylation probes, with mixture variances (0.0001,0.001,0.01) and (0.01,0.1,1) respectively. **b**. For BMI, the phenotypic variance explained by all markers in the model (blue) and for the markers with 95% posterior inclusion probability (IP; red) is shown. **c**. For cigarette consumption, the phenotypic variance explained by the mixtures for the SNPs and methylation probes, with same mixture specific variances as for BMI. **d**. Phenotypic variance explained by all markers in the model (blue) and variance explained by the markers with 95%IP for cigarette consumption. **e**. Distribution of proportion of all methylation probes in model for BMI (red) and smoking(blue).

For smoking behaviour, defined as the number of pack years, we find that 67.68% (95% CI 58.41-76.86) of phenotypic variance is captured by methylation probes **fig.2**. In contrast to BMI, we find evidence for 5 probes with 95%IP, which capture almost a half (37.26%, 95% CI 27.01-49.5) of the phenotypic variance **fig.2**. The remaining 26.3% (95% CI 15.27-37.10) of phenotypic variance was attributable to 56.33 (95% CI 35-77) methylation probes of effect size <1% **fig.2**. Of the probes mapped to genes, 0.25% (95% CI 0.20-0.30) are in the model for BMI, more than double of the 0.11% (95% CI 0.07-0.15) for smoking behaviour **fig.2**. In total, the variance captured by both methylation probes and SNP markers was estimated as 69.54% (95% CI 58.41-84.21). Again, we repeated the analysis but excluded close relatives and modelled only the methylation probe effects, finding 73.83% (95% CI 54.33-88.26) of the phenotypic variance was captured by methylation probe variation, which again implies that our estimates are independent of the variance attributable to SNP markers and do not reflect family effects. We also used a linear mixed effects model where probe values are used to calculate a co-variance matrix, which is then used in a restricted maximum likelihood estimation (REML) approach [14], but this approach did not produce a converged set of estimates for either phenotype, with or without relatives in the data. Taken together, these results highlight the ability of our approach to describe the architecture of epigenetic associations, in terms of the likely number and effect size of associated probes, and our results imply a higher degree of polygenicity of epigenetic probe associations, spread throughout the genome, for BMI as opposed to smoking behaviour.

We then proceeded to derive annotation information from the posterior distribution over effects. First, we looked for Gene Ontology (GO) enrichment for the probes whose IP was >95% and compared these to previous findings using the EWAS catalog. We replicate previous associations with BMI and smoking, finding enrichment for probes with >95%IP corresponding to GO terms of thrombin-related pathways for smoking(**table S7**) and lipid transport for BMI(**table S6**). We find two additional related terms for BMI of lipids and triglycerides (**fig.3a**), and one additional probe for smoking behaviour that has not been previously associated to any trait in the EWAS catalogue, probe *cg16663980* that is annotated for the *MNDA* gene, a myeloid cell nuclear differentiation antigen previously implicated as a biomarker for inflammation and non-Hodgkin lymphoma risk (see Methods **sec.**m:enrich; **fig.3b**). For both traits, probes with >95% IP were also associated with alcohol consumption.

**Figure 3.**
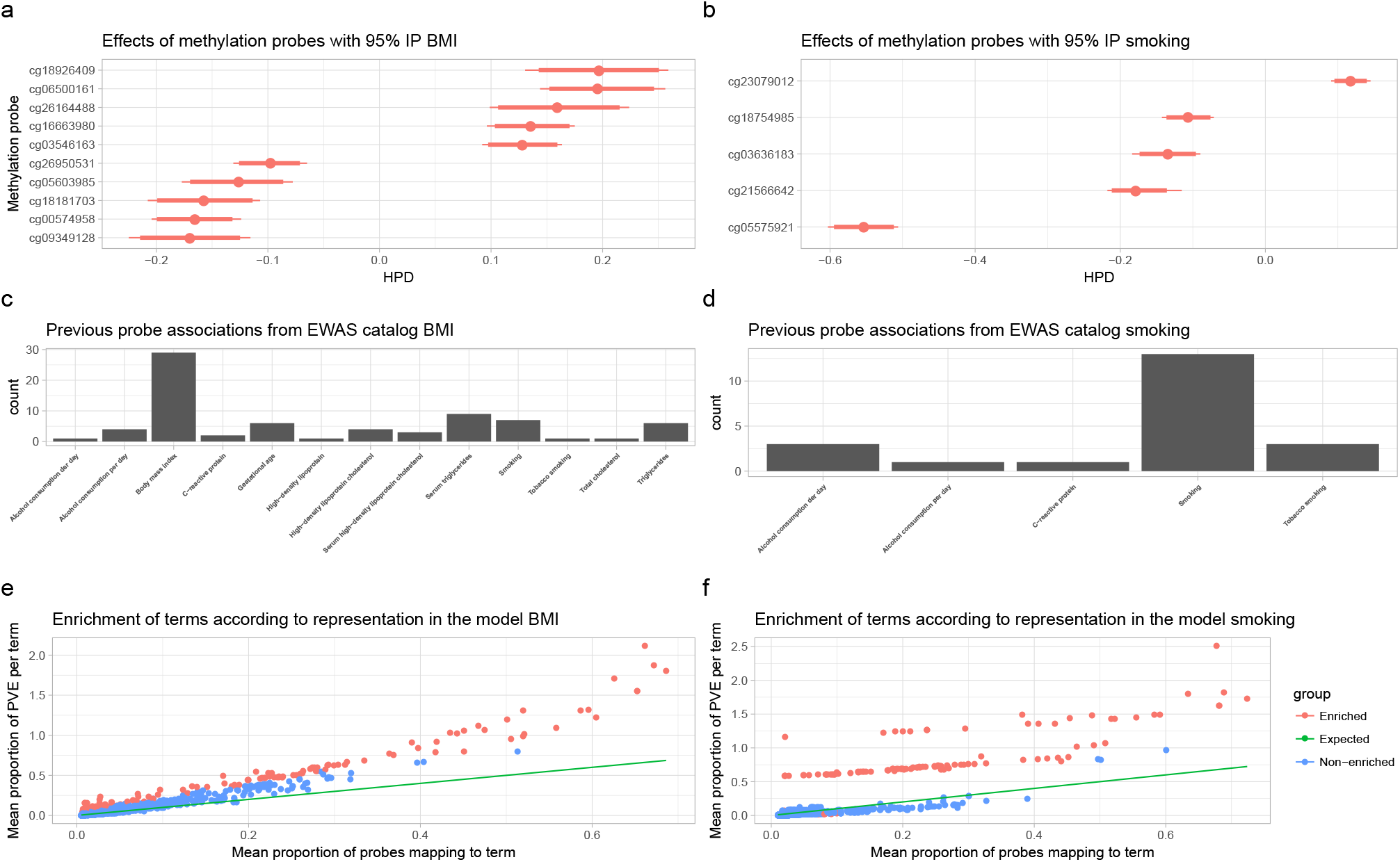
Annotation replication and enrichment analysis: **a**., **b**. Posterior distribution of effect sizes for methylation probes with 95% posterior inclusion probability (IP) for BMI and smoking respectively. **c**.,**d**. Previous associations found for the probes with 95% IP according to the EWAS catalog. **e**.,**f**. Results of a whole-genome enrichment analysis of all methylation probes in the model. Proportion of phenotypic variance explained by methylation probes is plotted against the proportion of probes mapping to each enrichment term tested (BMI left, smoking right). The one-to-one line represents the expected value of proportion of phenotypic variance explained if all probes explained the same amount of variance among annotation terms. Significantly enriched terms are those defined as having both IP >95% and enrichment of >95% IP of being >1.5.

We then take advantage of the fact that if we use the posterior distribution over all effects, we can derive a posterior distribution over GO terms and devise a definition of enrichment (see Methods **sec.5.5.2**). Under our enrichment statistic, we can measure those GO terms which explain a greater proportion of phenotypic variance than expected, given the proportion of probes that map to the GO term (**fig.3e** for BMI,**fig.3f** for smoking). Then, using a ROPE decision rule [21], we can define a term as being significantly enriched if the IP of the GO term in the model is >95%, and if 95% of the posterior distribution of enrichment is outside the interval (0.5, 1.5). We sorted significantly enriched terms by their mean enrichment and generated a tree map of the terms using REVIGO [22].

For BMI, there is a preponderance of lipid transport, cholesterol transport and sterol metabolism(**fig.S13**), and for smoking, response to xenobiotic stimulus and interestingly negative regulation of transcription from RNA polymerase II promoter (**fig.S14**). Taken together, this demonstrates the novel findings and additional inference that can be obtained from conducting whole-genome enrichment analyses, rather than testing for enrichment at only those effects that are singularly found to be above a significance threshold.

We repeated our whole-genome enrichment, mapping probes to genes that are differentially expressed across tissues, to ask whether associations between phenotype and methylation assessed in whole blood can be linked to tissue specific gene expression. We selected genes in the upper 99% percentile (those differentially over-expressed) and the lower 1% percentile (those differentially repressed) of those that show differential expression across tissues in the GTEX [23] and Depict [24] data (see Methods **sec.5.5.2**). We find only one significantly enriched term for BMI in GTEX repressed genes: *"Cells Transformed fibroblasts"*, one significantly enriched term in DEPICT over-expressed genes: *"KEGG ABC TRANSPORTERS"*, and one significantly enriched term for repressed genes in DEPICT: *"KEGG TRYPTOPHAN METABOLISM"*; with no term significantly enriched for smoking. These results suggest that while we can capture a large amount of phenotypic variance from methylation in whole blood, the probes of strongest effect may only reflect specific processes in the tissues sampled that are a consequence of differences among individuals in smoking behaviour or BMI.

Finally, we use the estimated methylation probe effects to predict BMI and smoking behaviour in two independent cohorts: The Lothian Birth Cohort 1936 (LBC) and the Accessible Resource of Integrated Epigenomics Studies (ARIES) dataset (see Methods **sec.5.5.4** and **sec.5.5.5**). We compared the prediction accuracy gained from our approach to recently obtained estimates from another model based on the LASSO estimator by using the *R*^2^ metric. For BMI, we achieve *R*^2^ of 15.7% for adult BMI in LBC, and in the ARIES dataset we achieve *R*^2^ of 3.56% for birth weight, 0.81% for BMI at age 7, 6.27% for BMI age 15, 11.65% for BMI in adult males, and 16.26% for BMI in adult females **table 1**. The increase in prediction accuracy with age in the ARIES results implies that methylation associations are a consequence of BMI, rather than a cause. Overall, these values amount to an improvement of 28.7% in comparison to the LASSO predictor in the LBC. For smoking, we observed values of 46.49% for LBC and 34.66% for males in ARIES, a 10.2% improvement over the LASSO predictor **table 1**. This replicates previous results showing that methylation profiles predicted BMI independently of genetic profiles in an additive manner [25] and shows that our approach can better capture the overall distribution of effects, including small effects, whilst accurately estimating larger effects, leading to improved phenotypic prediction.

Following [26], the squared correlation between a phenotype in an independent sample and a predictor of the phenotype can be approximated given the sample size of the initial study, the expected variance explained by the covariates, and the effective number of independent covariates. Assuming an effective number of covariates of 20,000 (approximate number of protein coding genes) and a variance explained of 70%, an *R*^2^ of 12% is expected. If the initial study sample size increased to 100,000 individuals than an *R*^2^ of over 60% is expected, which in combination with SNP array data, would lead to a predictor of BMI with an *R*^2^ of nearly 80% from a single blood test.

The theory described above does not account for the fact that ‘omics’ probe variation may be a consequence of the phenotype and thus in a regression equation, phenotypic variance will appear on both sides. If the consequential effects are large then it is conceivable that all phenotypic variance may be attributable to probe effects within this type of analysis. Additionally, given the highly variable nature of ‘omics’ measures, and the considerable changes with age that occur [27], it may also may also be likely that the phenotypic variance attributable to the probes differs across cohorts.

**Table 1.** Replication study: A phenotypic predictor was created in the Lothian Birth Cohort data 1936 (LBC1936) and Accessible Resource of Integrated Epigenomics Studies (ARIES) cohorts from the methylation effects estimated in the Generation Scotland (GS) data. The prediction accuracy as measured by the *R*^2^ statistic is presented as compared to LASSO estimates of the methylation effects. In the ARIES cohort, A.0 refers to measures at birth, A.7 refers to measures at age 7, A.15 refers to measures at age 15, A.FOF refers to adult males (fathers), and A.FOM refers to females (mothers).

| Trait | Method | $R^2$ in cohort (%) | | | | | | |
| --- | --- | --- | --- | --- | --- | --- | --- | --- |
|  |  | GS | LBC1936 | ARIES |  |  |  |  |
|  |  |  |  | A.0 | A.7 | A.15 | A.FOF | A.FOM. |
| BMI | LASSO | - | 12.2 | 1.07 | 0.13 | 2.95 | 9.37 | 14.95 |
| $\log(kg/m^2)$ | Bayes | - | 15.7 | 3.56 | 0.81 | 6.27 | 11.65 | 16.26 |
| Smoking | LASSO | - | 42.18 | - | - | - | 24.71 | - |
| $\log(p.p.y.^a + 1)$ | Bayes | - | 46.49 | - | - | - | 34.66 | - |

<sup>a</sup>packs per year

## 4 Discussion

We present BayesRR, a statistical model for joint inference of genetic and epigenetic effects over complex phenotypic traits. Using simulation, we show that BayesRR outperforms other approaches as it has the advantage of controlling for all factors at once and performing statistical inference jointly on all of the model parameters and adjusting estimates conditionally on each other. For both BMI and cigarette consumption, a large amount of phenotypic variance is captured by epigenetic markers in the training data set, which may be expected as trait-associated DNA methylation probe variation is likely to a large degree to be a consequence of the phenotype, as evidenced in our enrichment analyses and the prediction results from the ARIES study. These consequential effects lead to the expectation that ‘omics’ data may capture large amount of phenotypic variance. Thus, if applied to common complex disease, the model we present may enable accurate characterization of disease progression and better identification of individuals who are on a "path" to disease where future diagnosis is likely (i.e. those that are pre-diabetic, in the early stages of dementia, etc.). It remains to be seen whether such large amounts of phenotypic variance can be captured by a methylation array for common complex disease, but our prediction results shows that our approach can better describe the overall distribution of associations leading to improved phenotypic prediction.

By working in a Bayesian framework we derive a rich representation of the estimated effects through probability distributions, where all markers are taken into account and for which we can assess genome-wide enrichment of relevant biological features. From these distributions, we conclude that from the same set of probes in the same individuals, two example phenotypes show different architecture in the distribution of their effects, with the distribution of effects for cigarette consumption being more concentrated in a few epigenetic markers (5 markers with 95% IP explaining approx. 37%), while for BMI we have more probes associated with the phenotype (10 markers with 95% IP explain only 17%). We find novel associations as compared to previous approaches, with smoking behaviour associated with a myeloid cell nuclear differentiation antigen gene previously implicated as a biomarker for inflammation and non-Hodgkin lymphoma risk. Our genome-wide enrichment analyses, identified blood cholesterol, lipid transport and sterol metabolism pathways for BMI, and response to xenobiotic stimulus and negative regulation of RNA polymerase II promoter transcription for smoking, all with >95% posterior inclusion probability of having methylation probes with effects sizes >1.5 times larger than the average. Therefore, the approach we propose provides a more complete characterization of the molecular networks associated with disease.

There are a number of important considerations and caveats. It is important to punctuate that the inferred associations only relate to the present state of the biomarkers and are not intended to capture any causality between methylation status and outcome. Given the highly variable nature of ‘omics’ measures, the variation across data sets in the degree of confounding by experimental biases or unwanted biological variation that will contribute to the variation captured by probes, and the considerable changes with age that occur [27], it is highly unlikely that the phenotypic variance attributable to probes is stable across cohorts and with age. Determining biomarkers for future disease outcomes requires a different experimental design, for example longitudinal studies with a baseline, along with methodological extensions for causal inference within this framework, which our future work will focus on. Additionally, while we extend the model to ask how much additional phenotypic variance of each trait can be captured by methylation probes from whole blood above that captured by a SNP array, partitioning the phenotypic variance explained exactly may be difficult in data sets where factors are highly correlated. Furthermore, while we present a whole-genome enrichment approach, identifying novel pathways is currently limited and technological improvements are required to improve our ability to capture, define, and understand epigenetic marker variation. Finally, Bayesian inference comes at increased computational cost and requires the specification of prior distributions, for example here, that effects can be well described by a series of Gaussian distributions. A Student-t likelihood could be a path worth exploring as its inferences could be more robust to outliers [28] and additionally, although a Gaussian model may still be applied to categorical "disease or not" measurements, developing an extension to model binary response variables and explore performance in unbalanced case-control settings will likely be worthwhile.

In conclusion, our model can be applied to any kind of ‘omics’ data providing unbiased estimates of marker effects, conditional on other markers, covariates and on the data structure, without the need for specific cell-type proportion control. By operating in a Bayesian framework the uncertainties over the estimates given the data are represented explicitly, helping the researcher to interpret and draw conclusions over the architecture of the variance in the trait. We provide freely-available software with source code available to facilitate further replication and potential applications of the methodology (see URLs).

## 5 Materials and methods

### Statistical model

We assume additive probe effects 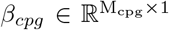, genetic effects 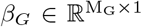, age and sex effects *α, γ* associated over a vector of measurements over a trait **y** ∈ ℝ^N*×*1^ such that,

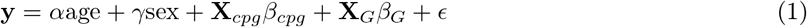

where 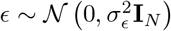, the methylation matrix **X**_*M*_ and the genotype matrix **X**_*G*_ have been centered and scaled to unit variance. We assume that only a subset of Θ = {*β_M_, β_G_, α, γ* }have an identifiable effect over trait **y**, as such, and proceeding in a Bayesian framework, we assign a sparsity inducing prior over Θ. The chosen prior follows the formulation of [29], which is a mixture of *L* Gaussian probability densities and a discrete “spike” at zero. As such, each Θ_*i*_ ∈ Θ is distributed according to:

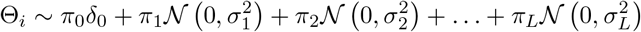

where {*π*_0_*, π*_1_*, π*_2_*, …, π_L_*} are the mixture proportions and 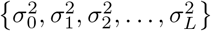 are the mixture-specific variances and *δ*_0_ is a discrete probability mass at zero.

We further constrain the prior by assuming a single parameter representing the total variance explained by the effects *σ*^2^, and the component-specific variances are proportional to *σ*^2^, that is

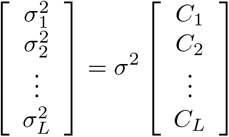

with {*C*_1_*, C*_2_*, …, C_L_*} being constants.

We expand on the previous formulation by allowing different subsets of Θ to have specific *σ*^2^ and *π* parameters. In our case, we assign to the genetic effects *β_G_* a set of mixture variances *C_G_* = {0.0001, 0.001, 0.01}, a proportion parameter *π_G_* and variance parameter 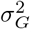. We assign to the methylation probes, age and sex effects *ϕ* = {*β_cpg_, α, γ*}the same prior variances *C_ϕ_* = {0.01, 0.1, 1} and parameters *π_ϕ_,* 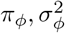

The rest of the model follows the prior hierarchy of [29] but with additional parameters for groups *G* and *ϕ*.

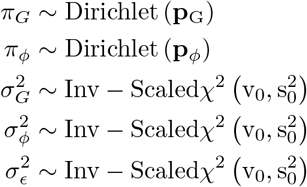

with the respective hyper parameters 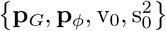 such that the prior distributions are weakly informative, **p**_*G*_ = **p**_*ϕ*_ = (1, 1, 1),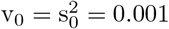

### Model Inference

Inference of the probabilistic model, follows a Gibbs sampling algorithm. Here the joint posterior probability density of parameters 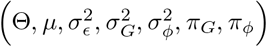 conditioned on observed phenotype **y** and observed covariates **Z** = [**X**_*G*_**X***_cpg_ age sex*] is denoted as 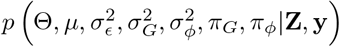 and decomposed according to the conditional distributions over each parameter:

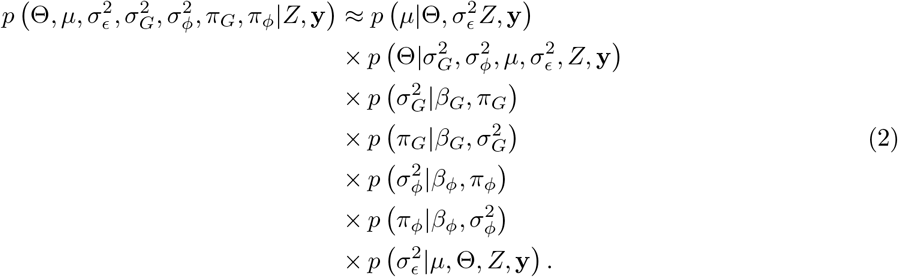

Given the prior distributions presented in eq.3, the conditional distributions are:

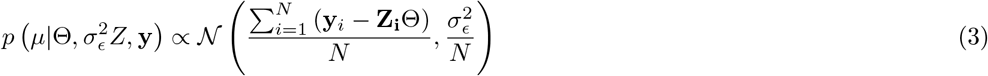

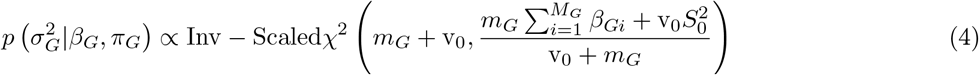

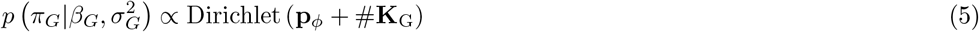

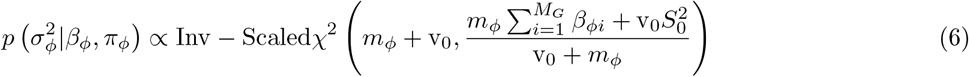

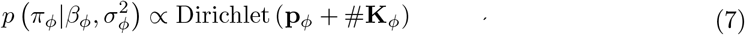

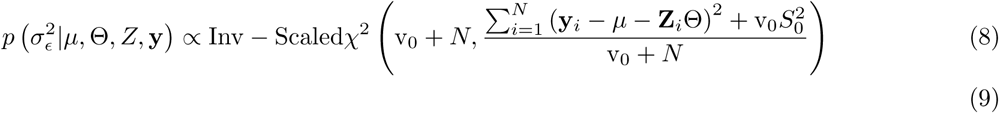

Where *m_G_* and *m_ϕ_* are the number of markers in each respective category in a sample and #**K**_*G*_,#**K**_*ϕ*_ are vectors which contain the number of markers in each mixture for the respective categories.

### Residual updating algorithm

The most computationally expensive step of sampling from the distribution in eq. 3 involves drawing from the conditional distribution

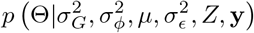

if conditioned on the Markov blanket of effects Θ, the distribution is a multivariate normal with mean **m** and co-variance Σ such that

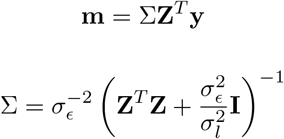

with 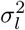 being the mixture-specific variance and the residual’s variance 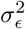. Inverting matrix Σ is of complexity *𝒪* (*M_G_* + *M_φ_*)^3^. If we use the properties of multivariate Gaussian distributions, we can decompose 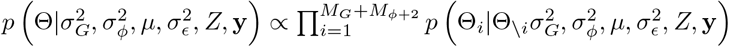, where Θ_\*i*_ represents all the effects except effect Θ_*i*_. Then each individual update consists of :

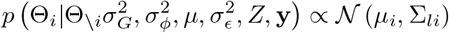

with

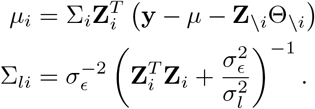

This obviates the necessity of inverting matrix Σ, if in addition we keep in memory the vector of residuals ∊ = **y** – *μ –* **Z**Θ, then we can compute efficiently **y** –*μ –* **Z**_*i*_Θ_*i*_ by the update **y** – *μ –* **Z**_*i*_Θ_*i*_ = ∊ + **Z**_*i*_Θ_*i*_ = ỹ, thus sampling from the joint distribution with a complexity 𝒪 (*M_G_* + *M_ϕ_*). Mixing and convergence issues that may arise in this formulation, have been shown to be alleviated by randomly choosing an effect Θ_*i*_ to update, as seen in successful implementations of the algorithm [29].

### Drawing from the mixtures

To select the mixture *l* from which to draw the the effect Θ_*i*_ we must evaluate the likelihood ratio between all the mixtures. Using the log-likelihood *ℒ*, this amounts to:

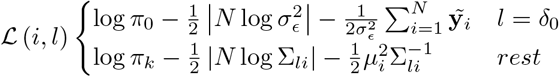

Finally, the probability of drawing effect Θ from mixture *l* is given by:

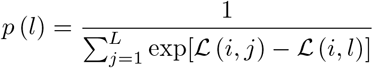

### Software implementation

The algorithm was implemented in C++-11.0, with an interface with R through Rcpp [30], and with the help of the templated matrix algebra library Eigen [31, 32]. Source code available in https://github.com/ctggroup/BayesRRcpp.

### 5.1 Simulation study of methylation data

We use a generative model of methylation levels as in [13], where the matrix **O** ∈ ℝ^N*×*M^ represents the methylation levels for M probes in N individuals. This matrix can be decomposed in a matrix of K cell proportions for N individuals, which we denote **R** ∈ ℝ^N*×*K^, and a matrix of cell-specific methylation levels **S** ∈ ℝ^K*×*M^. The decomposition assumes i.i.d. observation noise such that

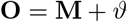

where the observation noise 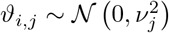, such that *I* ∈ (1 *… N*) and *j* ∈ (1 *… M*). The methylation level matrix *M* is decomposed in

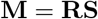

For the simulation, each row *i* of **R** is distributed according to

#### Algorithm 1

Algorithm for sampling over the posterior distribution *p* (*μ, β, ∊, σ_∊_, θ*), each sample (*μ, β, ∊, σ_∊_, θ*) is stored in a synchronised queue for a consumer thread to store in disk.. 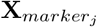 represents column of **X** corresponding to the column *j* of the vector *marker*. Given that *marker* is shuffled before sampling the effects, this is equivalent to permuting the order of the effects to be sampled.

Input: genotype matrix **X**_*G*_, methylation probe matrix **X**_*cpg*_, *age* and *sex*, vector of trait measurements **y**, prior hyperparameters 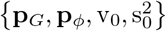, number of iterations I.

output: mean *μ*, effects vector Θ = {*β_M_, β_G_, α, γ*}, residual vector *∊*, residuals variance 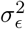 and posterior parameters, 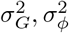

1. Initialize 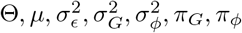
2. *effects*= (1 *… M_G_,* (*M_G_* + 1) *…* (*M_G_* + *M_ϕ_*), (*M_G_* + *M_ϕ_* + 1), (*M_G_* + *M_ϕ_* + 2))
3. set **Z** = [ **X**_*G*_ **X***_cpg_ age sex*]
4. *∊* = **y** *− μ −* **Z**Θ
5. For i in 1…I

(a) sample *μ* =
(b) *shuffle*(*effects*)
(c) For j in 1 *…* (*M_G_* + *M_ϕ_* + 2)

i. 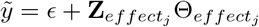
ii. Sample 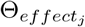
iii. 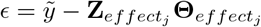
(d) sample 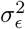
(e) sample 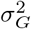
(f) sample 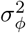
(g) *enqueue*(*μ, β, ∊, σ _∊_, θ*)

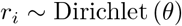

and each of the methylation levels *s_ij_* ∈ ℒ are distributed as follows for the differentially methylated regions (DMR)

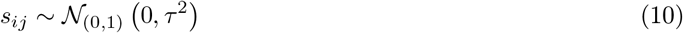

being 𝒩_(0,1)_ the truncated normal distribution with support [0, 1]. For the non-DMR we set the methylation levels to a base value

For these simulations we relied on software kindly provided by the authors [13]. We performed a slight modification to be able to change the variance of the cell proportions matrix **P**. For each simulation we generated a matrix of *N* = 2000 and *M* = 103638. The simulation parameters were in accordance to [13]:

- Proportion of differentially methylated probes (*p*).
- Variance of the DMRs(*τ*).
- Variance of the cell proportions (*sp*).
- Variance of the measurement noise (*ϑ*)

We assume 100 probes exert an effect over phenotypic trait **y**, thus for each probe selected we denote *β_d_* as the effect corresponding to a probe, and their respective effects are drawn from

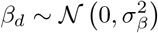

the rest of the effects are assign a value of 0. Finally we simulate **y** from the linear model **y** = **MB** + *∊* with 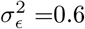 and 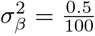 where the VE by the probes amounts to 0.6. For each of the following scenarios, 15 simulations were performed:

- *p* ∈ {0.1, 0.3, 0.7, 0.9*}*,*τ* = 0.07,*sp* = 1000,*ϑ* = 0.01
- *p* = 0.15,*τ* ∈ {0.01, 0.03, 0.05, 0.09}, *sp* = 1000,*ϑ* = 0.01
- *p* = 0.15,*τ* = 0.07,*sp* ∈ {0.001, 0.1, 1, 10, 1000},*ϑ* = 0.01
- *p* = 0.15,*τ* = 0.07,*sp* = 1000,*ϑ* ∈ {0.01, 0.025, 0.05, 0.075}, for this case the simulated model is **y** = **OB** +∊.

which amounts to 255 simulated data sets in total.

### 5.2 Inference over simulations of methylation effects

For each of the simulations, we ran our method for the observed phenotype **y** centered and scaled to variance 1, the observed methylation matrix **O** as inputs and with mixture variances (0.1, 0.01, 0.001, 0.0001), for 20000 samples with 10000 samples of burn-in after which a thinning of 10 samples was used to select samples of the posterior for the simulation. We selected trait-associated probes as those that were in the model in >95% of the posterior samples, which we define as 95% posterior inclusion probability.

We used the following metrics to assess model performance: the correlation between true effects and estimates (*ρ*(*β, β̂*)), the slope of a regression of the estimates on the true effects (*β_β̂_*_*~β*_), the number of genome-wide significant probes identified (loci), the mean square error (MSE), the MSE of the genome-wide significant probes (*MSE_sig_*), the false discovery rate (FDR), the norm of the correlation vector between a individual-level predictor made from the probe effects and the cell-type proportions (||*ρ*(**R**, **ĝ**)||), the correlation between the first principal component of the probe data and the difference between the estimated and true effect (*||ρ*(**P**, **R**)*||*) and the phenotypic variance attributable to the probes 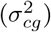. These statistics were obtained for the simulations and shown in **fig.S1**, **fig.S2** and **fig.S3**.

#### 5.2.1 Competing methods

For comparisons with our method we chose the common EWAS methodology, which is derived from the GWAS methodology. For each simulation replicate we ran the following models:

##### Single-probe least squares regression (GWAS)

We conducted a set of 103638 linear regressions using the function lm() from R version 3.4.2, and accepted the effect sizes whose p-value was less than 0.05/103638.

##### Single-probe least squares regression with sparse latent factors (ReFACTor)

Using the approach outlined in [13], we selected 5 sparse components(1 for each cell type in the simulation) and estimated probe associations conditional upon these one-by-one. We selected associated probes whose p-value was less than 0.05/103638.

##### Single-probe mixed linear model association analysis (OSCA)

Using the approach outlined in [14], we used a linear mixed effects model where probe values are used to calculate a covariance matrix, which is then used in a restricted maximum likelihood estimation (REML) approach to estimate the proportion of phenotypic variance attributable to the probes, and conditional upon this probe effects are estimated one-by-one. While this estimates probe associations conditional on the covariance matrix, it does not account for genome-wide covariance across probes. We selected associated probes whose p-value was less than 0.05/103638.

##### Multi-probe penalized regression with latent factors (LFMM)

Using the approach outlined in [17], we ran LASSO and ridge regression with 10-fold cross validation using the default settings(with 5 latent factors, one for each cell type in the simulation), whilst also fitting latent factors within the model that are intended to control for cell-type proportion confounding. We used the lfmm_function and selected probes based on the calibrated p-value whose p-value was less than 0.05/103638.

##### Multi-probe penalized regression without latent factors (glmnet)

We ran LASSO and ridge regression with 10-fold cross validation using the default settings of package glmnet [18] version 2.0-16. Previous experiments suggested that best performance was achieved by leaving the phenotype vector **y** un-scaled.

### 5.3 Simulations of genotype and methylation effects

We simulated a methylation matrix **M** with parameters *p* = 0.15,*τ* = 0.07,*sp* = 1000,*ϑ* = 0.01 as above. Then we generated a genotype matrix **X** ∈ ℝ^1000*×*103638^. For each of the DMRS with non-zero effects in **M** we select column *j* of *X* to have a correlated genotype by sampling its elements from the distribution:

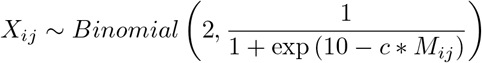

by sampling c from an uniform distribution between 20 and 25 for each column, we achieve a correlation between 0 and 0.6 among genotype and methylation levels.

Having both matrices **O** (the noisy observations over matrix **M**) and **X** centered and scaled, we generated 100 methylation effects 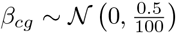 100 larger genotypic effects 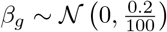 and 900 small genotypic effects 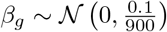. Thus the model for the simulated genotype is **y** = **O***β_cg_* + X*β_g_* +∊, with ∊ ~ 𝒩 (0.02) We repeated the process 15 times to have the same number of simulated data sets.

### 5.4 Inference over simulated genotype and methylation effects

For each of the simulated data sets we ran our method for 20000 samples, with a burnin of 10000 and a thinning of 10 samples. The mixture variances were set to (0.1, 0.01, 0.001, 0.0001) for methylation effects and to (0.01, 0.001, 0.0001, 0.00001) for genotype effects. We compared out approach to LASSO and Ridge regression implemented in glmnet [18], with a baseline of single marker regression (GWAS) where we first adjusted the phenotype by the first ten principal components of the genotype matrix and then regressed the residuals against the scaled methylation matrix. The methods were compared over the estimation of the true genetic and epigenetic VE and power to estimate the true effects. Results shown in **fig.S4**.

### 5.5 Data

#### 5.5.1 Generation Scotland

Generation Scotland: the Scottish Family Health Study is a large population-based, family-structured cohort of over 24,000 individuals aged 18-99 years. The study baseline took place between 2006-2011 and included detailed cognitive, physical, and health questionnaires, along with sample donation for genetic and biomarker data. DNA methylation data from whole blood was obtained on a subset of 5,200 participants. The Illumina HumanMethylationEPIC Bead Chips array was used to measure methylation and quality control details have been reported previously [27]. Briefly, outliers based on the visual inspection of methylated to unmethylated log intensities were excluded, along with poorly performing probes and samples, and sex mismatches (predicted based on genetics versus questionnaire data) yielding an analysis dataset of 5,101. As reported in McCartney et al. [27], further filtering was performed to exclude non-autosomal CpG sites and sites that were exclusive to the EPIC array. This allowed for the predictors to be applied to datasets that collected DNA methylation using an earlier version of the Illumina arrays (450k array) giving a total of 370,262 probes.

After the quality control steps described above, we integrated the SNP marker and methylation matrices, along with the log-transformed age and the sex of the individuals (encoded as 1 for Female). All matrices and phenotypes were centered and scaled to variance 1. The data was used as input for our Bayesian model, with parameters (0.0001, 0.001, 0.01) for genetic effects mixtures variances and (0.01, 0.1, 1) for the epigenetic effects, age and sex. Four chains for each trait with different starting values were executed. We assessed the convergence of the hyperparameters 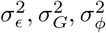 through the Geweke test [33] and the *R̂* criteria [34], with the help of the R package "ggmcmc" [35]. As result, the algorithm yielded a set of samples over the posterior distribution of effects conditioned on the observed phenotype, the genetic and epigenetic probes and controlled by age and sex. We further scaled in each sample the hyperparameters 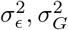 and 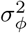 by dividing each one by their sum 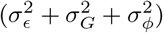. The posterior distribution is summarized in **table S1** for BMI and **table S2** for smoking. Posterior Inclusion Probabilities (IP) were computed by counting the times a probe is present in the model (in any of the mixtures) and divided by the total number of posterior samples.

#### 5.5.2 Enrichment analysis

Probes were associated to their respective gene ENTREZ identifiers using the R packages "IlluminaHumanMethylation450kanno.ilmn12.hg19" [36] and "biomaRT" [37]. We provide a list of each gene with IP greater than >5% for BMI **table S4**, and for smoking **table S5**.

Then we associated the mapped genes with their respective terms in the Gene Onthology (GO) using the R package [38]. With these probe-terms associations we computed enrichment as defined by:

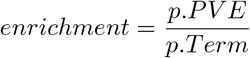

with *p.PV E* being the proportion of variance explained by probes associated with the term, having *β_Term_* being the effects associated with the term and being *β_Model_* the effects in the model(that is, those which are not coming from the "spike" at zero) in the current sample, we have

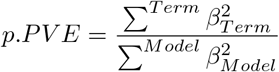

and *p.Term* being the proportion of probes mapping to the term among all the probes mapping to a term in the current sample. Having #*probes_Term_* being the number of probes mapping to a term and #*probes_Model_* the number of probes in the model in the current sample, we have

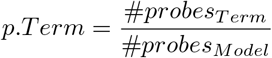

We also computed the posterior inclusion probability (IP) for a term by counting the times a term appears in the model and dividing by the number of samples. Finally, given that we have a posterior distribution over enrichment values, we adopt the ROPE decision rule [21], for which we accept the hypothesis that a term is significantly over/under-enriched if 95% of the posterior mass for the enrichment value is outside the interval (0.5,1.5) and the term has an IP >95%. Significantly enriched GO terms are presented in **fig.3e** and **fig.S15** for BMI; in **fig.3f** and **fig.S16** for smoking.

We then repeated the entire enrichment process, mapping probes to genes in the top 99% and lowest 1% percentiles of gene expression in the GTEx consortium [23] and DEPICT [24] datasets. We show the enrichment results in **fig.S17** for BMI, with the corresponding tables, **table S8**,**table S9** for GTEx; with **table S12**,**table S13** for DEPICT. For smoking the corresponding plots are shown in **fig.S18** and summarized in tables **table S10**,**table S11** for GTEx and **table S14**,**table S15** for DEPICT.

#### 5.5.3 Estimates for replication

For both BMI and smoking, the posterior samples over effects where averaged and associated to their respective probe and SNP identities. For each replication cohort a predictor was built by multiplying the posterior mean effects by the corresponding centered and scaled genetic and epigenetic marker readings, and predictive ability measured over the scaled and centered cohort trait was measured using the *R*^2^ statistic.

#### 5.5.4 Lothian Birth Cohort 1936

The Lothian Birth Cohort 1936 is a longitudinal study of aging [39]. It follows 1,091 members of the 1947 Scottish Mental Survey, who were recontacted in later life, when they were living in the Edinburgh area of Scotland. The cohort members were all born in 1936 and have been assessed for a wide variety of health and lifestyle outcomes at ages 70, 73, 76, 79, and 82 years. DNA has been collected at each clinical visit. In the present study, we considered DNA methylation data (Illumina 450k array) from whole blood, taken at mean age 70, for analysis. Details of the collection and processing of the data have been reported previously [27]. Briefly, after quality control to remove poorly performing methylation sites, samples, and individuals with mismatching genotypes or predicted sex, a sample of 906 individuals was available for prediction analysis. The genotype and methylation matrices were processed as with GS, given that the posterior effect sizes for age and sex were equal to zero for both traits, they were not included.

#### 5.5.5 Avon Longitudinal Study of Parents and Children

Samples were drawn from the Avon Longitudinal Study of Parents and Children [40], [41]. Blood from 1018 mother–child pairs (children at three time points and their mothers at two time points) were selected for analysis as part of the Accessible Resource for Integrative Epigenomic Studies (ARIES, http://www.ariesepigenomics.org.uk/ [42]. Following DNA extraction, samples were bisulphite converted using the Zymo EZ DNA MethylationTM kit (Zymo, Irvine, CA, USA). Following conversion, genome-wide methylation was measured using the Illumina Infinium HumanMethylation450 (HM450) BeadChip. The arrays were scanned using an Illumina iScan, with initial quality review using GenomeStudio. ARIES was preprocessed and normalized using the meffil R package [43]. ARIES consists of 5469 DNA methylation profiles obtained from 1022 mother-child pairs measured at five time points (three time points for children: birth, childhood and adolescence; and two for mothers: during pregnancy and at middle age). Low quality profiles were removed from further processing, and the remaining 4593 profiles were normalized using the Functional Normalization algorithm [44] with the top 10 control probe principal components. Full details of the preprocessing and normalization of ARIES has been described previously [43].

## 6 Acknowledgements

We thank our colleagues at the Complex Trait Genetics Group at the University of Lausanne for their comments. The Lausanne group is funded by a Swiss National Science Foundation project grant to MRR (31003A-179380), and by core funding from the University of Lausanne. We thank Eran Halperin and Elior Rahmani for kindly providing simulation software. We would also like to thank the participants of the cohort studies. Generation Scotland received core support from the Chief Scientist Office of the Scottish Government Health Directorates [CZD/16/6] and the Scottish Funding Council [HR03006]. Genotyping and DNA methylation profiling of the GS:SFHS samples was carried out by the Genetics Core Laboratory at the Wellcome Trust Clinical Research Facility, Edinburgh, Scotland and was funded by the Medical Research Council UK and the Wellcome Trust (Wellcome Trust Strategic Award “STratifying Resilience and Depression Longitudinally” ((STRADL) Reference 104036/Z/14/Z)). The Lothian Birth Cohort 1936 is supported by Age UK (Disconnected Mind programme) and the Medical Research Council (MR/M01311/1). Methylation typing was supported by Centre for Cognitive Ageing and Cognitive Epidemiology (Pilot Fund award), Age UK, The Wellcome Trust Institutional Strategic Support Fund, The University of Edinburgh, and The University of Queensland. This work was part-conducted in the Centre for Cognitive Ageing and Cognitive Epidemiology, which is supported by the Medical Research Council and Biotechnology and Biological Sciences Research Council (MR/K026992/1), and which supports IJD. DLM and REM are supported by Alzheimer’s Research UK major project grant ARUK-PG2017B-10. Methylation data in the ALSPAC cohort were generated as part of the UK BBSRC funded (BB/I025751/1 and BB/I025263/1) Accessible Resource for Integrated Epigenomic Studies (ARIES, http://www.ariesepigenomics.org.uk).

## 7 URLS, Code and Data Availability

BayesRR implementation and full open source code is available at: https://github.com/ctggroup/bayesRRcpp.

Simulation scripts and post-processing scripts ican be found here: https://github.com/ctggroup/BEpigenetics.

Data are available upon request from the cohort authors with appropriate research agreements.

## 8 Author contributions

MRR, REM and DTB conceived and designed the experiments. DTB conducted the experiments, with oversight by MRR and RM, and assistance of DLM, TB and GH. DTB and MRR derived the equations and the algorithm, developed the software, and wrote the paper. RMW, QZ, DJP, CSH, AFM, NRW, PMV, AMM, KLE and IJD provided study oversight and contributed data to the analysis. All authors approved the final manuscript prior to submission.

## Supplementary Online Material for Bayesian reassessment of the epigenetic architecture of complex traits

**Figure S1.**
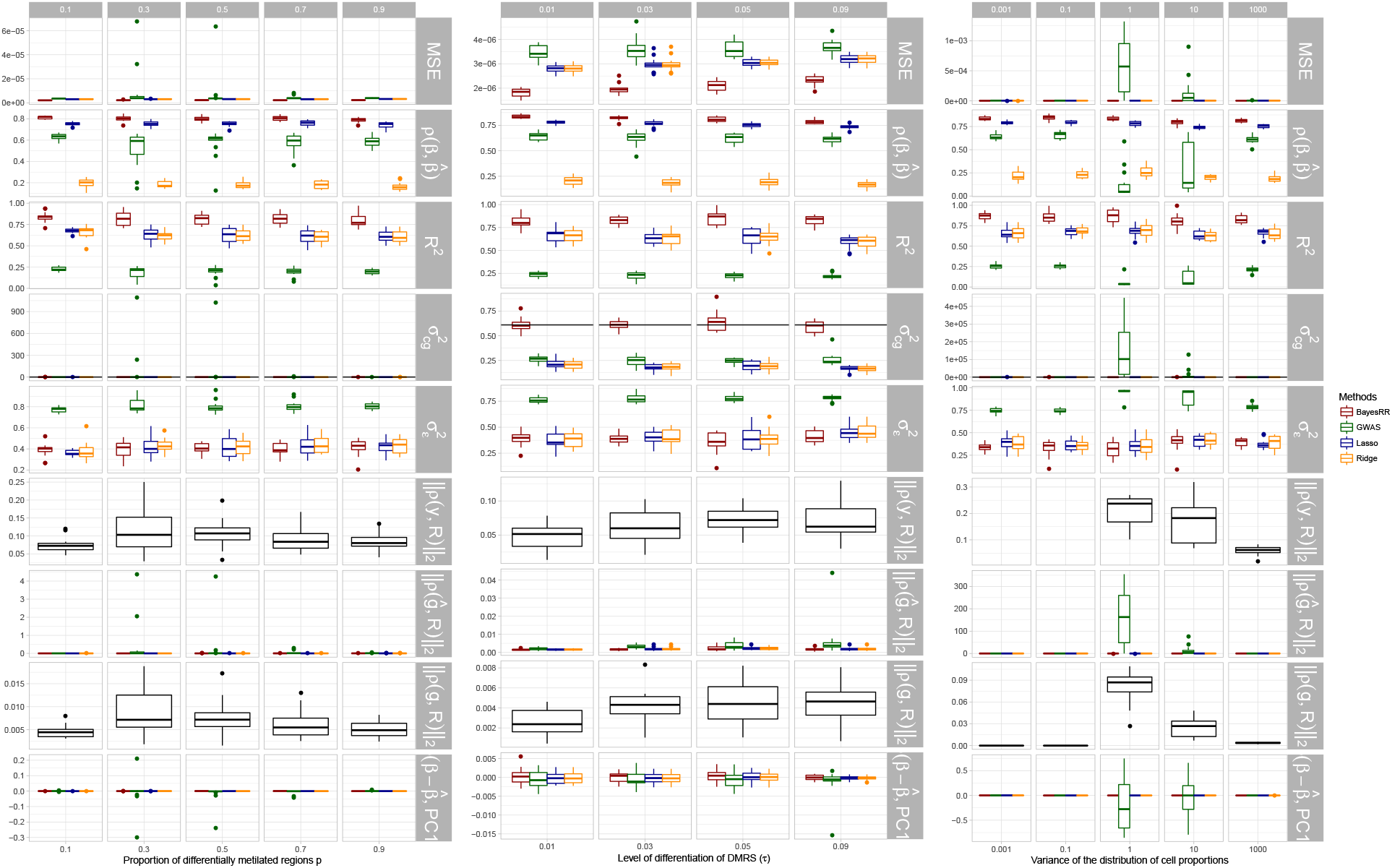
Simulation results of multi-probe approaches without latent factor correction with single-probe least squares regression included as a baseline. Estimation of phenotype-epigenetic associations comparing BayesRR to other multi-probe association methods that do not include latent factors within the model (LASSO and Ridge regression as implimented in glmnet), with a baseline of single-probe least squares regression (GWAS, see Methods), across different proportions of differentially methylated regions (DMRS), different levels of differentiation of of the DMRS, and different variances of the cell-type proportions. In many of these scenarios, probes are associated with cell-type proportion variation and the degree of cell-type proportion confounding is given by the norm of the correlation vector between the phenotype and the cell-type proportions that is shown in row panel (||*ρ*(**R**, **y**)||). The remaining panels give the mean square error (MSE), the correlation between true effects and estimates (*ρ*(*β, β̂*)), the variance attributable to the top probes (**R**^2^), the phenotypic variance attributable to the probes 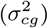, the error variance 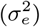, the norm of the correlation vector between a individual-level predictor made from the probe effects and the cell-type proportions (||*ρ*(**R**, **ĝ**)), the norm of the correlation vector between the true individual-level genetic value and the cell-type proportions (||*ρ*(**R**, **g**)||), and the correlation between the first principal component of the probe data and the difference between the estimated and true effect (||*ρ*(**P**, **R**)||). Across all simulation settings and degrees of cell-type confounding, BayesRR outperforms comparable multi-probe association methods and returns associations that are unbiased of cell-type effects.

**Figure S2.**
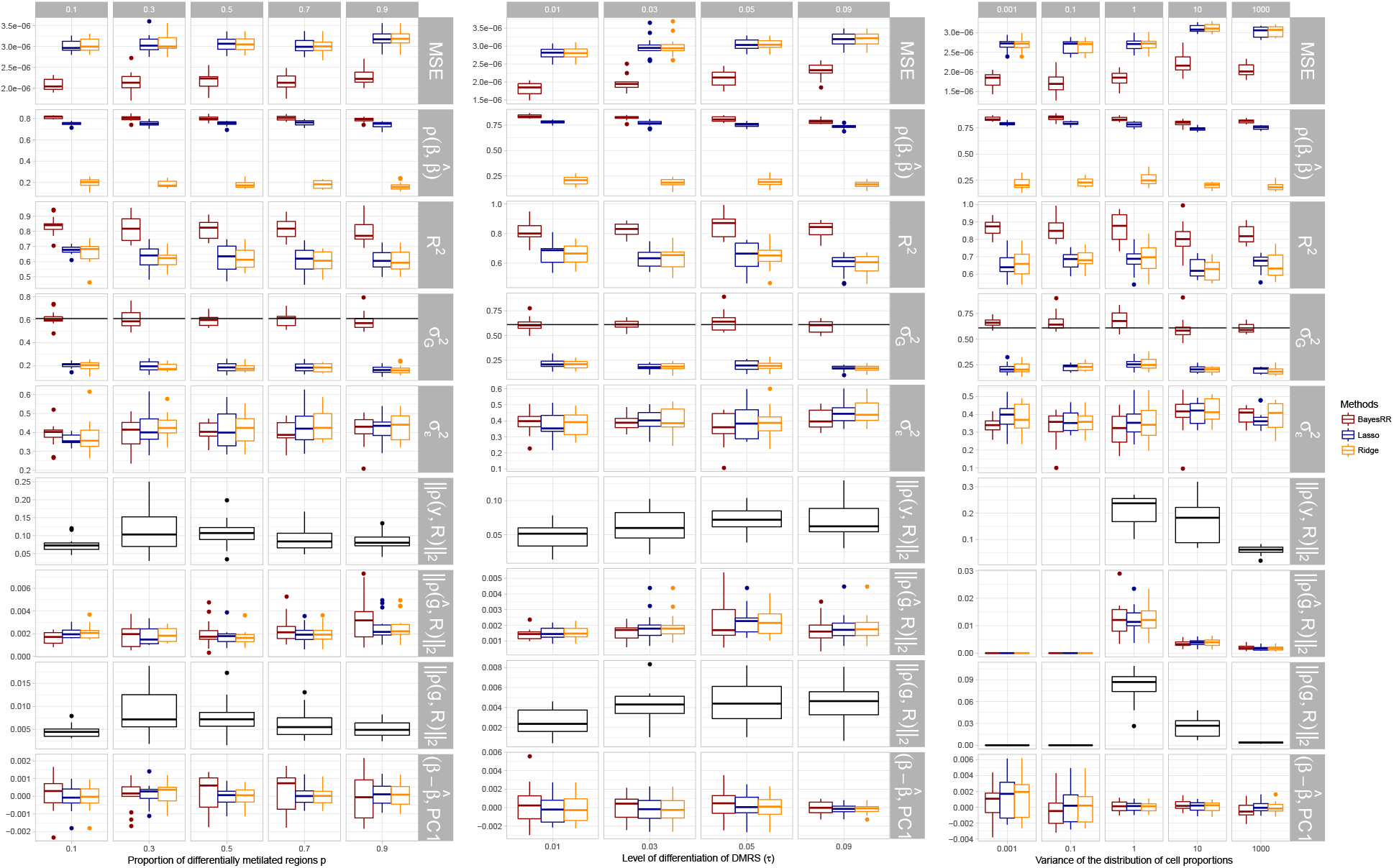
Simulation results of multi-probe approaches without latent factor correction. Estimation of phenotype-epigenetic associations comparing BayesRR to other multi-probe association methods that do not include latent factors within the model (LASSO and Ridge regression as implimented in glmnet, see Methods), across different proportions of differentially methylated regions (DMRS), different levels of differentiation of of the DMRS, and different variances of the cell-type proportions. In many of these scenarios, probes are associated with cell-type proportion variation and the degree of cell-type proportion confounding is given by the norm of the correlation vector between the phenotype and the cell-type proportions that is shown in row panel (||*ρ*(**R**, **y**)||). The remaining panels give the mean square error (MSE), the correlation between true effects and estimates (*ρ*(*β, β̂*)), the variance attributable to the top probes (**R**^2^), the phenotypic variance attributable to the probes 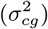, the error variance 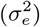, the norm of the correlation vector between a individual-level predictor made from the probe effects and the cell-type proportions (||*ρ*(**R**, **ĝ**)||), the norm of the correlation vector between the true individual-level genetic value and the cell-type proportions (||*ρ*(**R**, **g**)||), and the correlation between the first principal component of the probe data and the difference between the estimated and true effect (||*ρ*(**P**, **R**)||). Across all simulation settings and degrees of cell-type confounding, BayesRR outperforms comparable multi-probe association methods and returns associations that are unbiased of cell-type effects.

**Figure S3.**
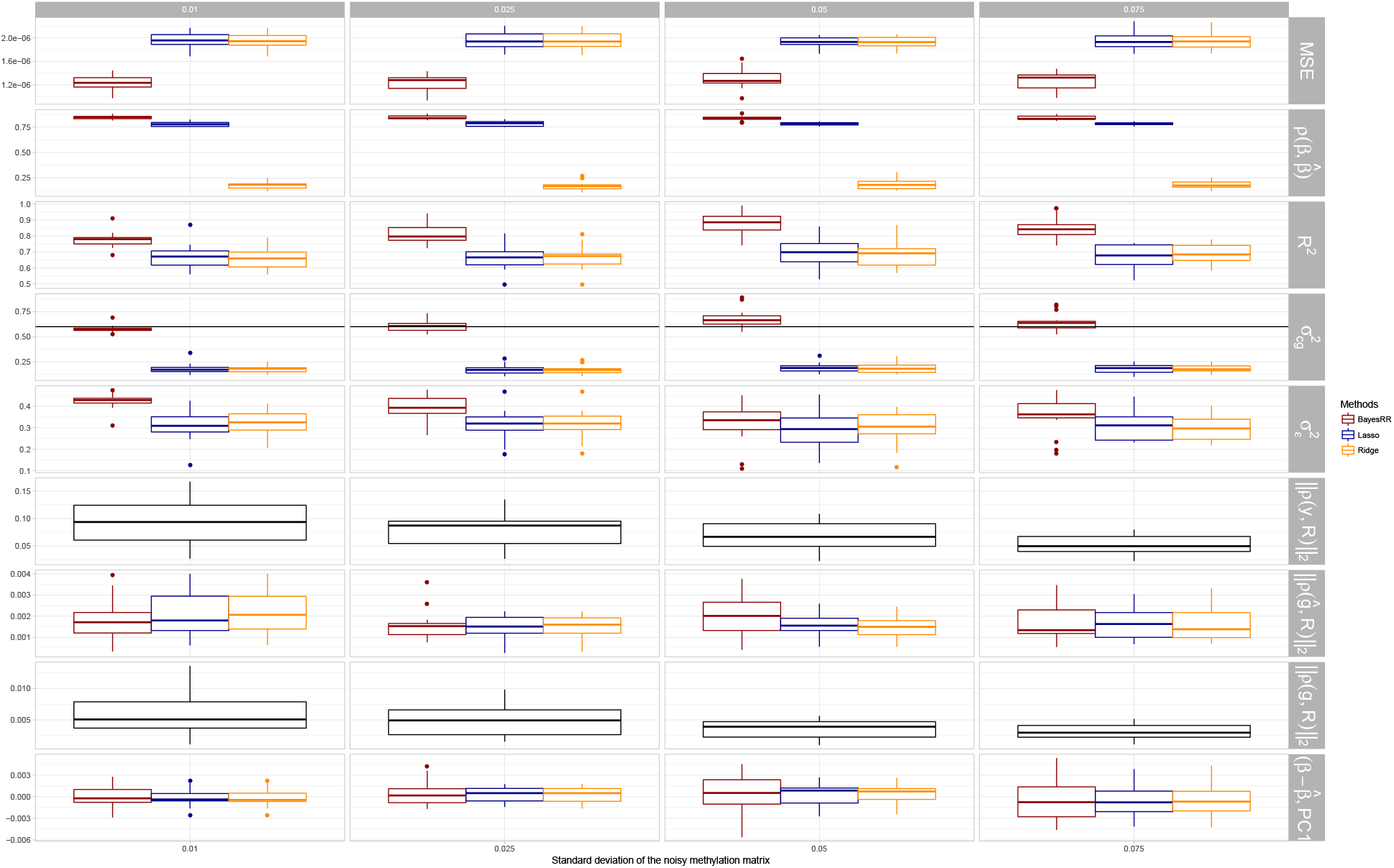
Simulation results of multi-probe approaches without latent factor correction with varying degrees of noise added to the probe data. Estimation of phenotype-epigenetic associations comparing BayesRR to other multi-probe association methods that do not include latent factors within the model (LASSO and Ridge regression as implimented in glmnet, see Methods), across different SD of random noise added to the methylation data. In many of these scenarios, probes are associated with cell-type proportion variation and the degree of cell-type proportion confounding is given by the norm of the correlation vector between the phenotype and the cell-type proportions that is shown in row panel (||*ρ*(**R**, **y**)||). The remaining panels give the mean square error (MSE), the correlation between true effects and estimates (*ρ*(*β, β*̂)), the variance attributable to the top probes (**R**^2^), the phenotypic variance attributable to the probes 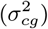, the error variance 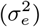, the norm of the correlation vector between a individual-level predictor made from the probe effects and the cell-type proportions (||*ρ*(**R**, **g**̂)||), the norm of the correlation vector between the true individual-level genetic value and the cell-type proportions (||*ρ*(**R**, **g**)||), and the correlation between the first principal component of the probe data and the difference between the estimated and true effect (||*ρ*(**P**, **R**)||). Even with increased degrees of experimental and measurement error, BayesRR outperforms comparable multi-probe association methods.

**Figure S4.**
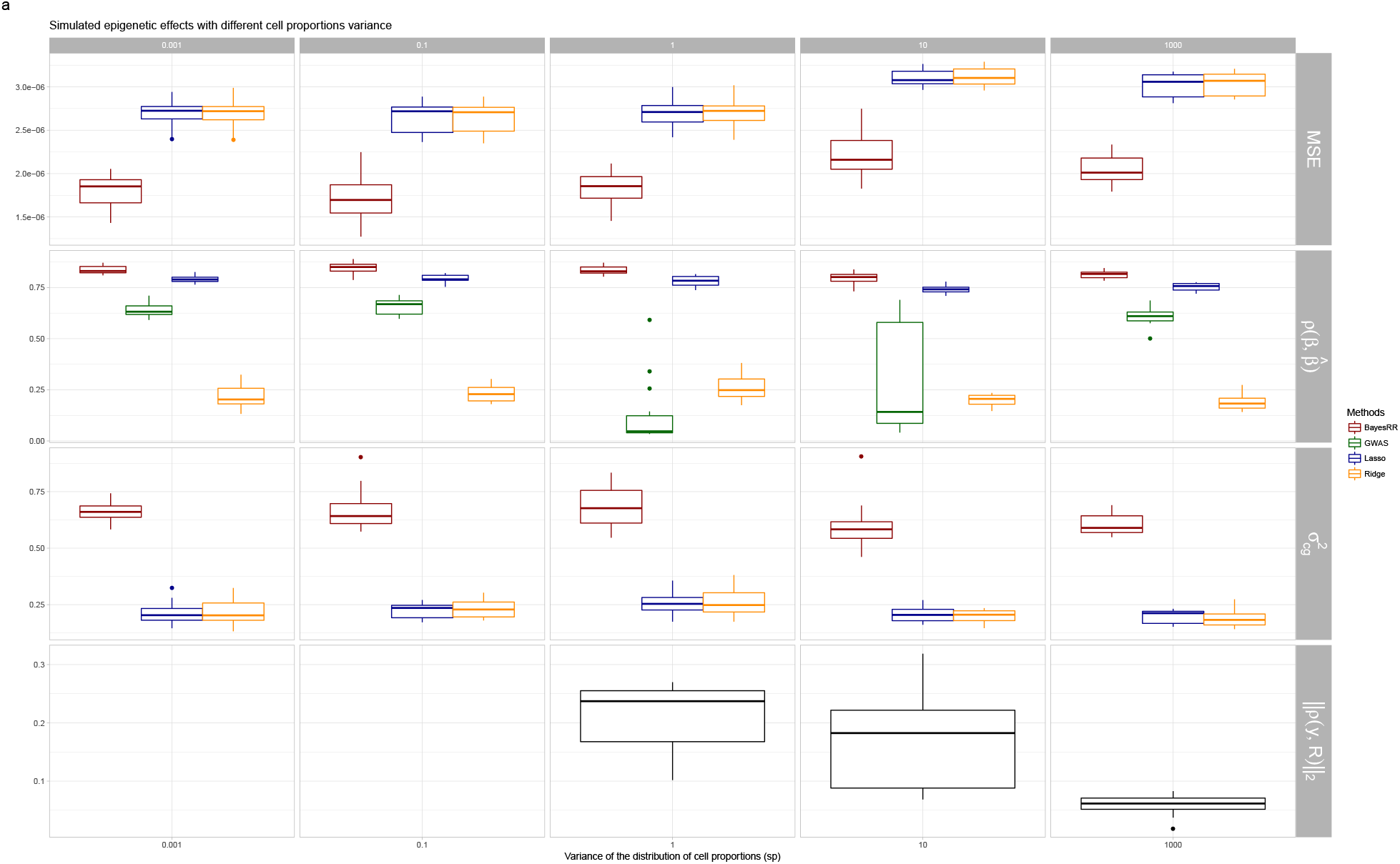
Simulation results of multi-probe approaches without latent factor correction with single-probe least squares regression included as a baseline for model that jointly estimated both SNP marker and methylation probe associations. Estimation of phenotype-epigenetic associations comparing BayesRR to other multi-probe association methods that do not include latent factors within the model (LASSO and Ridge regression as implimented in glmnet), with a baseline of single-probe least squares regression (GWAS, see Methods), across different variances of the cell-type proportions. Results are shown for the methylation probe associations only as this is the focus of this work and we wished to assess whether the performance differed when another set of correlated covariates were in the model. In many of these scenarios, probes are associated with cell-type proportion variation and the degree of cell-type proportion confounding is given by the norm of the correlation vector between the phenotype and the cell-type proportions that is shown in row panel (||*ρ*(**R**, **y**)||). The remaining panels give the mean square error (MSE), the correlation between true effects and estimates (*ρ*(*β, β*̂)), the variance attributable to the top probes (**R**^2^), the phenotypic variance attributable to the probes 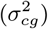, the error variance 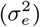, the norm of the correlation vector between a individual-level predictor made from the probe effects and the cell-type proportions (||*ρ*(**R**, **g**̂)||), the norm of the correlation vector between the true individual-level genetic value and the cell-type proportions (||*ρ*(**R**, **g**)||), and the correlation between the first principal component of the probe data and the difference between the estimated and true effect (||*ρ*(**P**, **R**)||). Across all simulation settings and degrees of cell-type confounding, BayesRR outperforms comparable multi-probe association methods and returns associations that are unbiased of cell-type effects and unbiased of the SNP marker effects.

**Figure S5.**
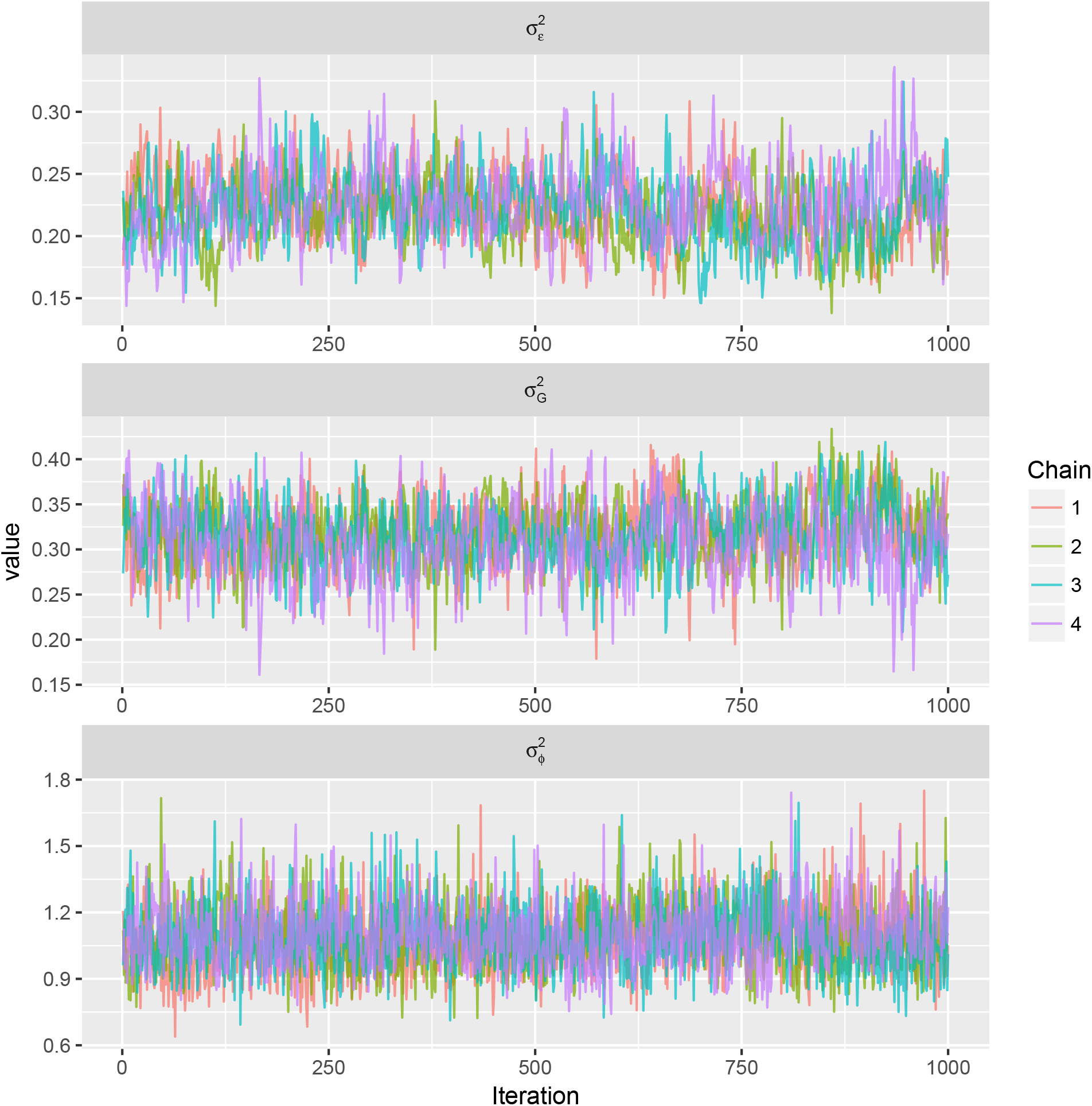
Traceplot for the hyper-parameters of the body mass index (BMI) model in the Generation Scotland dataset. Four chains for BMI were run first for 15000 samples with a burn-in of 10000 samples and thinning of 5 samples. After preliminary diagnostics it was decided to re-start the chains from the last sample, for another 15000 samples with a burn in of 10000 samples and a thinning of 10 samples. This resulted in 4 chains with 1000 samples each. Row panels give the estimates for the error variance 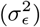, the variance attributable to SNP marker effects 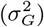, and the phenotypic variance associated with methylation probes 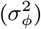.

**Figure S6.**
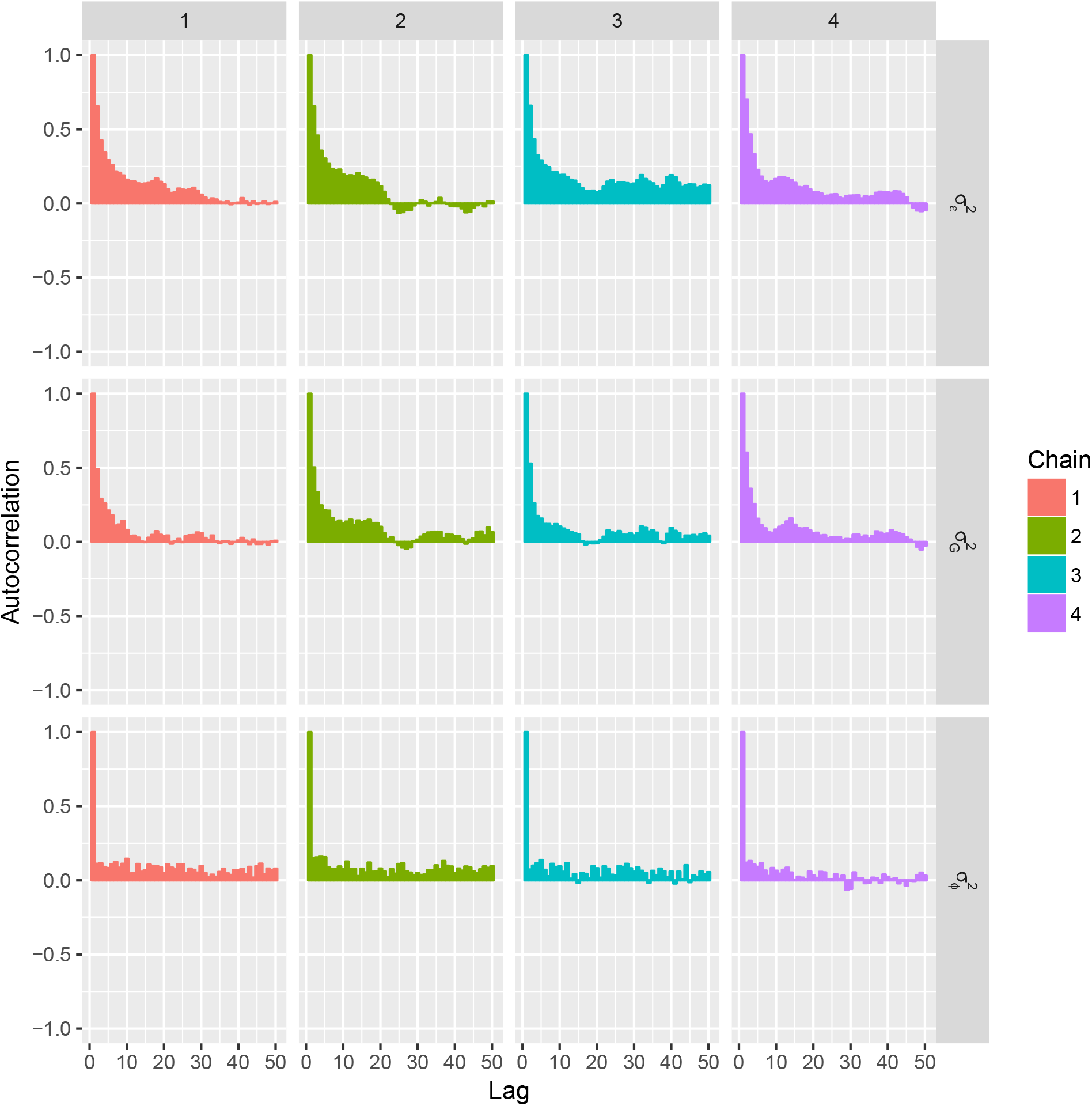
Autocorrelation plot for the hyper-parameters of the body mass index (BMI) model in the Generation Scotland dataset. Four chains for BMI were run first for 15000 samples with a burn-in of 10000 samples and thinning of 5 samples. After preliminary diagnostics it was decided to re-start the chains from the last sample, for another 15000 samples with a burn in of 10000 samples and a thinning of 10 samples. This resulted in 4 chains with 1000 samples each. Row panels give the autocorrelation (correlation of the chain with itself after a lag) for the four chains for the estimates of the error variance 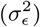, the variance attributable to SNP marker effects 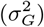, and the phenotypic variance associated with methylation probes 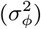.

**Figure S7.**
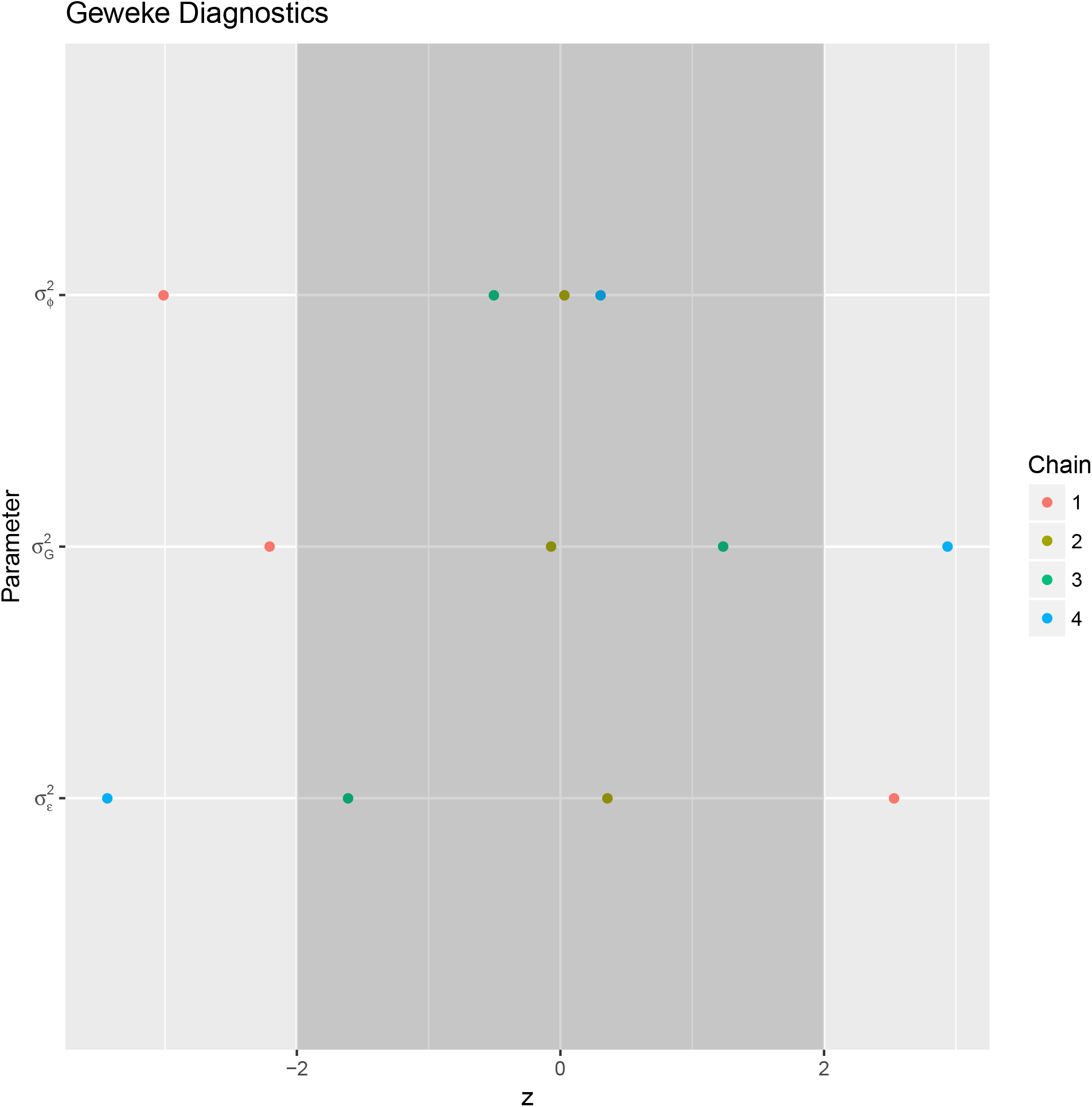
Geweke z-score diagnostic for the hyper-parameters of the body mass index (BMI) model in the Generation Scotland dataset. Results for the four chains for the estimates of the error variance 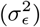, the variance attributable to SNP marker effects 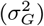, and the phenotypic variance associated with methylation probes 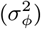, were compared using the Geweke z-score diagnostic with the acceptance interval(−2,2) highlighted in dark gray. Most chains reached the same parameter estimates, although there is some evidence of chain-level variance, meaning some effect of the starting values.

**Figure S8.**
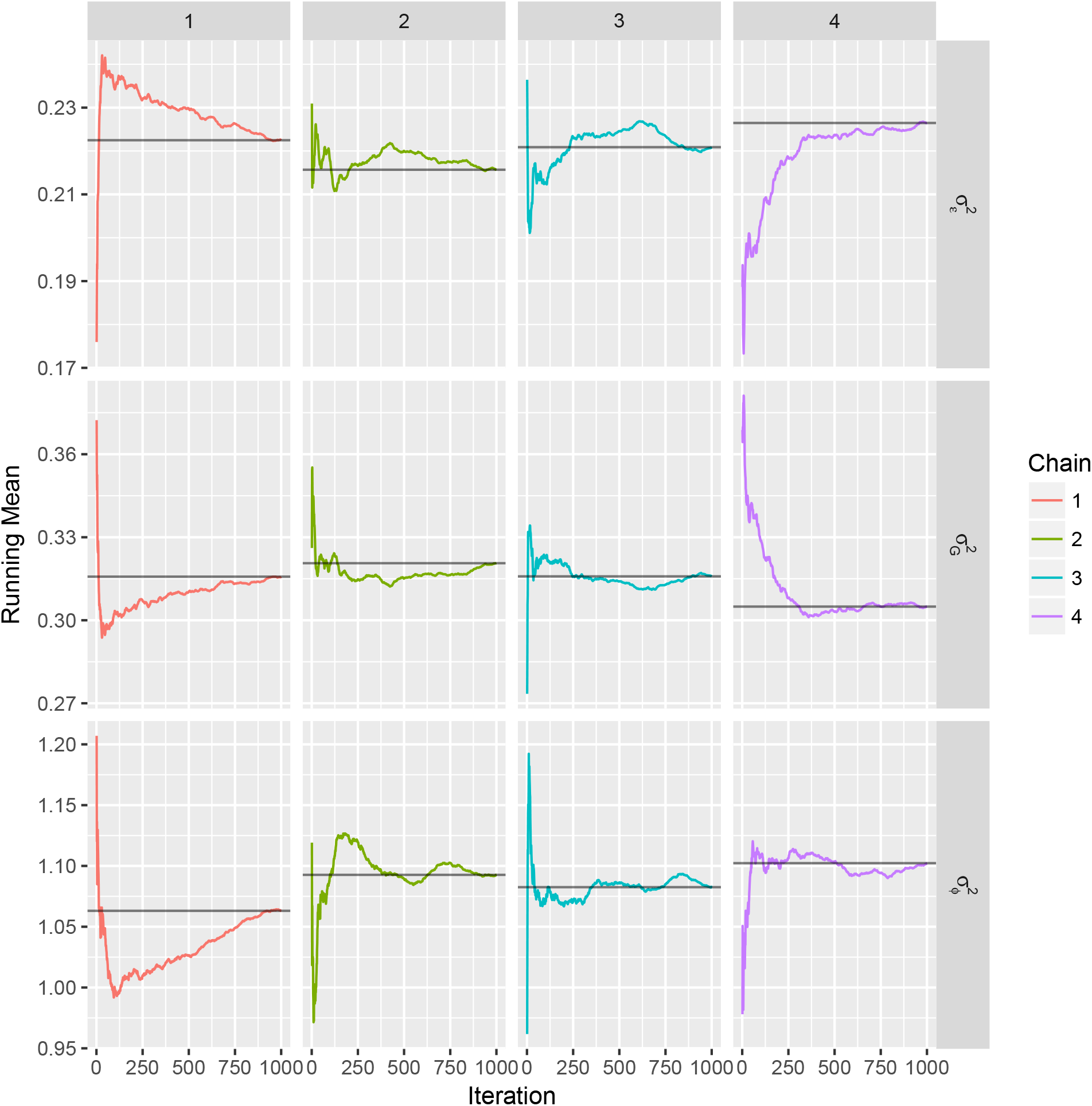
Running means plot for the hyper-parameters of the body mass index (BMI) model in the Generation Scotland dataset. The mean of the chain at each sample is compared against the chains mean(horizontal line) for the estimates of the error variance 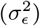, the variance attributable to SNP marker effects 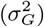, and the phenotypic variance associated with methylation probes 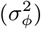. Given this, and the evidence of the among chain variance we combined the last 250 samples for each chain as samples from the posterior distribution for the subsequent analysis.

**Figure S9.**
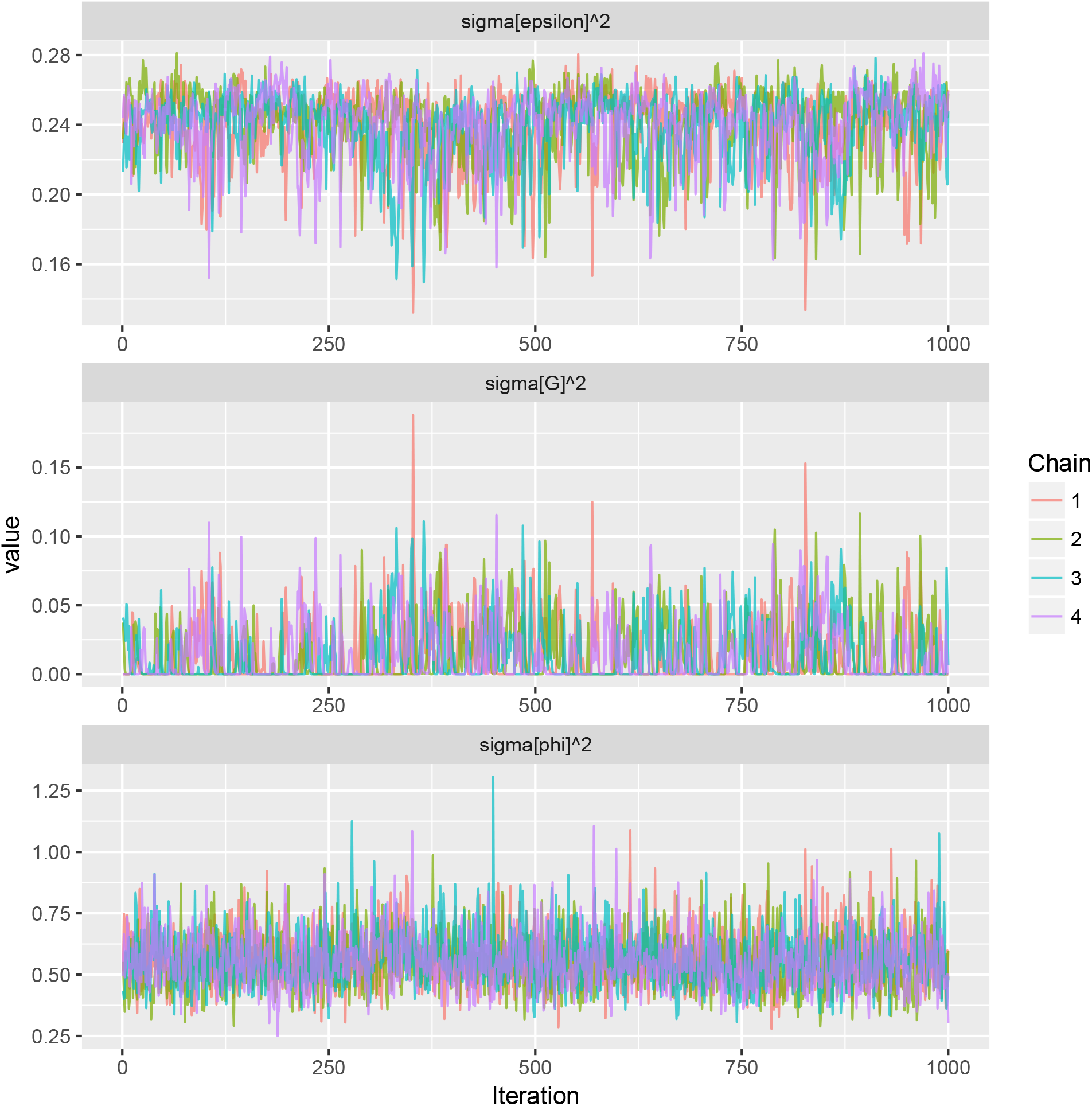
Traceplot for the hyper-parameters of the smoking model in the Generation Scotland dataset. Four chains for BMI were run first for 15000 samples with a burn-in of 10000 samples and thinning of 5 samples. After preliminary diagnostics it was decided to re-start the chains from the last sample, for another 15000 samples with a burn in of 10000 samples and a thinning of 10 samples. This resulted in 4 chains with 1000 samples each. Row panels give the estimates for the error variance 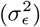, the variance attributable to SNP marker effects 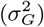, and the phenotypic variance associated with methylation probes 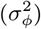.

**Figure S10.**
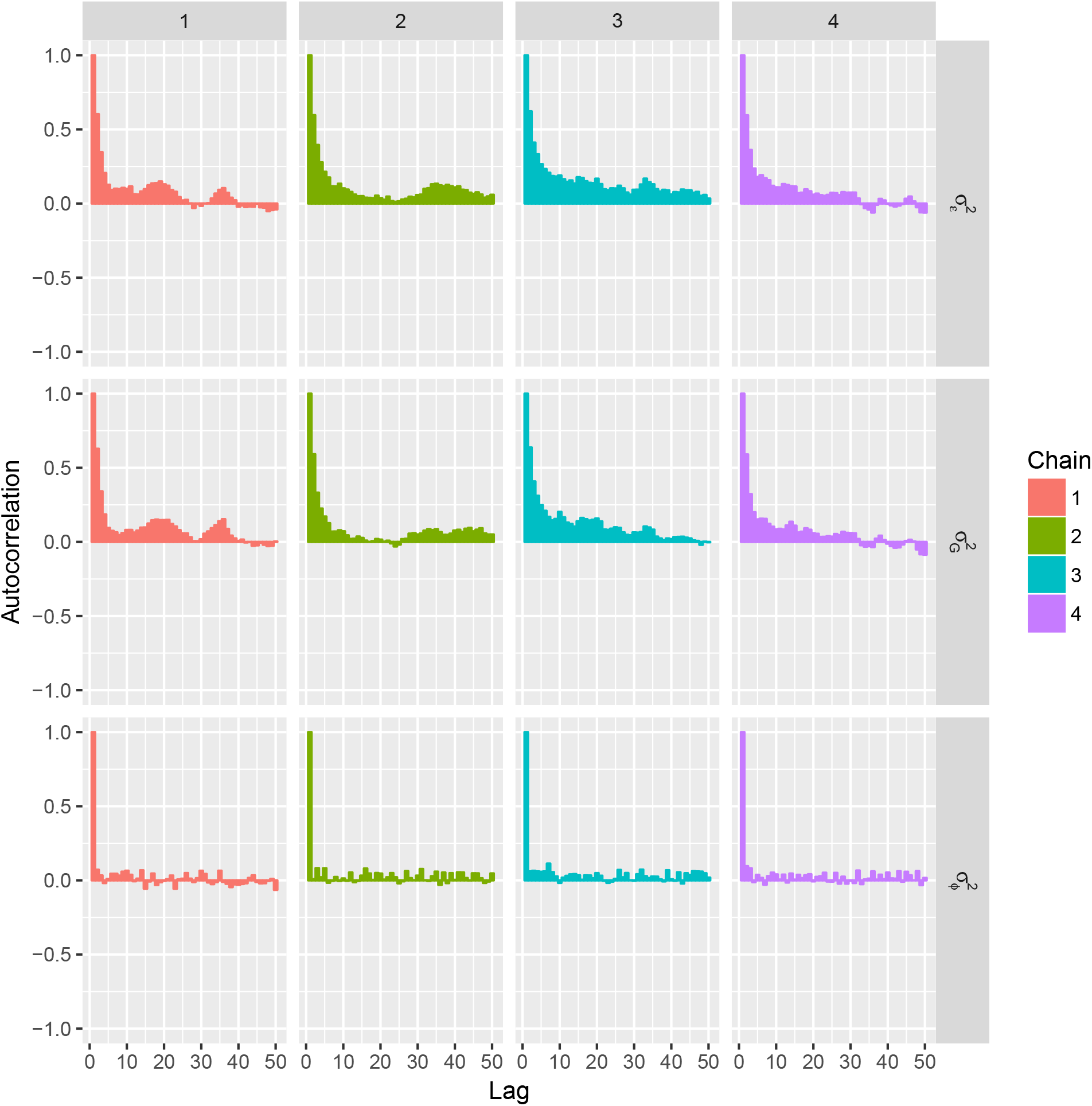
Autocorrelation plot for the hyper-parameters of the smoking model in the Generation Scotland dataset. Four chains for BMI were run first for 15000 samples with a burn-in of 10000 samples and thinning of 5 samples. After preliminary diagnostics it was decided to re-start the chains from the last sample, for another 15000 samples with a burn in of 10000 samples and a thinning of 10 samples. This resulted in 4 chains with 1000 samples each. Row panels give the autocorrelation (correlation of the chain with itself after a lag) for the four chains for the estimates of the error variance 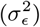, the variance attributable to SNP marker effects 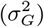, and the phenotypic variance associated with methylation probes 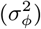.

**Figure S11.**
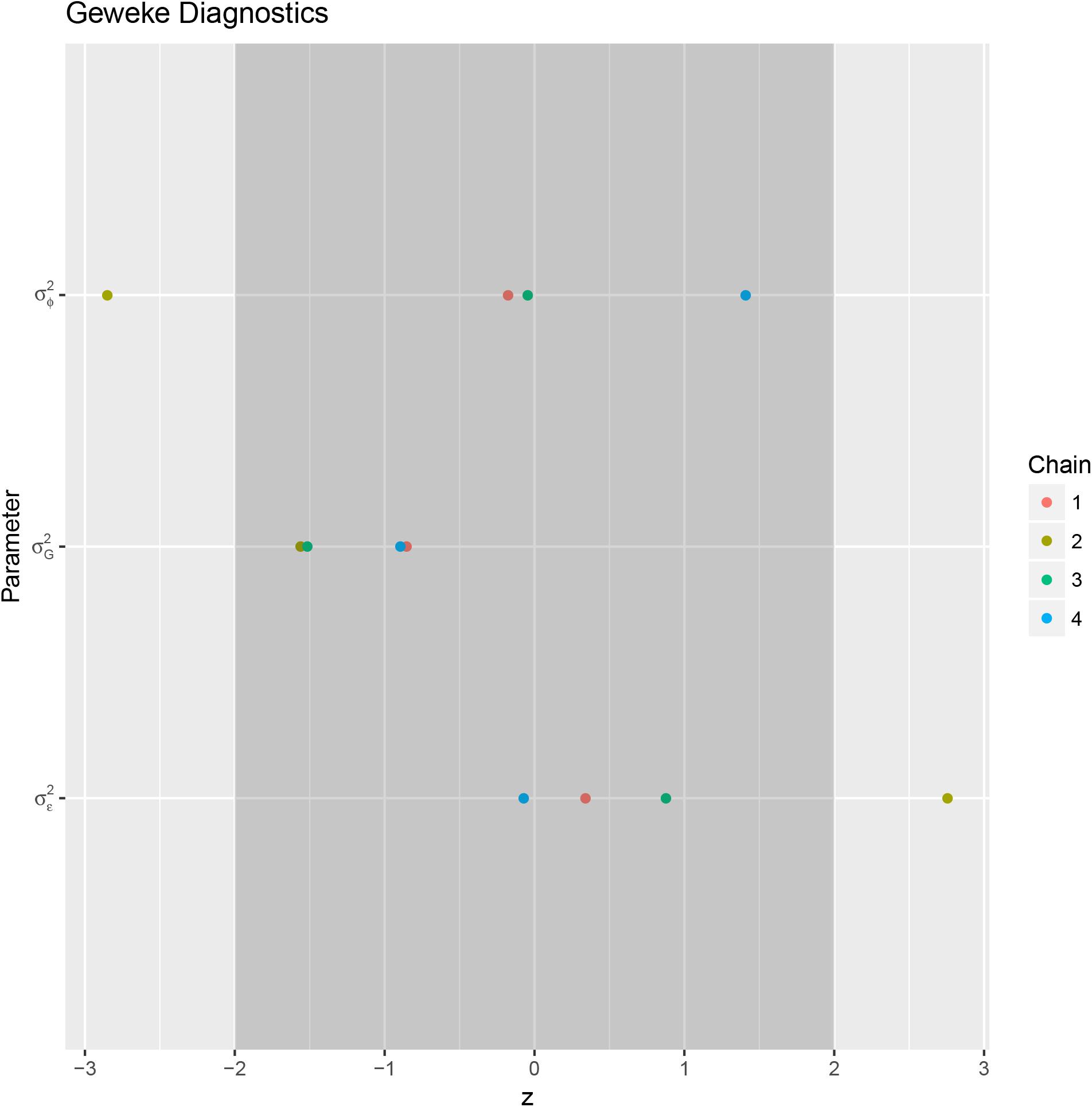
Geweke z-score diagnostic for the hyper-parameters of the body mass index (BMI) model in the Generation Scotland dataset. Results for the four chains for the estimates of the error variance 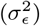, the variance attributable to SNP marker effects 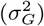, and the phenotypic variance associated with methylation probes 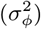, were compared using the Geweke z-score diagnostic with the acceptance interval(−2,2) highlighted in dark gray. Most chains reached the same parameter estimates, although there is some evidence of chain-level variance, meaning some effect of the starting values.

**Figure S12.**
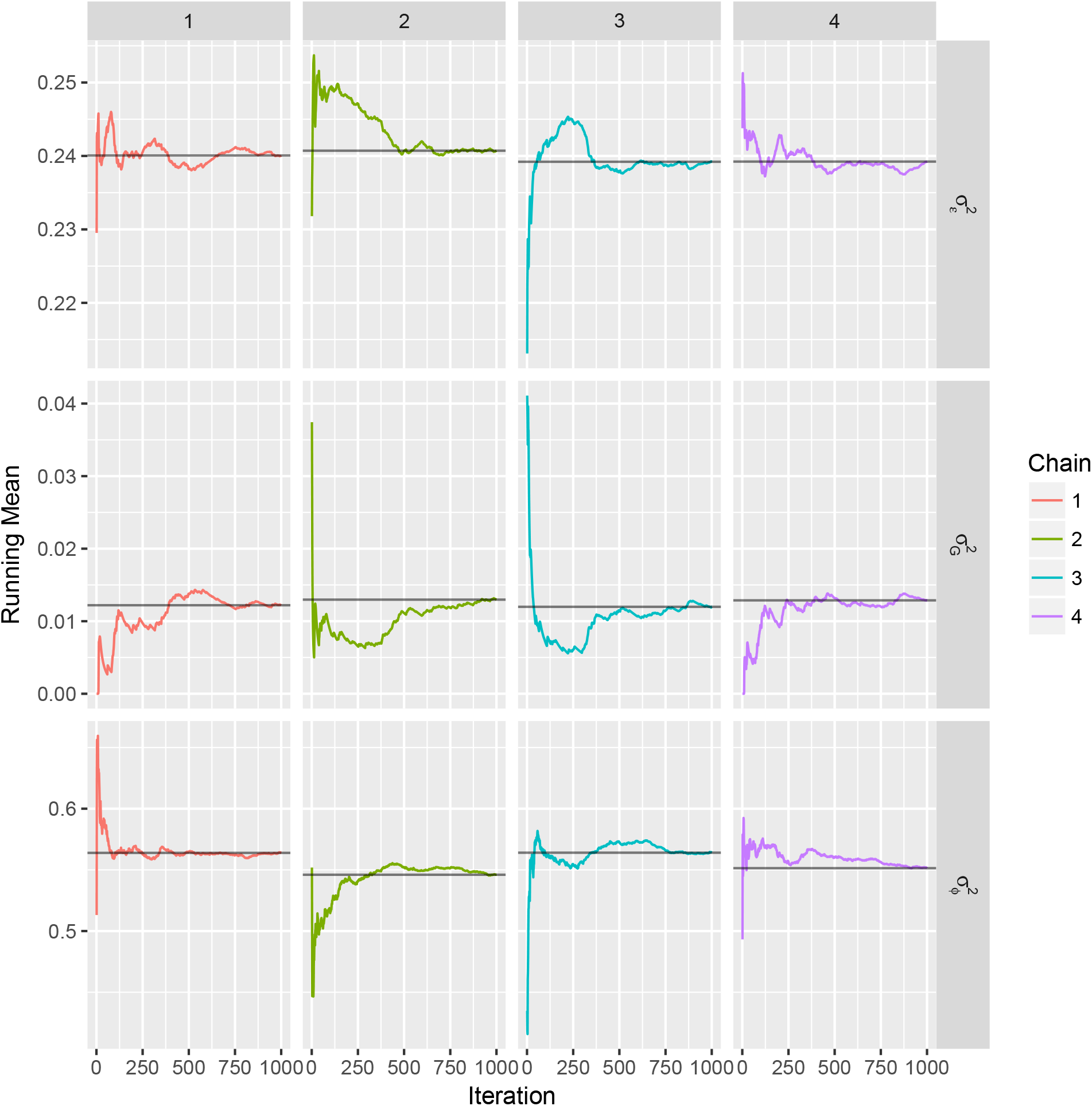
Running means plot for the hyper-parameters of the smoking model in the Generation Scotland dataset. The mean of the chain at each sample is compared against the chains mean(horizontal line) for the estimates of the error variance 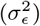, the variance attributable to SNP marker effects 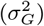, and the phenotypic variance associated with methylation probes 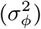. Given this, and the evidence of the among chain variance we combined the last 250 samples for each chain as samples from the posterior distribution for the subsequent analysis.

**Figure S13.**
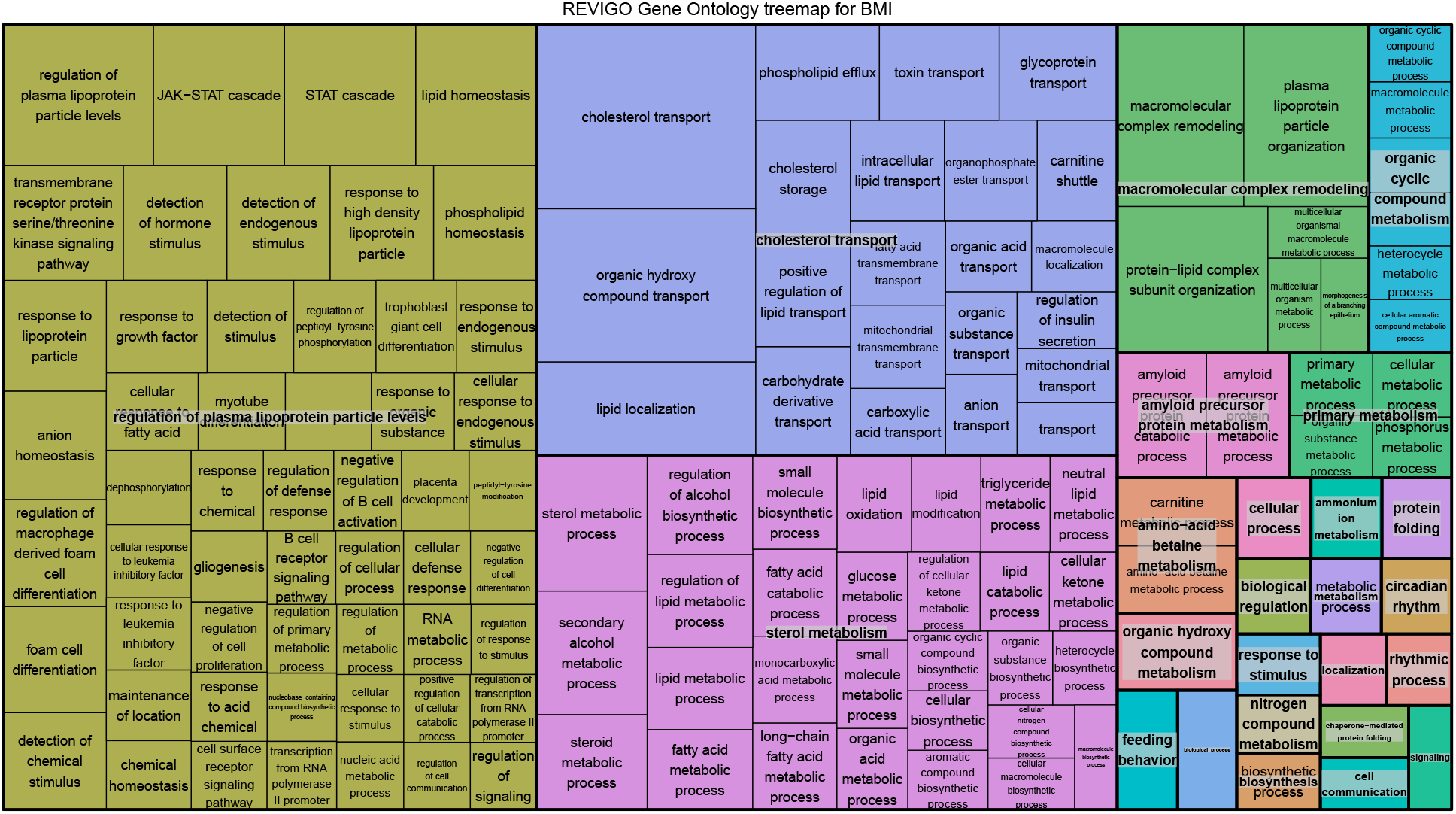
Treemap of the top GO enriched terms for BMI(enrichment>2.0 and PIP>95%). Plot generated with REVIGO

**Figure S14.**
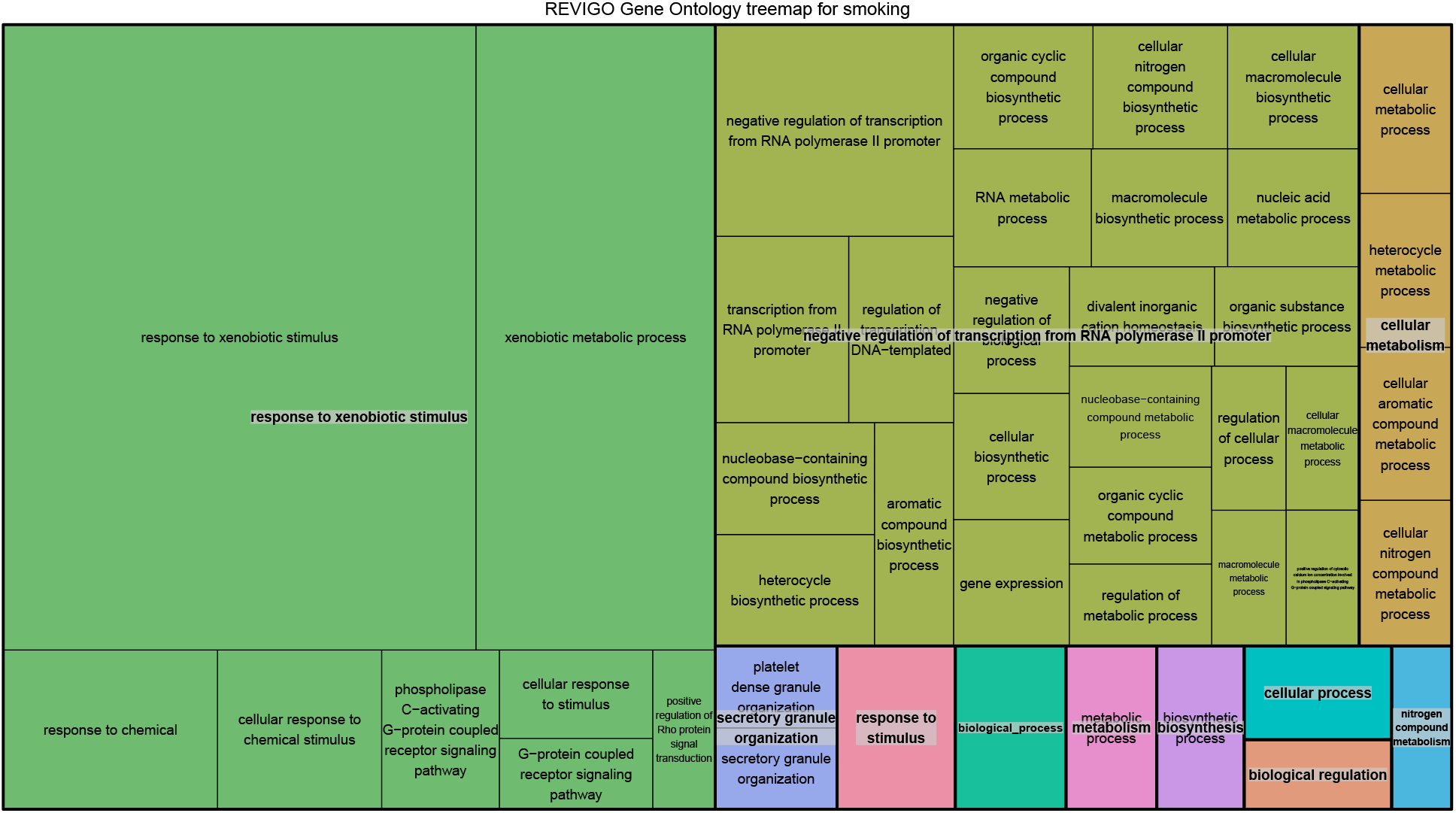
Treemap of the top GO enriched terms for smoking(enrichment>2.0 and PIP>95%). Plot generated with REVIGO

**Figure S15.**
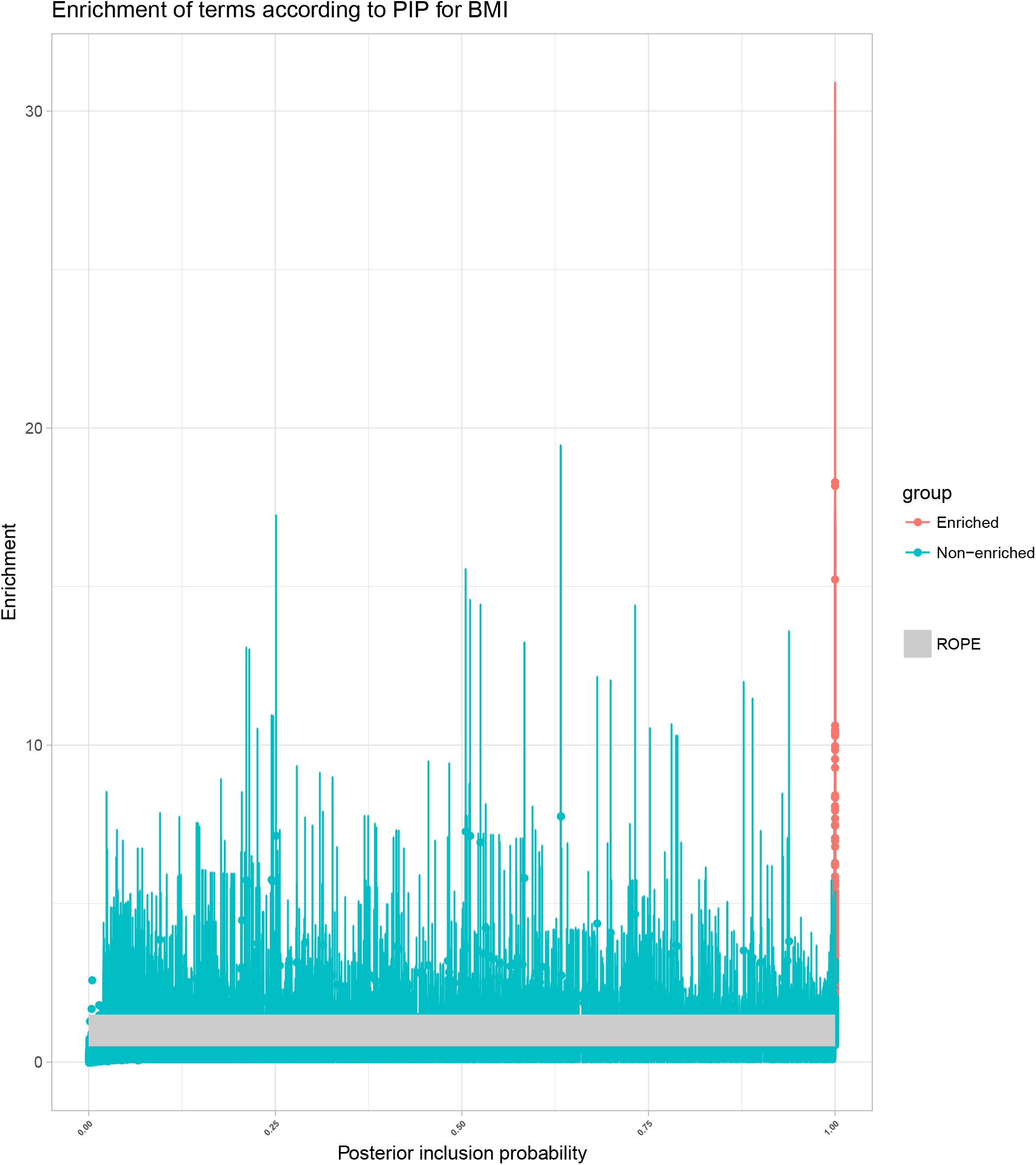
Enrichment of GO terms for BMI, plotted are the mean terms enrichment and the 95% highest posterior density intervals against the posterior inclusion probability of the term, on gray the rope area around enrichment 1.0(0.5-1.5).

**Figure S16.**
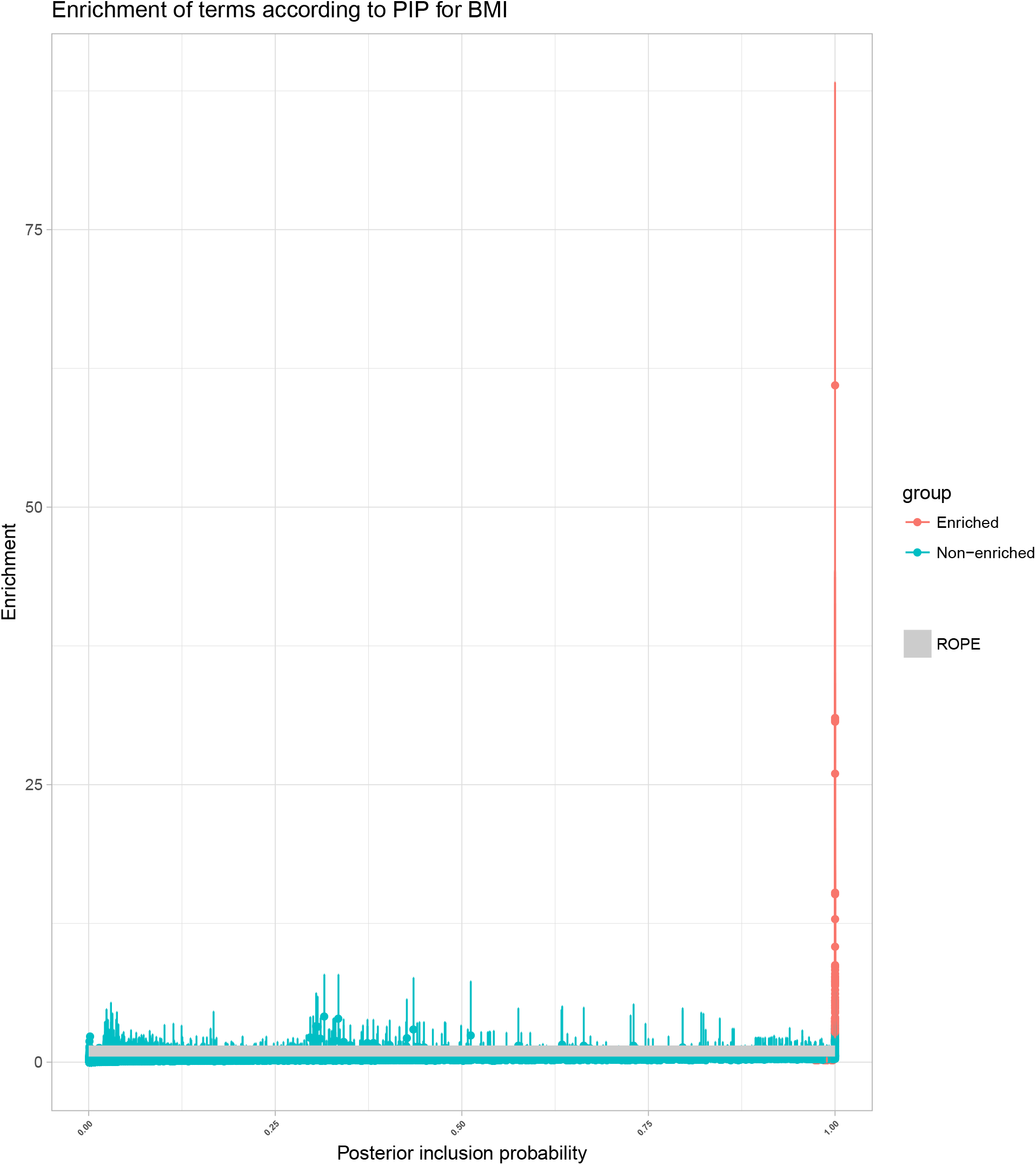
Enrichment of GO terms for smoking, plotted are the mean terms enrichment and the 95% highest posterior density intervals against the posterior inclusion probability of the term, on gray the rope area around enrichment 1.0(0.5-1.5).

**Figure S17.**
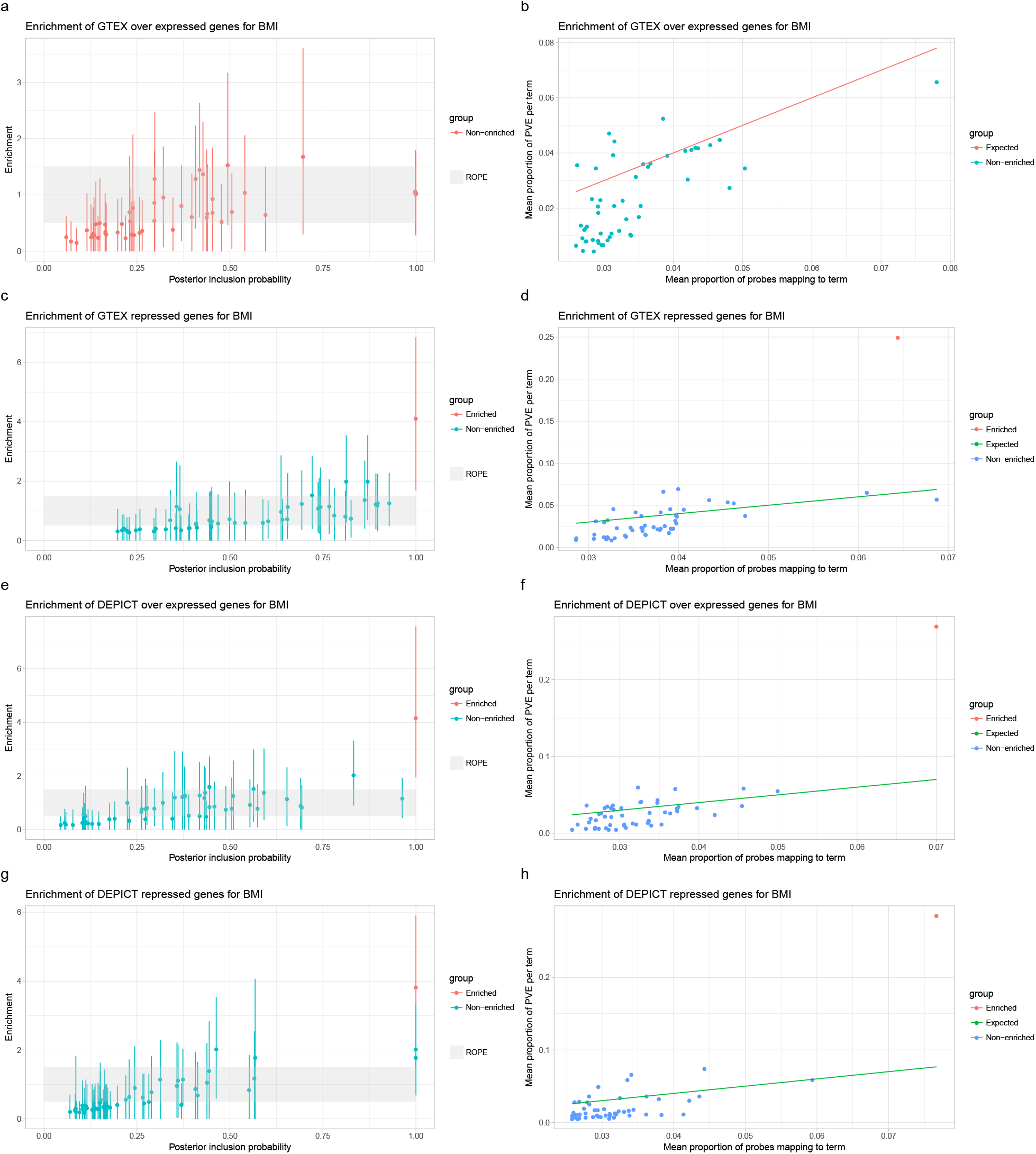
Enrichment analysis for BMI using over expressed(top 99% percentile) and repressed genes (lower 1% percentile) for GTEX and DEPICT

**Figure S18.**
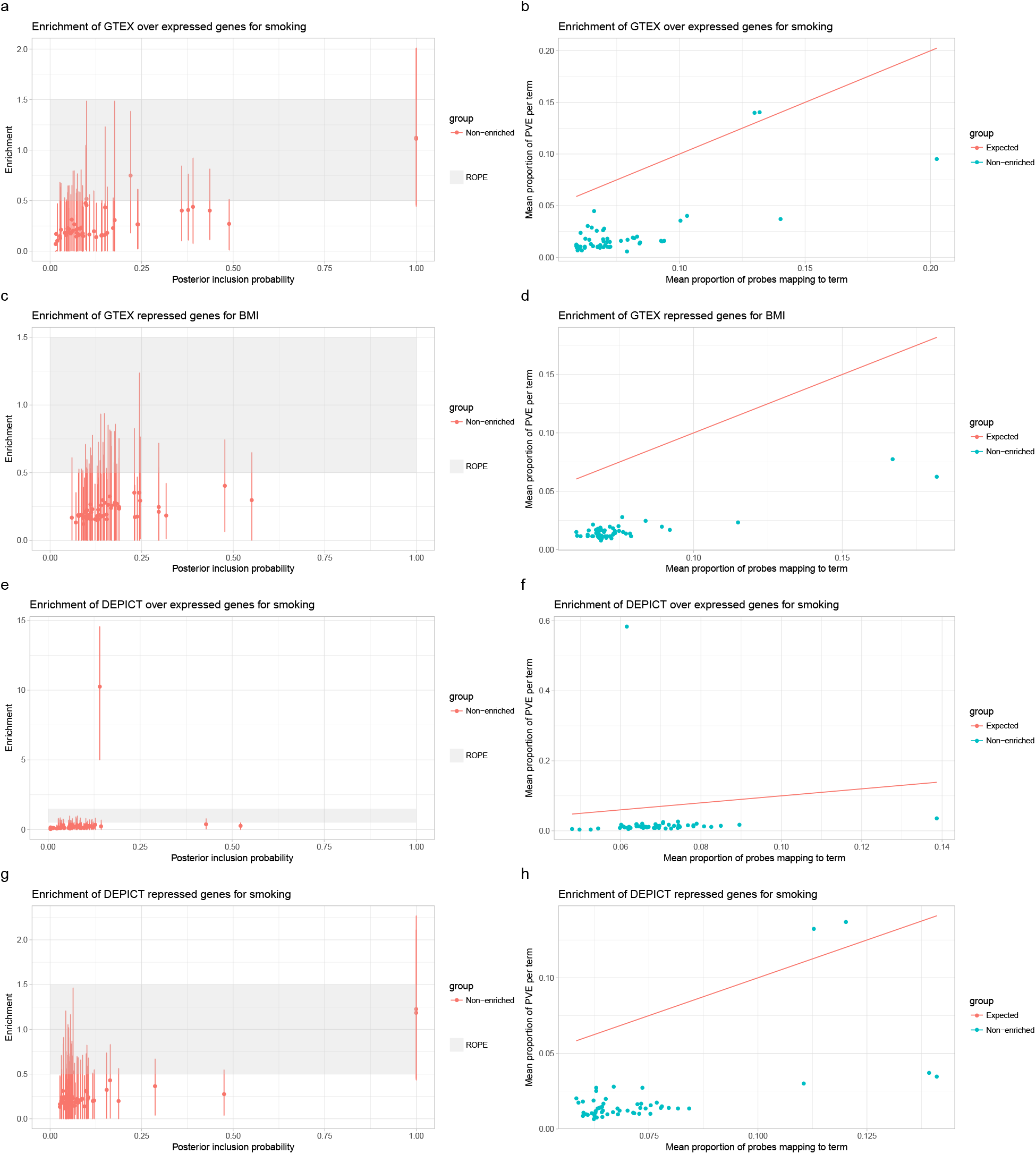
Enrichment analysis for smoking using over expressed(top 99% percentile) and repressed genes (lower 1% percentile) for GTEX and DEPICT

**Table S1.**
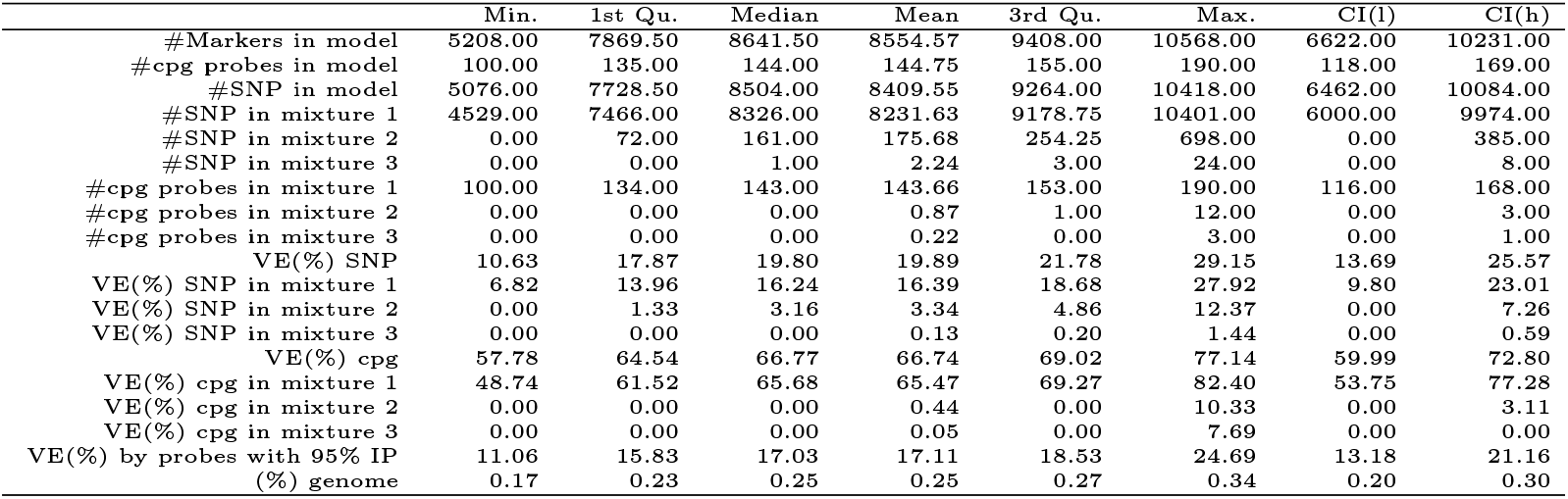
BMI posterior summary

**Table S2.** Smoking posterior summary

|  | Min. | 1st Qu. | Median | Mean | 3rd Qu. | Max. | CI(l) | CI(h) |
| --- | --- | --- | --- | --- | --- | --- | --- | --- |
| #Markers in model | 1741.00 | 3392.75 | 4256.50 | 4369.35 | 5312.75 | 7564.00 | 2084.00 | 6577.00 |
| #cpg probes in model | 32.00 | 51.00 | 59.00 | 59.05 | 66.00 | 99.00 | 37.00 | 78.00 |
| #SNP in model | 1690.00 | 3335.75 | 4197.00 | 4310.16 | 5253.75 | 7501.00 | 2018.00 | 6518.00 |
| #SNP in mixture 1 | 1162.00 | 2946.25 | 3899.00 | 3983.11 | 5018.25 | 7312.00 | 1741.00 | 6321.00 |
| #SNP in mixture 2 | 0.00 | 111.75 | 262.50 | 294.36 | 448.75 | 1118.00 | 0.00 | 660.00 |
| #SNP in mixture 3 | 0.00 | 8.00 | 22.00 | 32.69 | 46.00 | 250.00 | 0.00 | 99.00 |
| #cpg probes in mixture 1 | 27.00 | 48.00 | 56.00 | 56.31 | 64.00 | 91.00 | 35.00 | 77.00 |
| #cpg probes in mixture 2 | 0.00 | 0.00 | 1.00 | 1.63 | 2.00 | 12.00 | 0.00 | 5.00 |
| #cpg probes in mixture 3 | 0.00 | 1.00 | 1.00 | 1.12 | 2.00 | 6.00 | 0.00 | 3.00 |
| VE(%) SNP | 0.00 | 0.00 | 0.27 | 1.86 | 3.15 | 15.73 | 0.00 | 7.35 |
| VE(%) SNP in mixture 1 | 0.00 | 0.00 | 0.10 | 0.81 | 1.17 | 8.68 | 0.00 | 3.58 |
| VE(%) SNP in mixture 2 | 0.00 | 0.00 | 0.04 | 0.74 | 0.93 | 10.09 | 0.00 | 3.99 |
| VE(%) SNP in mixture 3 | 0.00 | 0.00 | 0.02 | 0.31 | 0.33 | 5.47 | 0.00 | 1.57 |
| VE(%) cpg | 51.91 | 64.48 | 67.87 | 67.68 | 71.05 | 80.81 | 58.41 | 76.86 |
| VE(%) cpg in mixture 1 | 11.85 | 22.36 | 26.34 | 26.30 | 29.98 | 44.97 | 15.27 | 37.10 |
| VE(%) cpg in mixture 2 | 0.00 | 0.00 | 0.00 | 9.29 | 8.67 | 65.60 | 0.00 | 46.75 |
| VE(%) cpg in mixture 3 | 0.00 | 0.00 | 0.00 | 9.39 | 12.88 | 69.23 | 0.00 | 49.44 |
| VE(%) by probes with 95% IP | 20.07 | 33.56 | 37.26 | 37.60 | 41.03 | 60.29 | 27.01 | 49.50 |
| (%) genome | 0.06 | 0.09 | 0.11 | 0.11 | 0.12 | 0.19 | 0.07 | 0.15 |

**Table S3.** Number of probes with posterior inclusion probabilities in ranges

|  | <5% | 5%<50% | 50%<95% | >95% |
| --- | --- | --- | --- | --- |
| BMI | 604180 | 605 | 29 | 10 |
| smoking | 587658 | 60 | 9 | 5 |

**Table S4.** Probes with IP > 5% for BMI

|  | cpg | PIP | symbol | group | alias | entrezid |
| --- | --- | --- | --- | --- | --- | --- |
| 8123 | cg00574958 | 1.00 | CPT1A | 5'UTR | CPT1A | 1374 |
| 47302 | cg03546163 | 1.00 | FKBP5 | 5'UTR | FKBP5 | 2289 |
| 73244 | cg05603985 | 1.00 | SKI | 1stExon | SKI | 6497 |
| 83655 | cg06500161 | 1.00 | ABCG1 | Body | ABCG1 | 9619 |
| 202317 | cg16663980 | 1.00 | MNDA | 5'UTR | MNDA | 4332 |
| 218488 | cg18181703 | 1.00 | SOCS3 | Body | SOCS3 | 9021 |
| 311327 | cg26950531 | 1.00 | DPF1 | Body | DPF1 | 8193 |
| 47249 | cg03541746 | 0.78 | CLSTN1 | Body | CLSTN1 | 22883 |
| 164095 | cg13478588 | 0.76 | BAT2 | Body | PRRC2A | 7916 |
| 80188 | cg06192883 | 0.75 | MYO5C | Body | MYO5C | 55930 |
| 85406 | cg06644515 | 0.75 | SNORD78 | TSS200 | SNORD78 | 692198 |
| 85407 | cg06644515 | 0.75 | SNORD80 | TSS1500 | SNORD80 | 26774 |
| 85408 | cg06644515 | 0.75 | GAS5 | Body | GAS5 | 60674 |
| 85409 | cg06644515 | 0.75 | SNORD81 | TSS1500 | SNORD81 | 26769 |
| 85410 | cg06644515 | 0.75 | SNORD79 | TSS1500 | SNORD79 | 26770 |
| 85411 | cg06644515 | 0.75 | SNORD47 | TSS1500 | SNORD47 | 26802 |
| 182583 | cg14870271 | 0.75 | LGALS3BP | 1stExon | LGALS3BP | 3959 |
| 271841 | cg23320499 | 0.75 | DUSP16 | Body | DUSP16 | 80824 |
| 52360 | cg03940776 | 0.72 | SYNJ2 | Body | SYNJ2 | 8871 |
| 71624 | cg05468843 | 0.69 | IL10RA | Body | IL10RA | 3587 |
| 155324 | cg12648759 | 0.67 | ATF6 | Body | ATF6 | 22926 |
| 189751 | cg15564619 | 0.64 | SKI | Body | SKI | 6497 |
| 296902 | cg25648203 | 0.63 | AHRR | Body | AHRR | 57491 |
| 137501 | cg11024682 | 0.60 | SREBF1 | Body | SREBF1 | 6720 |
| 117159 | cg09238801 | 0.59 | PNMA1 | 1stExon | PNMA1 | 9240 |
| 78192 | cg06012428 | 0.58 | ARID1B | Body | ARID1B | 57492 |
| 27479 | cg02043791 | 0.57 | FGFR1 | 5'UTR | FGFR1 | 2260 |
| 280171 | cg24088496 | 0.56 | MAML2 | Body | MAML2 | 84441 |
| 199041 | cg16398761 | 0.55 | C14orf43 | 5'UTR | ELMSAN1 | 91748 |
| 120843 | cg09554443 | 0.53 | CD247 | 1stExon | CD247 | 919 |
| 160053 | cg13123009 | 0.51 | LY6G6D | TSS1500 | LY6G6D | 58530 |
| 160054 | cg13123009 | 0.51 | LY6G6E | TSS200 | LY6G6E | 79136 |
| 36543 | cg02714303 | 0.50 | LMO7 | TSS200 | LMO7 | 4008 |
| 132625 | cg10581837 | 0.50 | LMO7 | TSS200 | LMO7 | 4008 |
| 193906 | cg15939920 | 0.50 | ACACA | Body | ACACA | 31 |
| 228934 | cg19169154 | 0.50 | MFAP4 | Body | MFAP4 | 4239 |
| 86902 | cg06760830 | 0.50 | MOBKL2C | Body | MOB3C | 148932 |
| 189822 | cg15571561 | 0.47 | ARPP-21 | 5'UTR | ARPP21 | 10777 |
| 72893 | cg05575921 | 0.46 | AHRR | Body | AHRR | 57491 |
| 112896 | cg08857797 | 0.44 | VPS25 | Body | VPS25 | 84313 |
| 60070 | cg04535521 | 0.43 | GABRP | TSS1500 | GABRP | 2568 |
| 279229 | cg24009074 | 0.39 | NAPA | Body | NAPA | 8775 |
| 60187 | cg04546097 | 0.39 | HGD | 1stExon | HGD | 3081 |
| 13418 | cg00968616 | 0.39 | CUEDC1 | 3'UTR | CUEDC1 | 404093 |
| 201062 | cg16565294 | 0.38 | TMX1 | Body | TMX1 | 81542 |
| 141370 | cg11376147 | 0.37 | SLC43A1 | Body | SLC43A1 | 8501 |
| 229229 | cg19202384 | 0.36 | PYCR1 | Body | PYCR1 | 5831 |
| 77961 | cg05991220 | 0.35 | ITIH1 | TSS200 | ITIH1 | 3697 |
| 180130 | cg14660446 | 0.34 | NAPA | Body | NAPA | 8775 |
| 21654 | cg01593673 | 0.33 | HOXB3 | 5'UTR | HOXB3 | 3213 |
| 278099 | cg23919844 | 0.33 | ARHGEF7 | Body | ARHGEF7 | 8874 |
| 315271 | cg27309702 | 0.33 | KIFC3 | 3'UTR | KIFC3 | 3801 |
| 173873 | cg14195216 | 0.31 | ANKS1B | TSS1500 | ANKS1B | 56899 |
| 314105 | cg27203924 | 0.31 | ADAMTS9 | Body | ADAMTS9 | 56999 |
| 286696 | cg24687805 | 0.30 | RAB27A | Body | RAB27A | 5873 |
| 100757 | cg07839457 | 0.29 | NLRC5 | TSS1500 | NLRC5 | 84166 |
| 154363 | cg12571570 | 0.29 | ZEB2 | Body | ZEB2 | 9839 |
| 289209 | cg24910161 | 0.28 | GSDMA | TSS200 | GSDMA | 284110 |
| 248992 | cg21113478 | 0.28 | DCLK3 | TSS1500 | DCLK3 | 85443 |
| 100596 | cg07826387 | 0.28 | CHRM3 | Body | CHRM3 | 1131 |
| 267034 | cg22884440 | 0.28 | ASAP1 | Body | ASAP1 | 50807 |
| 318020 | cg27554156 | 0.27 | GSG1 | Body | GSG1 | 83445 |
| 176726 | cg14410503 | 0.26 | TNS3 | Body | TNS3 | 64759 |
| 202290 | cg16662477 | 0.26 | RFFL | 5'UTR | RFFL | 117584 |
| 144815 | cg11704631 | 0.25 | RUNX1 | Body | RUNX1 | 861 |
| 214557 | cg17836177 | 0.25 | PEBP4 | Body | PEBP4 | 157310 |
| 215706 | cg17942763 | 0.25 | SKI | Body | SKI | 6497 |
| 239464 | cg20146909 | 0.25 | LRRC8D | 5'UTR | LRRC8D | 55144 |
| 108086 | cg08447387 | 0.25 | ASAP1 | Body | ASAP1 | 50807 |
| 49227 | cg03691549 | 0.25 | TENC1 | 5'UTR | TNS2 | 23371 |
| 48539 | cg03636183 | 0.25 | F2RL3 | Body | F2RL3 | 9002 |
| 272572 | cg23384027 | 0.24 | NFE2 | Body | NFE2 | 4778 |
| 252549 | cg21468274 | 0.24 | TMEM102 | Body | TMEM102 | 284114 |
| 139862 | cg11236746 | 0.23 | GLB1 | Body | GLB1 | 2720 |
| 169031 | cg13844237 | 0.22 | MAPKAP1 | Body | MAPKAP1 | 79109 |
| 152051 | cg12358527 | 0.21 | BCAS3 | Body | BCAS3 | 54828 |
| 49666 | cg03725309 | 0.20 | SARS | Body | SARS | 6301 |
| 114965 | cg09039672 | 0.20 | ACVR1B | Body | ACVR1B | 91 |
| 150676 | cg12216054 | 0.20 | ZBTB9 | Body | ZBTB9 | 221504 |
| 4453 | cg00318947 | 0.20 | HOXA3 | 5'UTR | HOXA3 | 3200 |
| 166891 | cg13680388 | 0.20 | GNAS | Body | GNAS | 2778 |
| 198683 | cg16365807 | 0.19 | BACE2 | Body | BACE2 | 25825 |
| 252984 | cg21507987 | 0.19 | AMPD3 | 3'UTR | AMPD3 | 272 |
| 310978 | cg26921611 | 0.19 | EIF4EBP1 | 3'UTR | EIF4EBP1 | 1978 |
| 116024 | cg09138671 | 0.18 | TMCO4 | 5'UTR | TMCO4 | 255104 |
| 16575 | cg01199327 | 0.18 | CD82 | Body | CD82 | 3732 |
| 37452 | cg02780017 | 0.18 | SLC22A20 | TSS200 | SLC22A20P | 440044 |
| 132827 | cg10596609 | 0.18 | UGT1A8 | TSS1500 | UGT1A8 | 54576 |
| 53215 | cg04006327 | 0.17 | ATG16L2 | Body | ATG16L2 | 89849 |
| 104312 | cg08135379 | 0.17 | AMIGO2 | TSS1500 | AMIGO2 | 347902 |
| 72799 | cg05568797 | 0.16 | BHLHE41 | Body | BHLHE41 | 79365 |
| 108433 | cg08472795 | 0.16 | HS6ST1 | Body | HS6ST1 | 9394 |
| 114151 | cg08958168 | 0.16 | SLC12A4 | Body | SLC12A4 | 6560 |
| 303178 | cg26253134 | 0.16 | TGFA | Body | TGFA | 7039 |
| 63986 | cg04858631 | 0.15 | DDIT4 | 3'UTR | DDIT4 | 54541 |
| 309971 | cg26841040 | 0.15 | GUK1 | TSS200 | GUK1 | 2987 |
| 24683 | cg01831904 | 0.14 | LRRC37B2 | Body | LRRC37BP1 | 147172 |
| 313550 | cg27153059 | 0.14 | TMEM86A | TSS1500 | TMEM86A | 144110 |
| 200256 | cg16500874 | 0.14 | LETM1 | Body | LETM1 | 3954 |
| 82942 | cg06437430 | 0.14 | NKD2 | Body | NKD2 | 85409 |
| 130324 | cg10381071 | 0.14 | TLE3 | TSS1500 | TLE3 | 7090 |
| 216332 | cg18004701 | 0.14 | PTPRN2 | Body | PTPRN2 | 5799 |
| 213527 | cg17741339 | 0.14 | ESR1 | 5'UTR | ESR1 | 2099 |
| 318414 | cg27588321 | 0.13 | CYLD | TSS1500 | CYLD | 1540 |
| 138174 | cg11087049 | 0.13 | NIN | 5'UTR | NIN | 51199 |
| 42461 | cg03162314 | 0.13 | CTU1 | 3'UTR | CTU1 | 90353 |
| 301103 | cg26066277 | 0.13 | DYSFIP1 | 1stExon | PPP1R27 | 116729 |
| 60942 | cg04604050 | 0.12 | SYNPO | Body | SYNPO | 11346 |
| 175684 | cg14323854 | 0.12 | ZNF177 | TSS1500 | ZNF177 | 7730 |
| 40490 | cg03010301 | 0.12 | MIR125A | TSS1500 | MIR125A | 406910 |
| 40491 | cg03010301 | 0.12 | MIR99B | TSS1500 | MIR99B | 407056 |
| 40492 | cg03010301 | 0.12 | MIRLET7E | TSS1500 | MIRLET7E | 406887 |
| 40493 | cg03010301 | 0.12 | NCRNA00085 | TSS1500 | SPACA6 | 147650 |
| 206049 | cg17024258 | 0.12 | VANGL2 | 5'UTR | VANGL2 | 57216 |
| 86442 | cg06728793 | 0.12 | PRKAR1B | Body | PRKAR1B | 5575 |
| 127850 | cg10169539 | 0.12 | NIN | 5'UTR | NIN | 51199 |
| 166132 | cg13624385 | 0.12 | PCDH7 | TSS200 | PCDH7 | 5099 |
| 268588 | cg23032421 | 0.12 | IL5RA | 5'UTR | IL5RA | 3568 |
| 73879 | cg05655915 | 0.11 | NARF | TSS1500 | NARF | 26502 |
| 299449 | cg25919221 | 0.11 | CA6 | Body | CA6 | 765 |
| 126192 | cg10028434 | 0.11 | KLC1 | Body | KLC1 | 3831 |
| 35143 | cg02612927 | 0.11 | DEFB124 | TSS200 | DEFB124 | 245937 |
| 253461 | cg21550372 | 0.11 | WDR25 | Body | WDR25 | 79446 |
| 121198 | cg09581649 | 0.11 | C16orf5 | 5'UTR | CDIP1 | 29965 |
| 232335 | cg19502457 | 0.11 | GPX2 | 1stExon | GPX2 | 2877 |
| 153632 | cg12499235 | 0.11 | ASCL2 | TSS1500 | ASCL2 | 430 |
| 192769 | cg15833940 | 0.11 | DLGAP2 | Body | DLGAP2 | 9228 |
| 111218 | cg08711064 | 0.10 | RASA1 | 1stExon | RASA1 | 5921 |
| 54750 | cg04125273 | 0.10 | ZG16 | Body | ZG16 | 653808 |
| 134402 | cg10738119 | 0.10 | YWHAG | 3'UTR | YWHAG | 7532 |
| 44875 | cg03358113 | 0.10 | MAD1L1 | Body | MAD1L1 | 8379 |
| 92999 | cg07217499 | 0.10 | CACNA1C | Body | CACNA1C | 775 |
| 182945 | cg14906103 | 0.10 | TMEM33 | TSS1500 | TMEM33 | 55161 |
| 285220 | cg24550026 | 0.10 | BCL2L15 | 1stExon | BCL2L15 | 440603 |
| 290230 | cg25006231 | 0.10 | C10orf41 | Body | ZNF503-AS2 | 100131213 |
| 226305 | cg18896032 | 0.10 | SSRP1 | Body | SSRP1 | 6749 |
| 295293 | cg25496297 | 0.10 | MYO18A | Body | MYO18A | 399687 |
| 88952 | cg06914133 | 0.09 | SAMD4A | Body | SAMD4A | 23034 |
| 122791 | cg09728393 | 0.09 | KIF13A | Body | KIF13A | 63971 |
| 232304 | cg19499748 | 0.09 | TRIL | 5'UTR | TRIL | 9865 |
| 243115 | cg20525527 | 0.09 | MPRIP | Body | MPRIP | 23164 |
| 28966 | cg02154448 | 0.09 | OVGP1 | Body | OVGP1 | 5016 |
| 117237 | cg09245872 | 0.09 | DOC2A | 3'UTR | DOC2A | 8448 |
| 117238 | cg09245872 | 0.09 | INO80E | 3'UTR | INO80E | 283899 |
| 224499 | cg18734095 | 0.09 | NOSIP | Body | NOSIP | 51070 |
| 30290 | cg02257517 | 0.09 | SLC2A6 | Body | SLC2A6 | 11182 |
| 49575 | cg03719040 | 0.09 | TMEM128 | TSS1500 | TMEM128 | 85013 |
| 30438 | cg02268192 | 0.09 | RIN3 | Body | RIN3 | 79890 |
| 183696 | cg14978174 | 0.09 | TIPARP | TSS1500 | TIPARP | 25976 |
| 24805 | cg01839993 | 0.09 | DDIT4 | Body | DDIT4 | 54541 |
| 179488 | cg14609407 | 0.09 | SLPI | 1stExon | SLPI | 6590 |
| 168842 | cg13828047 | 0.09 | MPI | TSS1500 | MPI | 4351 |
| 302185 | cg26160945 | 0.09 | STK38L | TSS1500 | STK38L | 23012 |
| 24042 | cg01787285 | 0.08 | SKI | Body | SKI | 6497 |
| 238199 | cg20018057 | 0.08 | GNAS | 3'UTR | GNAS | 2778 |
| 291873 | cg25158622 | 0.08 | BRD2 | Body | BRD2 | 6046 |
| 269400 | cg23097139 | 0.08 | SNUPN | TSS1500 | SNUPN | 10073 |
| 88447 | cg06876354 | 0.08 | RALB | 5'UTR | RALB | 5899 |
| 235263 | cg19759478 | 0.08 | C1orf183 | Body | FAM212B | 55924 |
| 14213 | cg01024458 | 0.08 | RERE | Body | RERE | 473 |
| 144156 | cg11649376 | 0.08 | ACSS3 | Body | ACSS3 | 79611 |
| 301889 | cg26134737 | 0.08 | DIP2C | Body | DIP2C | 22982 |
| 247406 | cg20961940 | 0.08 | SLC44A4 | Body | SLC44A4 | 80736 |
| 3975 | cg00285294 | 0.08 | C7orf50 | Body | C7orf50 | 84310 |
| 130104 | cg10363549 | 0.08 | FAM190A | 3'UTR | CCSER1 | 401145 |
| 215898 | cg17961943 | 0.08 | PLB1 | Body | PLB1 | 151056 |
| 75682 | cg05806375 | 0.08 | ERAP1 | TSS200 | ERAP1 | 51752 |
| 5088 | cg00363312 | 0.07 | HIPK2 | Body | HIPK2 | 28996 |
| 225627 | cg18835596 | 0.07 | MYB | 3'UTR | MYB | 4602 |
| 275765 | cg23685712 | 0.07 | CYP24A1 | 1stExon | CYP24A1 | 1591 |
| 275750 | cg23684682 | 0.07 | ZNF30 | 5'UTR | ZNF30 | 90075 |
| 247745 | cg20991801 | 0.07 | CDK13 | 1stExon | CDK13 | 8621 |
| 148173 | cg11992736 | 0.07 | IL12RB1 | Body | IL12RB1 | 3594 |
| 178602 | cg14547895 | 0.07 | PKNOX2 | 5'UTR | PKNOX2 | 63876 |
| 46638 | cg03498559 | 0.07 | UGT2B4 | 1stExon | UGT2B4 | 7363 |
| 111408 | cg08726900 | 0.07 | ANKRD11 | 5'UTR | ANKRD11 | 29123 |
| 190687 | cg15643574 | 0.07 | PICALM | Body | PICALM | 8301 |
| 207122 | cg17124844 | 0.07 | DLGAP2 | Body | DLGAP2 | 9228 |
| 277975 | cg23909013 | 0.07 | KCNJ6 | TSS1500 | KCNJ6 | 3763 |
| 302267 | cg26168117 | 0.07 | MDN1 | 3'UTR | MDN1 | 23195 |
| 11170 | cg00799631 | 0.06 | TNRC18 | Body | TNRC18 | 84629 |
| 283613 | cg24410469 | 0.06 | AGAP1 | Body | AGAP1 | 116987 |
| 34922 | cg02595681 | 0.06 | ZNF215 | TSS1500 | ZNF215 | 7762 |
| 39636 | cg02949469 | 0.06 | PNKD | 1stExon | PNKD | 25953 |
| 39637 | cg02949469 | 0.06 | AAMP | TSS1500 | AAMP | 14 |
| 112938 | cg08861449 | 0.06 | CSMD2 | 3'UTR | CSMD2 | 114784 |
| 269770 | cg23137881 | 0.06 | C4orf45 | Body | C4orf45 | 152940 |
| 84460 | cg06568880 | 0.06 | SMG6 | Body | SMG6 | 23293 |
| 204181 | cg16828491 | 0.06 | SH3TC1 | Body | SH3TC1 | 54436 |
| 297748 | cg25736198 | 0.06 | TPH2 | Body | TPH2 | 121278 |
| 167343 | cg13711063 | 0.06 | KCNJ8 | 3'UTR | KCNJ8 | 3764 |
| 225593 | cg18832247 | 0.06 | JARID2 | Body | JARID2 | 3720 |
| 203106 | cg16725642 | 0.06 | EPHB4 | Body | EPHB4 | 2050 |
| 203751 | cg16786297 | 0.06 | SDCBP2 | 3'UTR | SDCBP2 | 27111 |
| 231754 | cg19445335 | 0.06 | ADAP1 | Body | ADAP1 | 11033 |
| 74099 | cg05670596 | 0.06 | CCRL2 | TSS1500 | CCRL2 | 9034 |
| 126644 | cg10069470 | 0.06 | C5orf30 | 5'UTR | C5orf30 | 90355 |
| 198521 | cg16352321 | 0.06 | TP53INP2 | 3'UTR | TP53INP2 | 58476 |
| 9594 | cg00684043 | 0.06 | DNASE1L2 | 3'UTR | DNASE1L2 | 1775 |
| 13937 | cg01004980 | 0.06 | PRKAR2A | Body | PRKAR2A | 5576 |
| 171719 | cg14034270 | 0.06 | MEG3 | TSS1500 | MEG3 | 55384 |
| 189013 | cg15488009 | 0.06 | FSTL3 | Body | FSTL3 | 10272 |
| 96748 | cg07525299 | 0.06 | ASCL2 | TSS1500 | ASCL2 | 430 |
| 124924 | cg09914675 | 0.06 | SEPX1 | 1stExon | MSRB1 | 51734 |
| 182141 | cg14836313 | 0.06 | MRPL23 | Body | MRPL23 | 6150 |
| 277356 | cg23850913 | 0.06 | JARID2 | Body | JARID2 | 3720 |
| 245804 | cg20784591 | 0.05 | PILRA | Body | PILRA | 29992 |
| 164629 | cg13520520 | 0.05 | ECSIT | 3'UTR | ECSIT | 51295 |
| 164630 | cg13520520 | 0.05 | ZNF653 | TSS1500 | ZNF653 | 115950 |
| 278162 | cg23925173 | 0.05 | RASAL1 | 3'UTR | RASAL1 | 8437 |
| 109487 | cg08561775 | 0.05 | UBE2O | Body | UBE2O | 63893 |
| 286590 | cg24681499 | 0.05 | FOXJ3 | Body | FOXJ3 | 22887 |
| 237319 | cg19936912 | 0.05 | FAM160A1 | TSS1500 | FAM160A1 | 729830 |
| 283498 | cg24401026 | 0.05 | MAP7 | Body | MAP7 | 9053 |
| 314602 | cg27253454 | 0.05 | USP43 | Body | USP43 | 124739 |

**Table S5.** Probes with IP > 5% for smoking

|  | cpg | PIP | symbol | group | alias | entrezid |
| --- | --- | --- | --- | --- | --- | --- |
| 48539 | cg03636183 | 1.00 | F2RL3 | Body | F2RL3 | 9002 |
| 72893 | cg05575921 | 1.00 | AHRR | Body | AHRR | 57491 |
| 224710 | cg18754985 | 1.00 | CLDND1 | Body | CLDND1 | 56650 |
| 129621 | cg10321266 | 0.91 | MBNL1 | 5'UTR | MBNL1 | 4154 |
| 140769 | cg11321090 | 0.89 | ZFYVE28 | Body | ZFYVE28 | 57732 |
| 132514 | cg10573386 | 0.84 | CELSR3 | TSS200 | CELSR3 | 1951 |
| 139459 | cg11207515 | 0.83 | CNTNAP2 | Body | CNTNAP2 | 26047 |
| 153879 | cg12524061 | 0.67 | SDR9C7 | TSS1500 | SDR9C7 | 121214 |
| 175434 | cg14305420 | 0.65 | NOXA1 | Body | NOXA1 | 10811 |
| 192021 | cg15763258 | 0.56 | FLI1 | TSS1500 | FLI1 | 2313 |
| 155345 | cg12651044 | 0.46 | APH1A | TSS200 | APH1A | 51107 |
| 97753 | cg07597069 | 0.45 | PLEC1 | Body | PLEC | 5339 |
| 305460 | cg26450010 | 0.45 | ABHD8 | Body | ABHD8 | 79575 |
| 117421 | cg09256832 | 0.36 | SPSB1 | 5'UTR | SPSB1 | 80176 |
| 68570 | cg05218653 | 0.31 | ABCA3 | Body | ABCA3 | 21 |
| 244423 | cg20661303 | 0.31 | LEFTY2 | TSS1500 | LEFTY2 | 7044 |
| 49424 | cg03707168 | 0.30 | PPP1R15A | Body | PPP1R15A | 23645 |
| 223203 | cg18607609 | 0.23 | IQGAP2 | TSS1500 | IQGAP2 | 10788 |
| 67319 | cg05119988 | 0.21 | SC4MOL | 5'UTR | MSMO1 | 6307 |
| 229657 | cg19246197 | 0.20 | USP2 | Body | USP2 | 9099 |
| 80786 | cg06241380 | 0.19 | CTCF | TSS1500 | CTCF | 10664 |
| 138134 | cg11084015 | 0.16 | OSBPL10 | Body | OSBPL10 | 114884 |
| 143182 | cg11560197 | 0.15 | PCDHA3 | 1stExon | PCDHA3 | 56145 |
| 143183 | cg11560197 | 0.15 | PCDHA1 | Body | PCDHA1 | 56147 |
| 143184 | cg11560197 | 0.15 | PCDHA2 | Body | PCDHA2 | 56146 |
| 217745 | cg18119192 | 0.15 | GATA4 | Body | GATA4 | 2626 |
| 199041 | cg16398761 | 0.14 | C14orf43 | 5'UTR | ELMSAN1 | 91748 |
| 95978 | cg07468289 | 0.14 | NCRNA00171 | TSS1500 | ZNRD1ASP | 80862 |
| 95979 | cg07468289 | 0.14 | ZNRD1 | Body | ZNRD1 | 30834 |
| 261719 | cg22371743 | 0.13 | MATN1 | Body | MATN1 | 4146 |
| 135796 | cg10861146 | 0.13 | EFNA1 | TSS1500 | EFNA1 | 1942 |
| 281164 | cg24169820 | 0.12 | CAMKK2 | Body | CAMKK2 | 10645 |
| 160873 | cg13209613 | 0.11 | WNT3 | Body | WNT3 | 7473 |
| 173423 | cg14163311 | 0.11 | RASSF5 | Body | RASSF5 | 83593 |
| 236825 | cg19895882 | 0.10 | KIAA0564 | Body | VWA8 | 23078 |
| 223693 | cg18654464 | 0.10 | MIR1227 | Body | MIR1227 | 100302283 |
| 223694 | cg18654464 | 0.10 | PLEKHJ1 | Body | PLEKHJ1 | 55111 |
| 245612 | cg20764780 | 0.09 | PTPRN2 | Body | PTPRN2 | 5799 |
| 95557 | cg07435237 | 0.07 | ARMC5 | TSS1500 | ARMC5 | 79798 |
| 185843 | cg15164708 | 0.07 | SLC17A7 | Body | SLC17A7 | 57030 |
| 37002 | cg02746869 | 0.07 | VIM | Body | VIM | 7431 |
| 176229 | cg14371731 | 0.07 | ZMIZ1 | Body | ZMIZ1 | 57178 |
| 309591 | cg26813561 | 0.07 | ACAD10 | Body | ACAD10 | 80724 |
| 18941 | cg01389283 | 0.07 | UBQLNL | TSS1500 | UBQLNL | 143630 |
| 108347 | cg08466269 | 0.07 | DHRS12 | Body | DHRS12 | 79758 |
| 94457 | cg07339236 | 0.06 | ATP9A | Body | ATP9A | 10079 |
| 147135 | cg11897145 | 0.06 | ETV1 | 5'UTR | ETV1 | 2115 |
| 196297 | cg16163140 | 0.06 | ADAMTS16 | Body | ADAMTS16 | 170690 |
| 316610 | cg27425996 | 0.06 | COBL | Body | COBL | 23242 |
| 170690 | cg13967864 | 0.06 | NCBP1 | 1stExon | NCBP1 | 4686 |
| 170691 | cg13967864 | 0.06 | TSTD2 | TSS200 | TSTD2 | 158427 |
| 192934 | cg15847614 | 0.06 | FAM64A | 5'UTR | PIMREG | 54478 |
| 211054 | cg17480760 | 0.06 | CA3 | TSS200 | CA3 | 761 |
| 100013 | cg07782603 | 0.06 | GALNT9 | Body | GALNT9 | 50614 |
| 110477 | cg08647039 | 0.06 | NFIX | Body | NFIX | 4784 |
| 200001 | cg16476048 | 0.06 | C10orf122 | TSS1500 | TEX36 | 387718 |
| 57912 | cg04367503 | 0.05 | NFIX | Body | NFIX | 4784 |
| 65751 | cg04992150 | 0.05 | NUP210 | Body | NUP210 | 23225 |
| 163182 | cg13415795 | 0.05 | SH2D7 | 3'UTR | SH2D7 | 646892 |
| 238783 | cg20077343 | 0.05 | MUC6 | Body | MUC6 | 4588 |
| 243459 | cg20560182 | 0.05 | TAF7 | TSS200 | TAF7 | 6879 |

**Table S6.** GO enrichment for the probes with 95%IP in BMI

|  | Term | P.DE |
| --- | --- | --- |
| GO:0032365 | intracellular lipid transport | 0.00 |
| GO:0009720 | detection of hormone stimulus | 0.00 |
| GO:0034437 | glycoprotein transporter activity | 0.00 |
| GO:0034041 | sterol-transporting ATPase activity | 0.00 |
| GO:1990698 | positive regulation of cytosolic calcium ion... | 0.00 |
| GO:0055099 | response to high density lipoprotein particle | 0.00 |
| GO:0034040 | lipid-transporting ATPase activity | 0.00 |
| GO:0009726 | detection of endogenous stimulus | 0.00 |
| GO:0046811 | histone deacetylase inhibitor activity | 0.00 |
| GO:0045834 | positive regulation of lipid metabolic process | 0.00 |

**Table S7.** GO enrichment for the probes with 95%IP in smoking

|  | Term | P.DE |
| --- | --- | --- |
| GO:0015057 | thrombin-activated receptor activity | 0.00 |
| GO:0060155 | platelet dense granule organization | 0.00 |
| GO:0070493 | thrombin-activated receptor signaling pathway | 0.00 |
| GO:0035025 | positive regulation of Rho protein signal transduction | 0.00 |
| GO:0051482 | positive regulation of cytosolic calcium ... | 0.00 |
| GO:0033363 | secretory granule organization | 0.01 |
| GO:0051281 | positive regulation of release of ... | 0.01 |
| GO:0010524 | positive regulation of calcium ion ... | 0.01 |
| GO:0046579 | positive regulation of Ras protein signal transduction | 0.01 |
| GO:0001191 | transcriptional repressor activity, RNA polymerase II ... | 0.01 |

**Table S8.** Enrichment summary of over-expressed genes in GTEx for BMI

|  | tissue | mean_pprobes | mean_pve | Min. | 1st Qu. | Median | Mean | 3rd Qu. | Max. | CI(l) | CI(h) | ROPE | prob | group |
| --- | --- | --- | --- | --- | --- | --- | --- | --- | --- | --- | --- | --- | --- | --- |
| 1 | Adipose_Subcutaneous | 0.03 | 0.01 | 0.00 | 0.11 | 0.25 | 0.34 | 0.47 | 1.44 | 0.00 | 0.92 | 0.05 | 0.20 | Non-enriched |
| 2 | Adipose_Visceral_Omentum. | 0.03 | 0.03 | 0.18 | 0.93 | 1.24 | 1.28 | 1.59 | 3.08 | 0.40 | 2.22 | 0.27 | 0.41 | Non-enriched |
| 3 | Adrenal_Gland | 0.03 | 0.05 | 0.14 | 1.00 | 1.53 | 1.68 | 2.18 | 4.96 | 0.30 | 3.61 | 0.30 | 0.70 | Non-enriched |
| 4 | Artery_Aorta | 0.03 | 0.02 | 0.00 | 0.33 | 0.65 | 0.68 | 0.96 | 2.44 | 0.00 | 1.49 | 0.26 | 0.45 | Non-enriched |
| 5 | Artery_Coronary | 0.03 | 0.02 | 0.01 | 0.41 | 0.66 | 0.70 | 0.96 | 1.89 | 0.04 | 1.38 | 0.31 | 0.50 | Non-enriched |
| 6 | Artery_Tibial | 0.03 | 0.01 | 0.00 | 0.17 | 0.32 | 0.39 | 0.56 | 1.54 | 0.00 | 0.95 | 0.11 | 0.35 | Non-enriched |
| 7 | Bladder | 0.03 | 0.01 | 0.00 | 0.33 | 0.50 | 0.54 | 0.70 | 1.32 | 0.00 | 1.12 | 0.12 | 0.23 | Non-enriched |
| 8 | Brain_Amygdala | 0.04 | 0.04 | 0.02 | 0.75 | 0.99 | 1.03 | 1.26 | 2.66 | 0.29 | 1.75 | 0.82 | 1.00 | Non-enriched |
| 9 | Brain_Anterior_cingulate_cortex_BA24. | 0.04 | 0.04 | 0.02 | 0.74 | 0.97 | 1.02 | 1.26 | 2.52 | 0.32 | 1.75 | 0.82 | 1.00 | Non-enriched |
| 10 | Brain_Caudate_basal_ganglia. | 0.04 | 0.04 | 0.17 | 0.80 | 1.02 | 1.06 | 1.29 | 2.86 | 0.33 | 1.75 | 0.82 | 1.00 | Non-enriched |
| 11 | Brain_Cerebellar_Hemisphere | 0.03 | 0.01 | 0.00 | 0.08 | 0.20 | 0.29 | 0.43 | 1.64 | 0.00 | 0.87 | 0.04 | 0.24 | Non-enriched |
| 12 | Brain_Cerebellum | 0.03 | 0.01 | 0.00 | 0.07 | 0.17 | 0.23 | 0.32 | 1.36 | 0.00 | 0.63 | 0.03 | 0.22 | Non-enriched |
| 13 | Brain_Cortex | 0.04 | 0.04 | 0.02 | 0.75 | 0.98 | 1.03 | 1.26 | 2.52 | 0.32 | 1.75 | 0.82 | 1.00 | Non-enriched |
| 14 | Brain_Frontal_Cortex_BA9. | 0.04 | 0.04 | 0.02 | 0.75 | 0.97 | 1.02 | 1.26 | 2.52 | 0.33 | 1.75 | 0.82 | 1.00 | Non-enriched |
| 15 | Brain_Hippocampus | 0.05 | 0.04 | 0.02 | 0.73 | 0.97 | 1.01 | 1.25 | 2.52 | 0.32 | 1.75 | 0.82 | 1.00 | Non-enriched |
| 16 | Brain_Hypothalamus | 0.04 | 0.04 | 0.02 | 0.76 | 0.99 | 1.04 | 1.26 | 2.86 | 0.29 | 1.79 | 0.82 | 1.00 | Non-enriched |
| 17 | Brain_Nucleus_accumbens_basal_ganglia. | 0.05 | 0.04 | 0.02 | 0.76 | 0.97 | 1.01 | 1.22 | 2.52 | 0.35 | 1.69 | 0.85 | 1.00 | Non-enriched |
| 18 | Brain_Putamen_basal_ganglia. | 0.04 | 0.04 | 0.17 | 0.79 | 1.00 | 1.04 | 1.24 | 2.52 | 0.46 | 1.75 | 0.86 | 1.00 | Non-enriched |
| 19 | Brain_Spinal_cord_cervical_c1. | 0.04 | 0.04 | 0.00 | 0.63 | 0.99 | 1.04 | 1.41 | 3.00 | 0.00 | 2.06 | 0.33 | 0.54 | Non-enriched |
| 20 | Brain_Substantia_nigra | 0.03 | 0.02 | 0.00 | 0.33 | 0.59 | 0.66 | 0.94 | 2.47 | 0.00 | 1.53 | 0.23 | 0.44 | Non-enriched |
| 21 | Breast_Mammary_Tissue | 0.03 | 0.03 | 0.00 | 0.54 | 0.94 | 0.93 | 1.25 | 2.90 | 0.10 | 1.83 | 0.29 | 0.45 | Non-enriched |
| 22 | Cells_EBV.transformed_lymphocytes | 0.03 | 0.01 | 0.00 | 0.10 | 0.19 | 0.29 | 0.41 | 1.47 | 0.00 | 0.95 | 0.02 | 0.13 | Non-enriched |
| 23 | Cells_Transformed_fibroblasts | 0.03 | 0.01 | 0.00 | 0.10 | 0.21 | 0.29 | 0.40 | 1.21 | 0.00 | 0.89 | 0.04 | 0.24 | Non-enriched |
| 24 | Cervix_Ectocervix | 0.03 | 0.01 | 0.00 | 0.13 | 0.27 | 0.32 | 0.45 | 1.03 | 0.00 | 0.84 | 0.03 | 0.17 | Non-enriched |
| 25 | Cervix_Endocervix | 0.03 | 0.01 | 0.00 | 0.13 | 0.26 | 0.30 | 0.42 | 1.05 | 0.00 | 0.85 | 0.02 | 0.17 | Non-enriched |
| 26 | Colon_Sigmoid | 0.08 | 0.07 | 0.00 | 0.18 | 0.41 | 0.60 | 0.79 | 3.61 | 0.00 | 1.80 | 0.14 | 0.44 | Non-enriched |
| 27 | Colon_Transverse | 0.04 | 0.03 | 0.00 | 0.26 | 0.59 | 0.69 | 1.02 | 3.16 | 0.00 | 1.68 | 0.10 | 0.23 | Non-enriched |
| 28 | Esophagus_Gastroesophageal_Junction | 0.04 | 0.02 | 0.00 | 0.36 | 0.56 | 0.65 | 0.87 | 2.54 | 0.00 | 1.50 | 0.31 | 0.60 | Non-enriched |
| 29 | Esophagus_Mucosa | 0.03 | 0.00 | 0.00 | 0.06 | 0.14 | 0.17 | 0.26 | 0.64 | 0.00 | 0.53 | 0.00 | 0.07 | Non-enriched |
| 30 | Esophagus_Muscularis | 0.04 | 0.02 | 0.00 | 0.27 | 0.47 | 0.52 | 0.73 | 1.99 | 0.00 | 1.19 | 0.22 | 0.48 | Non-enriched |
| 31 | Fallopian_Tube | 0.03 | 0.04 | 0.33 | 1.04 | 1.34 | 1.44 | 1.75 | 3.98 | 0.60 | 2.63 | 0.25 | 0.42 | Non-enriched |
| 32 | Heart_Atrial_Appendage | 0.03 | 0.02 | 0.00 | 0.18 | 0.40 | 0.50 | 0.80 | 1.58 | 0.00 | 1.29 | 0.06 | 0.15 | Non-enriched |
| 33 | Heart_Left_Ventricle | 0.03 | 0.01 | 0.00 | 0.14 | 0.28 | 0.34 | 0.48 | 1.56 | 0.00 | 0.88 | 0.04 | 0.17 | Non-enriched |
| 34 | Kidney_Cortex | 0.03 | 0.00 | 0.00 | 0.05 | 0.11 | 0.15 | 0.23 | 0.69 | 0.00 | 0.41 | 0.00 | 0.09 | Non-enriched |
| 35 | Liver | 0.04 | 0.05 | 0.00 | 1.00 | 1.35 | 1.37 | 1.78 | 3.54 | 0.00 | 2.30 | 0.21 | 0.43 | Non-enriched |
| 36 | Lung | 0.03 | 0.02 | 0.00 | 0.25 | 0.58 | 0.77 | 1.11 | 2.86 | 0.00 | 2.07 | 0.09 | 0.24 | Non-enriched |
| 37 | Minor_Salivary_Gland | 0.04 | 0.04 | 0.01 | 0.62 | 0.90 | 0.96 | 1.28 | 2.90 | 0.07 | 1.85 | 0.22 | 0.32 | Non-enriched |
| 38 | Muscle_Skeletal | 0.03 | 0.01 | 0.00 | 0.12 | 0.24 | 0.33 | 0.46 | 1.81 | 0.00 | 0.96 | 0.05 | 0.26 | Non-enriched |
| 39 | Nerve_Tibial | 0.05 | 0.03 | 0.00 | 0.22 | 0.49 | 0.54 | 0.82 | 1.89 | 0.00 | 1.28 | 0.14 | 0.30 | Non-enriched |
| 40 | Ovary | 0.03 | 0.01 | 0.00 | 0.09 | 0.21 | 0.24 | 0.32 | 0.76 | 0.00 | 0.63 | 0.02 | 0.15 | Non-enriched |
| 41 | Pancreas | 0.03 | 0.02 | 0.00 | 0.52 | 0.82 | 0.86 | 1.16 | 2.46 | 0.01 | 1.77 | 0.20 | 0.30 | Non-enriched |
| 42 | Pituitary | 0.05 | 0.03 | 0.00 | 0.31 | 0.54 | 0.60 | 0.85 | 1.77 | 0.00 | 1.37 | 0.21 | 0.40 | Non-enriched |
| 43 | Prostate | 0.03 | 0.01 | 0.00 | 0.13 | 0.32 | 0.38 | 0.58 | 1.29 | 0.00 | 1.03 | 0.04 | 0.12 | Non-enriched |
| 44 | Skin_Not_Sun_Exposed_Suprapubic. | 0.03 | 0.01 | 0.00 | 0.23 | 0.46 | 0.48 | 0.71 | 1.43 | 0.00 | 1.03 | 0.08 | 0.16 | Non-enriched |
| 45 | Skin_Sun_Exposed_Lower_leg. | 0.03 | 0.01 | 0.00 | 0.27 | 0.46 | 0.48 | 0.67 | 1.43 | 0.00 | 0.95 | 0.09 | 0.21 | Non-enriched |
| 46 | Small_Intestine_Terminal_Ileum | 0.03 | 0.01 | 0.01 | 0.21 | 0.40 | 0.49 | 0.71 | 1.73 | 0.01 | 1.23 | 0.05 | 0.14 | Non-enriched |
| 47 | Spleen | 0.03 | 0.02 | 0.07 | 0.49 | 0.76 | 0.80 | 1.09 | 2.31 | 0.18 | 1.52 | 0.26 | 0.37 | Non-enriched |
| 48 | Stomach | 0.03 | 0.01 | 0.00 | 0.12 | 0.26 | 0.31 | 0.44 | 0.95 | 0.00 | 0.77 | 0.02 | 0.13 | Non-enriched |
| 49 | Testis | 0.03 | 0.04 | 0.00 | 0.84 | 1.24 | 1.28 | 1.76 | 3.48 | 0.00 | 2.46 | 0.14 | 0.30 | Non-enriched |
| 50 | Thyroid | 0.03 | 0.01 | 0.00 | 0.06 | 0.20 | 0.25 | 0.37 | 1.15 | 0.00 | 0.83 | 0.02 | 0.13 | Non-enriched |
| 51 | Uterus | 0.03 | 0.01 | 0.00 | 0.17 | 0.31 | 0.36 | 0.48 | 1.68 | 0.00 | 0.91 | 0.06 | 0.26 | Non-enriched |
| 52 | Vagina | 0.03 | 0.01 | 0.00 | 0.10 | 0.23 | 0.25 | 0.31 | 1.06 | 0.00 | 0.62 | 0.01 | 0.06 | Non-enriched |
| 53 | Whole_Blood | 0.03 | 0.04 | 0.24 | 0.92 | 1.38 | 1.53 | 2.01 | 3.96 | 0.47 | 3.17 | 0.26 | 0.49 | Non-enriched |

**Table S9.** Enrichment summary of repressed genes in GTEx for BMI

|  | tissue | mean_pprobes | mean_pve | Min. | 1st Qu. | Median | Mean | 3rd Qu. | Max. | CI(l) | CI(h) | ROPE | prob | group |
| --- | --- | --- | --- | --- | --- | --- | --- | --- | --- | --- | --- | --- | --- | --- |
| 1 | Adipose_Subcutaneous | 0.04 | 0.04 | 0.09 | 0.79 | 1.07 | 1.15 | 1.46 | 3.33 | 0.22 | 2.06 | 0.54 | 0.77 | Non-enriched |
| 2 | Adipose_Visceral_Omentum. | 0.04 | 0.04 | 0.22 | 0.75 | 1.01 | 1.08 | 1.34 | 3.15 | 0.27 | 1.96 | 0.56 | 0.74 | Non-enriched |
| 3 | Adrenal_Gland | 0.05 | 0.04 | 0.00 | 0.50 | 0.76 | 0.81 | 1.07 | 2.82 | 0.00 | 1.67 | 0.55 | 0.81 | Non-enriched |
| 4 | Artery_Aorta | 0.03 | 0.01 | 0.00 | 0.20 | 0.37 | 0.41 | 0.58 | 1.22 | 0.00 | 0.92 | 0.12 | 0.35 | Non-enriched |
| 5 | Artery_Coronary | 0.04 | 0.04 | 0.00 | 0.34 | 0.66 | 0.96 | 1.35 | 5.05 | 0.00 | 2.86 | 0.27 | 0.64 | Non-enriched |
| 6 | Artery_Tibial | 0.03 | 0.01 | 0.00 | 0.14 | 0.30 | 0.34 | 0.49 | 1.11 | 0.00 | 0.84 | 0.09 | 0.37 | Non-enriched |
| 7 | Bladder | 0.04 | 0.04 | 0.28 | 0.85 | 1.13 | 1.18 | 1.44 | 3.19 | 0.40 | 2.10 | 0.69 | 0.90 | Non-enriched |
| 8 | Brain_Amygdala | 0.04 | 0.02 | 0.00 | 0.19 | 0.40 | 0.53 | 0.75 | 2.57 | 0.00 | 1.44 | 0.16 | 0.45 | Non-enriched |
| 9 | Brain_Anterior_cingulate_cortex_BA24. | 0.04 | 0.02 | 0.00 | 0.21 | 0.45 | 0.58 | 0.81 | 2.67 | 0.00 | 1.55 | 0.19 | 0.47 | Non-enriched |
| 10 | Brain_Caudate_basal_ganglia. | 0.03 | 0.01 | 0.00 | 0.16 | 0.37 | 0.43 | 0.63 | 1.94 | 0.00 | 1.02 | 0.15 | 0.41 | Non-enriched |
| 11 | Brain_Cerebellar_Hemisphere | 0.04 | 0.07 | 0.51 | 1.35 | 1.91 | 1.99 | 2.48 | 4.81 | 0.61 | 3.54 | 0.25 | 0.81 | Non-enriched |
| 12 | Brain_Cerebellum | 0.03 | 0.01 | 0.00 | 0.12 | 0.28 | 0.39 | 0.53 | 2.25 | 0.00 | 1.04 | 0.08 | 0.33 | Non-enriched |
| 13 | Brain_Cortex | 0.04 | 0.02 | 0.00 | 0.24 | 0.49 | 0.64 | 0.88 | 2.74 | 0.00 | 1.80 | 0.19 | 0.45 | Non-enriched |
| 14 | Brain_Frontal_Cortex_BA9. | 0.04 | 0.02 | 0.00 | 0.24 | 0.47 | 0.59 | 0.81 | 2.74 | 0.00 | 1.60 | 0.21 | 0.51 | Non-enriched |
| 15 | Brain_Hippocampus | 0.04 | 0.02 | 0.00 | 0.30 | 0.51 | 0.59 | 0.81 | 2.57 | 0.00 | 1.38 | 0.28 | 0.59 | Non-enriched |
| 16 | Brain_Hypothalamus | 0.03 | 0.02 | 0.00 | 0.19 | 0.44 | 0.58 | 0.84 | 2.74 | 0.00 | 1.63 | 0.16 | 0.41 | Non-enriched |
| 17 | Brain_Nucleus_accumbens_basal_ganglia. | 0.04 | 0.02 | 0.00 | 0.18 | 0.46 | 0.60 | 0.87 | 2.74 | 0.00 | 1.66 | 0.18 | 0.45 | Non-enriched |
| 18 | Brain_Putamen_basal_ganglia. | 0.04 | 0.02 | 0.00 | 0.16 | 0.42 | 0.57 | 0.84 | 2.74 | 0.00 | 1.62 | 0.18 | 0.45 | Non-enriched |
| 19 | Brain_Spinal_cord_cervical_c1. | 0.04 | 0.07 | 0.58 | 1.39 | 1.88 | 1.99 | 2.50 | 4.81 | 0.70 | 3.54 | 0.27 | 0.87 | Non-enriched |
| 20 | Brain_Substantia_nigra | 0.04 | 0.02 | 0.00 | 0.45 | 0.64 | 0.70 | 0.92 | 2.57 | 0.00 | 1.39 | 0.41 | 0.64 | Non-enriched |
| 21 | Breast_Mammary_Tissue | 0.06 | 0.06 | 0.09 | 0.73 | 1.04 | 1.10 | 1.44 | 3.40 | 0.23 | 2.11 | 0.49 | 0.74 | Non-enriched |
| 22 | Cells_EBV.transformed_lymphocytes | 0.05 | 0.05 | 0.00 | 0.62 | 1.02 | 1.13 | 1.55 | 3.41 | 0.00 | 2.45 | 0.41 | 0.74 | Non-enriched |
| 23 | Cells_Transformed_fibroblasts | 0.06 | 0.25 | 1.17 | 3.16 | 3.91 | 4.11 | 4.80 | 11.26 | 1.70 | 6.85 | 0.00 | 1.00 | Enriched |
| 24 | Cervix_Ectocervix | 0.03 | 0.01 | 0.00 | 0.09 | 0.21 | 0.30 | 0.40 | 1.63 | 0.00 | 1.06 | 0.04 | 0.20 | Non-enriched |
| 25 | Cervix_Endocervix | 0.03 | 0.03 | 0.11 | 0.76 | 1.05 | 1.12 | 1.40 | 2.83 | 0.42 | 2.26 | 0.50 | 0.66 | Non-enriched |
| 26 | Colon_Sigmoid | 0.04 | 0.06 | 0.24 | 0.90 | 1.21 | 1.36 | 1.71 | 4.78 | 0.40 | 2.68 | 0.54 | 0.86 | Non-enriched |
| 27 | Colon_Transverse | 0.04 | 0.02 | 0.00 | 0.38 | 0.64 | 0.65 | 0.89 | 2.05 | 0.06 | 1.34 | 0.37 | 0.60 | Non-enriched |
| 28 | Esophagus_Gastroesophageal_Junction | 0.05 | 0.05 | 0.17 | 0.89 | 1.22 | 1.26 | 1.57 | 3.21 | 0.32 | 2.24 | 0.61 | 0.90 | Non-enriched |
| 29 | Esophagus_Mucosa | 0.03 | 0.01 | 0.00 | 0.13 | 0.30 | 0.39 | 0.54 | 1.87 | 0.00 | 1.06 | 0.07 | 0.26 | Non-enriched |
| 30 | Esophagus_Muscularis | 0.04 | 0.04 | 0.12 | 0.85 | 1.16 | 1.22 | 1.52 | 3.21 | 0.38 | 2.26 | 0.62 | 0.89 | Non-enriched |
| 31 | Fallopian_Tube | 0.04 | 0.05 | 0.17 | 0.84 | 1.19 | 1.25 | 1.58 | 3.51 | 0.35 | 2.36 | 0.45 | 0.69 | Non-enriched |
| 32 | Heart_Atrial_Appendage | 0.03 | 0.01 | 0.00 | 0.15 | 0.29 | 0.32 | 0.44 | 1.23 | 0.00 | 0.75 | 0.05 | 0.30 | Non-enriched |
| 33 | Heart_Left_Ventricle | 0.04 | 0.02 | 0.00 | 0.22 | 0.39 | 0.41 | 0.56 | 1.76 | 0.00 | 0.87 | 0.13 | 0.39 | Non-enriched |
| 34 | Kidney_Cortex | 0.04 | 0.02 | 0.00 | 0.15 | 0.36 | 0.45 | 0.66 | 2.08 | 0.00 | 1.20 | 0.17 | 0.45 | Non-enriched |
| 35 | Liver | 0.03 | 0.05 | 0.27 | 1.01 | 1.42 | 1.52 | 1.96 | 4.88 | 0.39 | 2.84 | 0.38 | 0.72 | Non-enriched |
| 36 | Lung | 0.03 | 0.01 | 0.00 | 0.18 | 0.41 | 0.40 | 0.57 | 1.42 | 0.00 | 0.87 | 0.07 | 0.22 | Non-enriched |
| 37 | Minor_Salivary_Gland | 0.04 | 0.03 | 0.07 | 0.51 | 0.68 | 0.74 | 0.92 | 1.96 | 0.10 | 1.36 | 0.59 | 0.83 | Non-enriched |
| 38 | Muscle_Skeletal | 0.03 | 0.01 | 0.00 | 0.13 | 0.30 | 0.33 | 0.46 | 1.61 | 0.00 | 0.78 | 0.04 | 0.22 | Non-enriched |
| 39 | Nerve_Tibial | 0.03 | 0.01 | 0.00 | 0.11 | 0.20 | 0.27 | 0.36 | 1.93 | 0.00 | 0.79 | 0.04 | 0.23 | Non-enriched |
| 40 | Ovary | 0.03 | 0.01 | 0.00 | 0.14 | 0.34 | 0.40 | 0.56 | 2.39 | 0.00 | 1.08 | 0.09 | 0.30 | Non-enriched |
| 41 | Pancreas | 0.03 | 0.01 | 0.00 | 0.11 | 0.27 | 0.34 | 0.50 | 1.60 | 0.00 | 0.88 | 0.06 | 0.25 | Non-enriched |
| 42 | Pituitary | 0.04 | 0.03 | 0.00 | 0.32 | 0.61 | 0.68 | 0.94 | 2.78 | 0.00 | 1.62 | 0.24 | 0.45 | Non-enriched |
| 43 | Prostate | 0.03 | 0.02 | 0.00 | 0.27 | 0.55 | 0.72 | 1.05 | 3.85 | 0.00 | 1.98 | 0.21 | 0.50 | Non-enriched |
| 44 | Skin_Not_Sun_Exposed_Suprapubic. | 0.03 | 0.03 | 0.00 | 0.44 | 1.01 | 1.06 | 1.47 | 4.46 | 0.00 | 2.53 | 0.17 | 0.37 | Non-enriched |
| 45 | Skin_Sun_Exposed_Lower_leg. | 0.03 | 0.03 | 0.00 | 0.49 | 1.06 | 1.16 | 1.59 | 5.72 | 0.00 | 2.65 | 0.16 | 0.36 | Non-enriched |
| 46 | Small_Intestine_Terminal_Ileum | 0.04 | 0.01 | 0.00 | 0.14 | 0.30 | 0.42 | 0.58 | 2.60 | 0.00 | 1.24 | 0.10 | 0.39 | Non-enriched |
| 47 | Spleen | 0.04 | 0.03 | 0.00 | 0.47 | 0.79 | 0.84 | 1.11 | 3.73 | 0.00 | 1.75 | 0.49 | 0.78 | Non-enriched |
| 48 | Stomach | 0.04 | 0.03 | 0.04 | 0.44 | 0.68 | 0.72 | 0.95 | 2.28 | 0.05 | 1.39 | 0.44 | 0.65 | Non-enriched |
| 49 | Testis | 0.03 | 0.02 | 0.00 | 0.25 | 0.48 | 0.54 | 0.74 | 2.10 | 0.00 | 1.27 | 0.19 | 0.41 | Non-enriched |
| 50 | Thyroid | 0.07 | 0.06 | 0.00 | 0.19 | 0.45 | 0.60 | 0.80 | 3.44 | 0.00 | 1.71 | 0.21 | 0.54 | Non-enriched |
| 51 | Uterus | 0.04 | 0.04 | 0.27 | 0.90 | 1.22 | 1.26 | 1.55 | 3.28 | 0.49 | 2.27 | 0.65 | 0.93 | Non-enriched |
| 52 | Vagina | 0.03 | 0.02 | 0.00 | 0.27 | 0.55 | 0.69 | 0.98 | 3.85 | 0.00 | 1.69 | 0.15 | 0.34 | Non-enriched |
| 53 | Whole_Blood | 0.03 | 0.01 | 0.00 | 0.13 | 0.28 | 0.34 | 0.50 | 1.41 | 0.00 | 0.87 | 0.05 | 0.21 | Non-enriched |

**Table S10.** Enrichment summary of over-expressed genes in GTEx for smoking

|  | tissue | mean_pprobes | mean_pve | Min. | 1st Qu. | Median | Mean | 3rd Qu. | Max. | CI(l) | CI(h) | ROPE | prob | group |
| --- | --- | --- | --- | --- | --- | --- | --- | --- | --- | --- | --- | --- | --- | --- |
| 1 | Adipose_Subcutaneous | 0.08 | 0.02 | 0.00 | 0.08 | 0.17 | 0.22 | 0.29 | 1.09 | 0.00 | 0.77 | 0.01 | 0.07 | Non-enriched |
| 2 | Adipose_Visceral_Omentum. | 0.07 | 0.02 | 0.00 | 0.11 | 0.18 | 0.23 | 0.33 | 0.77 | 0.01 | 0.58 | 0.01 | 0.08 | Non-enriched |
| 3 | Adrenal_Gland | 0.07 | 0.01 | 0.00 | 0.08 | 0.12 | 0.18 | 0.26 | 0.75 | 0.00 | 0.62 | 0.01 | 0.07 | Non-enriched |
| 4 | Artery_Aorta | 0.06 | 0.01 | 0.00 | 0.05 | 0.13 | 0.17 | 0.24 | 0.70 | 0.00 | 0.56 | 0.00 | 0.11 | Non-enriched |
| 5 | Artery_Coronary | 0.07 | 0.03 | 0.00 | 0.28 | 0.38 | 0.41 | 0.51 | 1.28 | 0.11 | 0.76 | 0.10 | 0.38 | Non-enriched |
| 6 | Artery_Tibial | 0.07 | 0.03 | 0.00 | 0.28 | 0.38 | 0.41 | 0.50 | 1.04 | 0.11 | 0.81 | 0.11 | 0.44 | Non-enriched |
| 7 | Bladder | 0.07 | 0.02 | 0.00 | 0.11 | 0.18 | 0.21 | 0.28 | 0.59 | 0.00 | 0.58 | 0.00 | 0.07 | Non-enriched |
| 8 | Brain_Amygdala | 0.07 | 0.01 | 0.00 | 0.04 | 0.13 | 0.16 | 0.23 | 0.65 | 0.00 | 0.44 | 0.00 | 0.14 | Non-enriched |
| 9 | Brain_Anterior_cingulate_cortex_BA24. | 0.07 | 0.01 | 0.00 | 0.04 | 0.14 | 0.16 | 0.22 | 0.65 | 0.00 | 0.48 | 0.00 | 0.14 | Non-enriched |
| 10 | Brain_Caudate_basal_ganglia. | 0.07 | 0.01 | 0.00 | 0.03 | 0.10 | 0.16 | 0.24 | 0.65 | 0.00 | 0.50 | 0.00 | 0.09 | Non-enriched |
| 11 | Brain_Cerebellar_Hemisphere | 0.10 | 0.04 | 0.00 | 0.28 | 0.38 | 0.44 | 0.54 | 1.70 | 0.08 | 0.92 | 0.12 | 0.39 | Non-enriched |
| 12 | Brain_Cerebellum | 0.10 | 0.04 | 0.00 | 0.26 | 0.36 | 0.40 | 0.50 | 1.16 | 0.10 | 0.84 | 0.09 | 0.36 | Non-enriched |
| 13 | Brain_Cortex | 0.07 | 0.02 | 0.00 | 0.16 | 0.23 | 0.26 | 0.32 | 0.85 | 0.02 | 0.61 | 0.02 | 0.24 | Non-enriched |
| 14 | Brain_Frontal_Cortex_BA9. | 0.07 | 0.02 | 0.00 | 0.16 | 0.23 | 0.27 | 0.32 | 0.90 | 0.02 | 0.61 | 0.02 | 0.24 | Non-enriched |
| 15 | Brain_Hippocampus | 0.07 | 0.01 | 0.00 | 0.04 | 0.13 | 0.16 | 0.23 | 0.65 | 0.00 | 0.47 | 0.00 | 0.15 | Non-enriched |
| 16 | Brain_Hypothalamus | 0.07 | 0.02 | 0.00 | 0.14 | 0.21 | 0.23 | 0.29 | 0.79 | 0.00 | 0.53 | 0.01 | 0.17 | Non-enriched |
| 17 | Brain_Nucleus_accumbens_basal_ganglia. | 0.07 | 0.01 | 0.00 | 0.04 | 0.09 | 0.14 | 0.21 | 0.65 | 0.00 | 0.48 | 0.00 | 0.13 | Non-enriched |
| 18 | Brain_Putamen_basal_ganglia. | 0.07 | 0.01 | 0.00 | 0.03 | 0.10 | 0.15 | 0.23 | 0.65 | 0.00 | 0.44 | 0.00 | 0.09 | Non-enriched |
| 19 | Brain_Spinal_cord_cervical_c1. | 0.13 | 0.14 | 0.24 | 0.81 | 1.05 | 1.11 | 1.35 | 2.64 | 0.46 | 2.01 | 0.79 | 1.00 | Non-enriched |
| 20 | Brain_Substantia_nigra | 0.13 | 0.14 | 0.18 | 0.81 | 1.07 | 1.13 | 1.36 | 3.17 | 0.44 | 2.01 | 0.78 | 1.00 | Non-enriched |
| 21 | Breast_Mammary_Tissue | 0.08 | 0.02 | 0.00 | 0.07 | 0.19 | 0.22 | 0.30 | 0.98 | 0.00 | 0.58 | 0.00 | 0.05 | Non-enriched |
| 22 | Cells_EBV.transformed_lymphocytes | 0.06 | 0.01 | 0.00 | 0.09 | 0.18 | 0.22 | 0.28 | 0.72 | 0.00 | 0.65 | 0.01 | 0.05 | Non-enriched |
| 23 | Cells_Transformed_fibroblasts | 0.08 | 0.01 | 0.00 | 0.06 | 0.12 | 0.16 | 0.24 | 0.60 | 0.00 | 0.59 | 0.00 | 0.08 | Non-enriched |
| 24 | Cervix_Ectocervix | 0.07 | 0.01 | 0.00 | 0.10 | 0.16 | 0.19 | 0.30 | 0.43 | 0.00 | 0.42 | 0.00 | 0.05 | Non-enriched |
| 25 | Cervix_Endocervix | 0.06 | 0.02 | 0.00 | 0.13 | 0.32 | 0.45 | 0.67 | 2.08 | 0.00 | 1.49 | 0.03 | 0.10 | Non-enriched |
| 26 | Colon_Sigmoid | 0.20 | 0.10 | 0.00 | 0.08 | 0.15 | 0.31 | 0.28 | 2.62 | 0.00 | 1.48 | 0.02 | 0.18 | Non-enriched |
| 27 | Colon_Transverse | 0.09 | 0.02 | 0.00 | 0.04 | 0.15 | 0.20 | 0.25 | 1.01 | 0.00 | 0.80 | 0.01 | 0.06 | Non-enriched |
| 28 | Esophagus_Gastroesophageal_Junction | 0.07 | 0.01 | 0.00 | 0.04 | 0.13 | 0.16 | 0.23 | 1.45 | 0.00 | 0.50 | 0.01 | 0.14 | Non-enriched |
| 29 | Esophagus_Mucosa | 0.06 | 0.01 | 0.00 | 0.06 | 0.13 | 0.16 | 0.22 | 0.63 | 0.00 | 0.49 | 0.00 | 0.04 | Non-enriched |
| 30 | Esophagus_Muscularis | 0.07 | 0.01 | 0.00 | 0.04 | 0.14 | 0.18 | 0.25 | 1.45 | 0.00 | 0.64 | 0.01 | 0.16 | Non-enriched |
| 31 | Fallopian_Tube | 0.06 | 0.01 | 0.00 | 0.06 | 0.14 | 0.18 | 0.25 | 0.57 | 0.00 | 0.53 | 0.00 | 0.04 | Non-enriched |
| 32 | Heart_Atrial_Appendage | 0.09 | 0.02 | 0.00 | 0.06 | 0.15 | 0.17 | 0.24 | 0.52 | 0.00 | 0.45 | 0.00 | 0.09 | Non-enriched |
| 33 | Heart_Left_Ventricle | 0.09 | 0.02 | 0.00 | 0.07 | 0.16 | 0.18 | 0.25 | 0.51 | 0.00 | 0.47 | 0.00 | 0.09 | Non-enriched |
| 34 | Kidney_Cortex | 0.06 | 0.01 | 0.00 | 0.06 | 0.11 | 0.15 | 0.19 | 0.68 | 0.00 | 0.68 | 0.00 | 0.03 | Non-enriched |
| 35 | Liver | 0.06 | 0.01 | 0.01 | 0.03 | 0.13 | 0.17 | 0.23 | 0.54 | 0.00 | 0.01 | 0.03 | 0.02 | Non-enriched |
| 36 | Lung | 0.07 | 0.04 | 0.15 | 0.53 | 0.70 | 0.75 | 0.96 | 1.78 | 0.18 | 1.38 | 0.17 | 0.22 | Non-enriched |
| 37 | Minor_Salivary_Gland | 0.06 | 0.01 | 0.00 | 0.04 | 0.18 | 0.23 | 0.37 | 0.69 | 0.00 | 0.65 | 0.01 | 0.05 | Non-enriched |
| 38 | Muscle_Skeletal | 0.08 | 0.02 | 0.00 | 0.04 | 0.13 | 0.20 | 0.31 | 1.03 | 0.00 | 0.60 | 0.01 | 0.12 | Non-enriched |
| 39 | Nerve_Tibial | 0.14 | 0.04 | 0.00 | 0.18 | 0.25 | 0.27 | 0.36 | 1.05 | 0.01 | 0.51 | 0.03 | 0.49 | Non-enriched |
| 40 | Ovary | 0.07 | 0.01 | 0.00 | 0.06 | 0.12 | 0.14 | 0.21 | 0.39 | 0.00 | 0.36 | 0.00 | 0.07 | Non-enriched |
| 41 | Pancreas | 0.06 | 0.01 | 0.00 | 0.02 | 0.09 | 0.11 | 0.12 | 0.47 | 0.00 | 0.47 | 0.00 | 0.02 | Non-enriched |
| 42 | Pituitary | 0.06 | 0.01 | 0.00 | 0.04 | 0.12 | 0.18 | 0.24 | 0.76 | 0.00 | 0.65 | 0.01 | 0.09 | Non-enriched |
| 43 | Prostate | 0.08 | 0.02 | 0.00 | 0.04 | 0.09 | 0.21 | 0.36 | 0.67 | 0.00 | 0.67 | 0.01 | 0.03 | Non-enriched |
| 44 | Skin_Not_Sun_Exposed_Suprapubic. | 0.06 | 0.03 | 0.01 | 0.35 | 0.52 | 0.52 | 0.69 | 1.31 | 0.01 | 1.02 | 0.05 | 0.10 | Non-enriched |
| 45 | Skin_Sun_Exposed_Lower_leg. | 0.07 | 0.03 | 0.01 | 0.31 | 0.44 | 0.48 | 0.64 | 1.31 | 0.01 | 1.05 | 0.04 | 0.10 | Non-enriched |
| 46 | Small_Intestine_Terminal_Ileum | 0.06 | 0.02 | 0.00 | 0.08 | 0.20 | 0.27 | 0.43 | 1.01 | 0.00 | 0.80 | 0.01 | 0.07 | Non-enriched |
| 47 | Spleen | 0.06 | 0.02 | 0.00 | 0.17 | 0.29 | 0.31 | 0.43 | 1.00 | 0.00 | 0.65 | 0.01 | 0.06 | Non-enriched |
| 48 | Stomach | 0.06 | 0.01 | 0.00 | 0.07 | 0.13 | 0.18 | 0.25 | 0.64 | 0.00 | 0.56 | 0.01 | 0.06 | Non-enriched |
| 49 | Testis | 0.08 | 0.01 | 0.00 | 0.03 | 0.06 | 0.07 | 0.09 | 0.35 | 0.00 | 0.00 | 0.03 | 0.01 | Non-enriched |
| 50 | Thyroid | 0.08 | 0.02 | 0.00 | 0.03 | 0.14 | 0.24 | 0.40 | 1.21 | 0.00 | 0.81 | 0.01 | 0.08 | Non-enriched |
| 51 | Uterus | 0.07 | 0.03 | 0.00 | 0.19 | 0.36 | 0.44 | 0.57 | 1.94 | 0.00 | 1.23 | 0.05 | 0.15 | Non-enriched |
| 52 | Vagina | 0.06 | 0.01 | 0.00 | 0.02 | 0.08 | 0.13 | 0.22 | 0.63 | 0.00 | 0.63 | 0.00 | 0.03 | Non-enriched |
| 53 | Whole_Blood | 0.08 | 0.01 | 0.00 | 0.04 | 0.13 | 0.18 | 0.28 | 0.86 | 0.00 | 0.51 | 0.00 | 0.04 | Non-enriched |

**Table S11.** Enrichment summary of over-expressed genes in GTEx for smoking

|  | tissue | mean_pprobes | mean_pve | Min. | 1st Qu. | Median | Mean | 3rd Qu. | Max. | CI(l) | CI(h) | ROPE | prob | group |
| --- | --- | --- | --- | --- | --- | --- | --- | --- | --- | --- | --- | --- | --- | --- |
| 1 | Adipose_Subcutaneous | 0.09 | 0.02 | 0.00 | 0.07 | 0.17 | 0.22 | 0.32 | 1.05 | 0.00 | 0.54 | 0.01 | 0.10 | Non-enriched |
| 2 | Adipose_Visceral_Omentum. | 0.07 | 0.01 | 0.00 | 0.07 | 0.12 | 0.18 | 0.25 | 0.83 | 0.00 | 0.50 | 0.00 | 0.10 | Non-enriched |
| 3 | Adrenal_Gland | 0.07 | 0.02 | 0.00 | 0.16 | 0.25 | 0.30 | 0.40 | 0.94 | 0.01 | 0.77 | 0.03 | 0.25 | Non-enriched |
| 4 | Artery_Aorta | 0.07 | 0.02 | 0.00 | 0.11 | 0.20 | 0.25 | 0.32 | 0.99 | 0.00 | 0.72 | 0.04 | 0.30 | Non-enriched |
| 5 | Artery_Coronary | 0.07 | 0.01 | 0.00 | 0.10 | 0.16 | 0.19 | 0.24 | 1.05 | 0.01 | 0.42 | 0.01 | 0.32 | Non-enriched |
| 6 | Artery_Tibial | 0.07 | 0.01 | 0.00 | 0.10 | 0.17 | 0.21 | 0.29 | 0.77 | 0.00 | 0.54 | 0.02 | 0.30 | Non-enriched |
| 7 | Bladder | 0.07 | 0.02 | 0.00 | 0.17 | 0.33 | 0.35 | 0.48 | 1.05 | 0.00 | 0.83 | 0.05 | 0.23 | Non-enriched |
| 8 | Brain_Amygdala | 0.07 | 0.01 | 0.00 | 0.05 | 0.13 | 0.19 | 0.25 | 1.21 | 0.00 | 0.63 | 0.01 | 0.15 | Non-enriched |
| 9 | Brain_Anterior_cingulate_cortex_BA24. | 0.08 | 0.01 | 0.00 | 0.04 | 0.11 | 0.18 | 0.24 | 1.21 | 0.00 | 0.49 | 0.01 | 0.14 | Non-enriched |
| 10 | Brain_Caudate_basal_ganglia. | 0.07 | 0.01 | 0.00 | 0.05 | 0.12 | 0.18 | 0.25 | 0.87 | 0.00 | 0.55 | 0.01 | 0.13 | Non-enriched |
| 11 | Brain_Cerebellar_Hemisphere | 0.07 | 0.02 | 0.00 | 0.08 | 0.20 | 0.26 | 0.36 | 1.64 | 0.00 | 0.85 | 0.02 | 0.16 | Non-enriched |
| 12 | Brain_Cerebellum | 0.07 | 0.02 | 0.00 | 0.10 | 0.22 | 0.28 | 0.39 | 1.64 | 0.00 | 0.81 | 0.02 | 0.18 | Non-enriched |
| 13 | Brain_Cortex | 0.07 | 0.01 | 0.00 | 0.05 | 0.12 | 0.16 | 0.23 | 0.71 | 0.00 | 0.42 | 0.00 | 0.12 | Non-enriched |
| 14 | Brain_Frontal_Cortex_BA9. | 0.08 | 0.02 | 0.00 | 0.06 | 0.14 | 0.19 | 0.26 | 1.21 | 0.00 | 0.53 | 0.01 | 0.15 | Non-enriched |
| 15 | Brain_Hippocampus | 0.07 | 0.02 | 0.00 | 0.06 | 0.18 | 0.25 | 0.35 | 1.64 | 0.00 | 0.75 | 0.02 | 0.19 | Non-enriched |
| 16 | Brain_Hypothalamus | 0.07 | 0.02 | 0.00 | 0.06 | 0.21 | 0.28 | 0.39 | 1.64 | 0.00 | 0.94 | 0.02 | 0.15 | Non-enriched |
| 17 | Brain_Nucleus_accumbens_basal_ganglia. | 0.07 | 0.02 | 0.00 | 0.06 | 0.17 | 0.24 | 0.33 | 1.64 | 0.00 | 0.81 | 0.02 | 0.17 | Non-enriched |
| 18 | Brain_Putamen_basal_ganglia. | 0.08 | 0.01 | 0.00 | 0.05 | 0.10 | 0.16 | 0.22 | 0.81 | 0.00 | 0.46 | 0.00 | 0.13 | Non-enriched |
| 19 | Brain_Spinal_cord_cervical_c1. | 0.07 | 0.02 | 0.00 | 0.06 | 0.20 | 0.26 | 0.36 | 1.37 | 0.00 | 0.86 | 0.02 | 0.18 | Non-enriched |
| 20 | Brain_Substantia_nigra | 0.07 | 0.02 | 0.00 | 0.09 | 0.19 | 0.24 | 0.34 | 1.30 | 0.00 | 0.61 | 0.02 | 0.19 | Non-enriched |
| 21 | Breast_Mammary_Tissue | 0.17 | 0.08 | 0.00 | 0.13 | 0.24 | 0.36 | 0.42 | 2.62 | 0.00 | 1.24 | 0.04 | 0.24 | Non-enriched |
| 22 | Cells_EBV.transformed_lymphocytes | 0.07 | 0.01 | 0.00 | 0.05 | 0.11 | 0.16 | 0.23 | 0.72 | 0.00 | 0.45 | 0.00 | 0.16 | Non-enriched |
| 23 | Cells_Transformed_fibroblasts | 0.07 | 0.02 | 0.00 | 0.16 | 0.23 | 0.27 | 0.37 | 1.07 | 0.00 | 0.57 | 0.02 | 0.18 | Non-enriched |
| 24 | Cervix_Ectocervix | 0.06 | 0.01 | 0.00 | 0.07 | 0.16 | 0.19 | 0.31 | 0.56 | 0.00 | 0.50 | 0.00 | 0.08 | Non-enriched |
| 25 | Cervix_Endocervix | 0.07 | 0.02 | 0.00 | 0.13 | 0.26 | 0.30 | 0.42 | 1.44 | 0.00 | 0.73 | 0.02 | 0.14 | Non-enriched |
| 26 | Colon_Sigmoid | 0.07 | 0.01 | 0.00 | 0.09 | 0.14 | 0.17 | 0.21 | 0.75 | 0.00 | 0.41 | 0.01 | 0.23 | Non-enriched |
| 27 | Colon_Transverse | 0.07 | 0.01 | 0.00 | 0.05 | 0.11 | 0.16 | 0.21 | 0.90 | 0.00 | 0.50 | 0.00 | 0.11 | Non-enriched |
| 28 | Esophagus_Gastroesophageal_Junction | 0.09 | 0.02 | 0.00 | 0.04 | 0.11 | 0.17 | 0.28 | 1.05 | 0.00 | 0.50 | 0.00 | 0.10 | Non-enriched |
| 29 | Esophagus_Mucosa | 0.07 | 0.01 | 0.00 | 0.06 | 0.14 | 0.19 | 0.28 | 0.98 | 0.00 | 0.57 | 0.00 | 0.11 | Non-enriched |
| 30 | Esophagus_Muscularis | 0.08 | 0.01 | 0.00 | 0.09 | 0.15 | 0.18 | 0.22 | 1.05 | 0.00 | 0.47 | 0.01 | 0.24 | Non-enriched |
| 31 | Fallopian_Tube | 0.08 | 0.02 | 0.00 | 0.16 | 0.28 | 0.33 | 0.42 | 1.17 | 0.00 | 0.82 | 0.03 | 0.16 | Non-enriched |
| 32 | Heart_Atrial_Appendage | 0.07 | 0.01 | 0.00 | 0.06 | 0.16 | 0.21 | 0.31 | 1.47 | 0.00 | 0.63 | 0.01 | 0.11 | Non-enriched |
| 33 | Heart_Left_Ventricle | 0.07 | 0.01 | 0.00 | 0.03 | 0.08 | 0.12 | 0.16 | 1.18 | 0.00 | 0.39 | 0.00 | 0.09 | Non-enriched |
| 34 | Kidney_Cortex | 0.08 | 0.03 | 0.00 | 0.27 | 0.39 | 0.40 | 0.52 | 1.37 | 0.06 | 0.74 | 0.13 | 0.48 | Non-enriched |
| 35 | Liver | 0.07 | 0.01 | 0.00 | 0.06 | 0.15 | 0.23 | 0.35 | 1.35 | 0.00 | 0.78 | 0.01 | 0.12 | Non-enriched |
| 36 | Lung | 0.11 | 0.02 | 0.00 | 0.05 | 0.14 | 0.18 | 0.27 | 0.73 | 0.00 | 0.46 | 0.00 | 0.09 | Non-enriched |
| 37 | Minor_Salivary_Gland | 0.06 | 0.01 | 0.00 | 0.06 | 0.14 | 0.19 | 0.28 | 0.92 | 0.00 | 0.59 | 0.01 | 0.14 | Non-enriched |
| 38 | Muscle_Skeletal | 0.07 | 0.01 | 0.00 | 0.03 | 0.12 | 0.17 | 0.19 | 1.18 | 0.00 | 0.61 | 0.00 | 0.06 | Non-enriched |
| 39 | Nerve_Tibial | 0.07 | 0.01 | 0.00 | 0.06 | 0.13 | 0.16 | 0.22 | 0.70 | 0.00 | 0.52 | 0.00 | 0.09 | Non-enriched |
| 40 | Ovary | 0.06 | 0.01 | 0.00 | 0.07 | 0.18 | 0.22 | 0.32 | 1.16 | 0.00 | 0.60 | 0.01 | 0.10 | Non-enriched |
| 41 | Pancreas | 0.07 | 0.02 | 0.00 | 0.06 | 0.18 | 0.26 | 0.38 | 1.37 | 0.00 | 0.93 | 0.02 | 0.14 | Non-enriched |
| 42 | Pituitary | 0.07 | 0.01 | 0.00 | 0.03 | 0.11 | 0.16 | 0.25 | 0.71 | 0.00 | 0.53 | 0.01 | 0.10 | Non-enriched |
| 43 | Prostate | 0.07 | 0.02 | 0.00 | 0.09 | 0.22 | 0.26 | 0.36 | 1.21 | 0.00 | 0.70 | 0.02 | 0.17 | Non-enriched |
| 44 | Skin_Not_Sun_Exposed_Suprapubic. | 0.07 | 0.01 | 0.00 | 0.05 | 0.15 | 0.19 | 0.28 | 0.83 | 0.00 | 0.53 | 0.00 | 0.09 | Non-enriched |
| 45 | Skin_Sun_Exposed_Lower_leg. | 0.07 | 0.01 | 0.00 | 0.04 | 0.13 | 0.18 | 0.29 | 0.83 | 0.00 | 0.53 | 0.00 | 0.08 | Non-enriched |
| 46 | Small_Intestine_Terminal_Ileum | 0.07 | 0.01 | 0.00 | 0.06 | 0.11 | 0.15 | 0.20 | 0.70 | 0.00 | 0.41 | 0.00 | 0.13 | Non-enriched |
| 47 | Spleen | 0.06 | 0.02 | 0.00 | 0.14 | 0.24 | 0.27 | 0.37 | 0.90 | 0.00 | 0.58 | 0.01 | 0.18 | Non-enriched |
| 48 | Stomach | 0.07 | 0.01 | 0.00 | 0.08 | 0.17 | 0.23 | 0.33 | 0.89 | 0.00 | 0.73 | 0.02 | 0.13 | Non-enriched |
| 49 | Testis | 0.07 | 0.01 | 0.00 | 0.05 | 0.14 | 0.18 | 0.24 | 1.16 | 0.00 | 0.53 | 0.01 | 0.11 | Non-enriched |
| 50 | Thyroid | 0.18 | 0.06 | 0.00 | 0.17 | 0.24 | 0.30 | 0.35 | 2.17 | 0.00 | 0.65 | 0.04 | 0.55 | Non-enriched |
| 51 | Uterus | 0.06 | 0.02 | 0.00 | 0.11 | 0.22 | 0.27 | 0.37 | 0.99 | 0.00 | 0.68 | 0.02 | 0.11 | Non-enriched |
| 52 | Vagina | 0.06 | 0.01 | 0.00 | 0.07 | 0.15 | 0.21 | 0.29 | 0.85 | 0.00 | 0.71 | 0.01 | 0.10 | Non-enriched |
| 53 | Whole_Blood | 0.07 | 0.01 | 0.00 | 0.05 | 0.10 | 0.13 | 0.18 | 0.65 | 0.00 | 0.35 | 0.00 | 0.07 | Non-enriched |

**Table S12.**
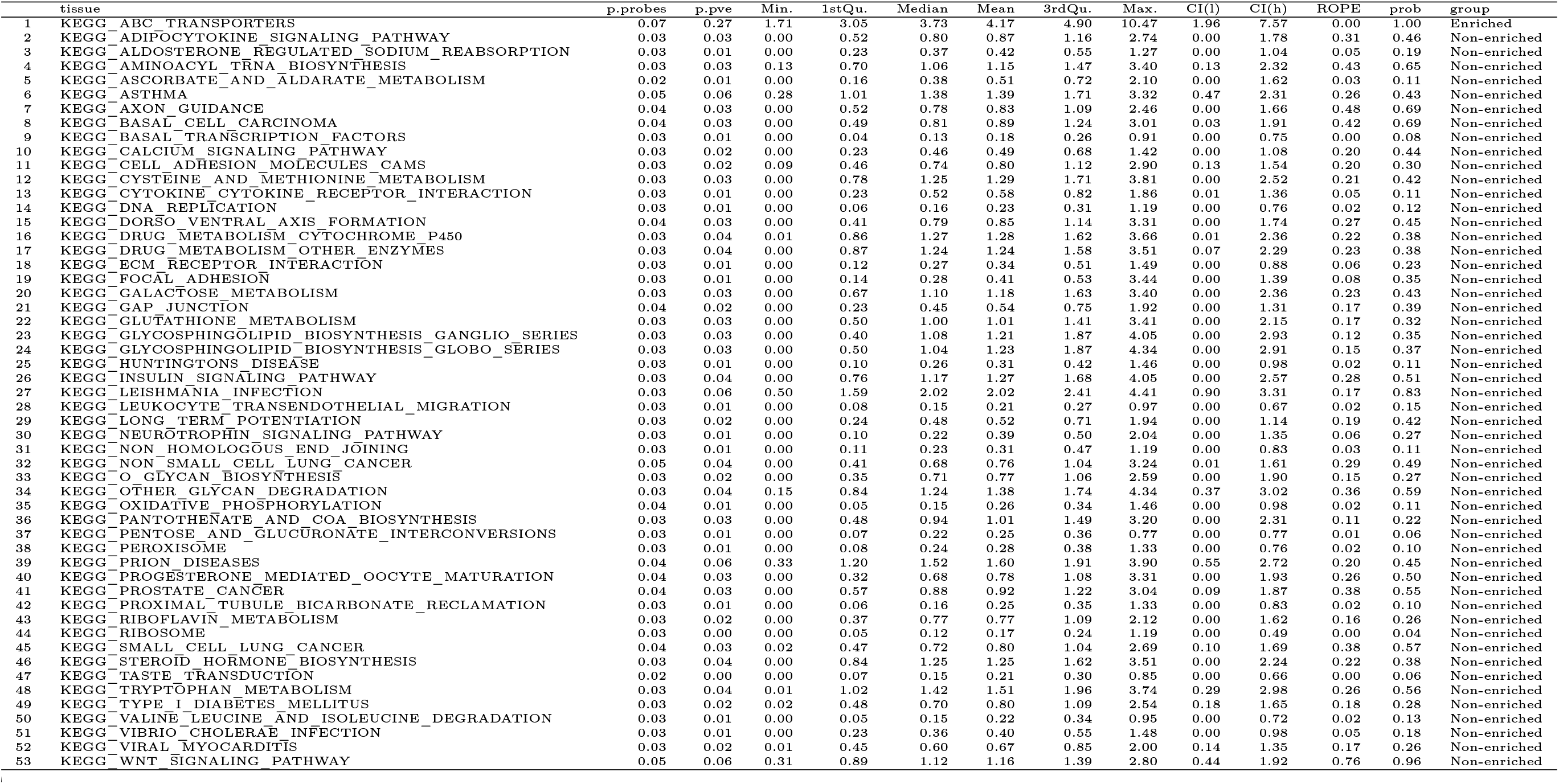
Enrichment summary of over-expressed genes in DEPICT for BMI

**Table S13.**
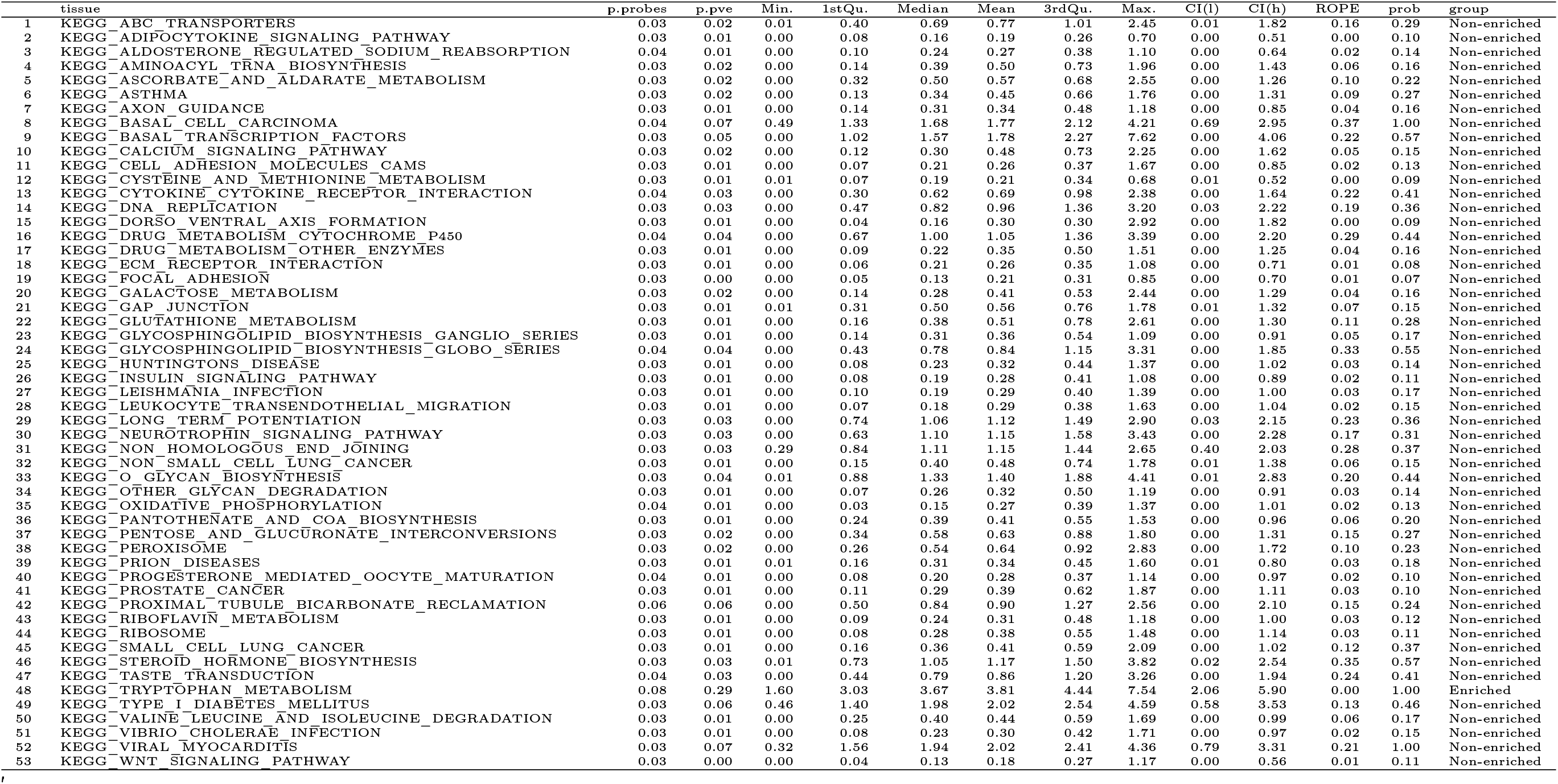
Enrichment summary of repressed genes in DEPICT for BMI

**Table S14.**
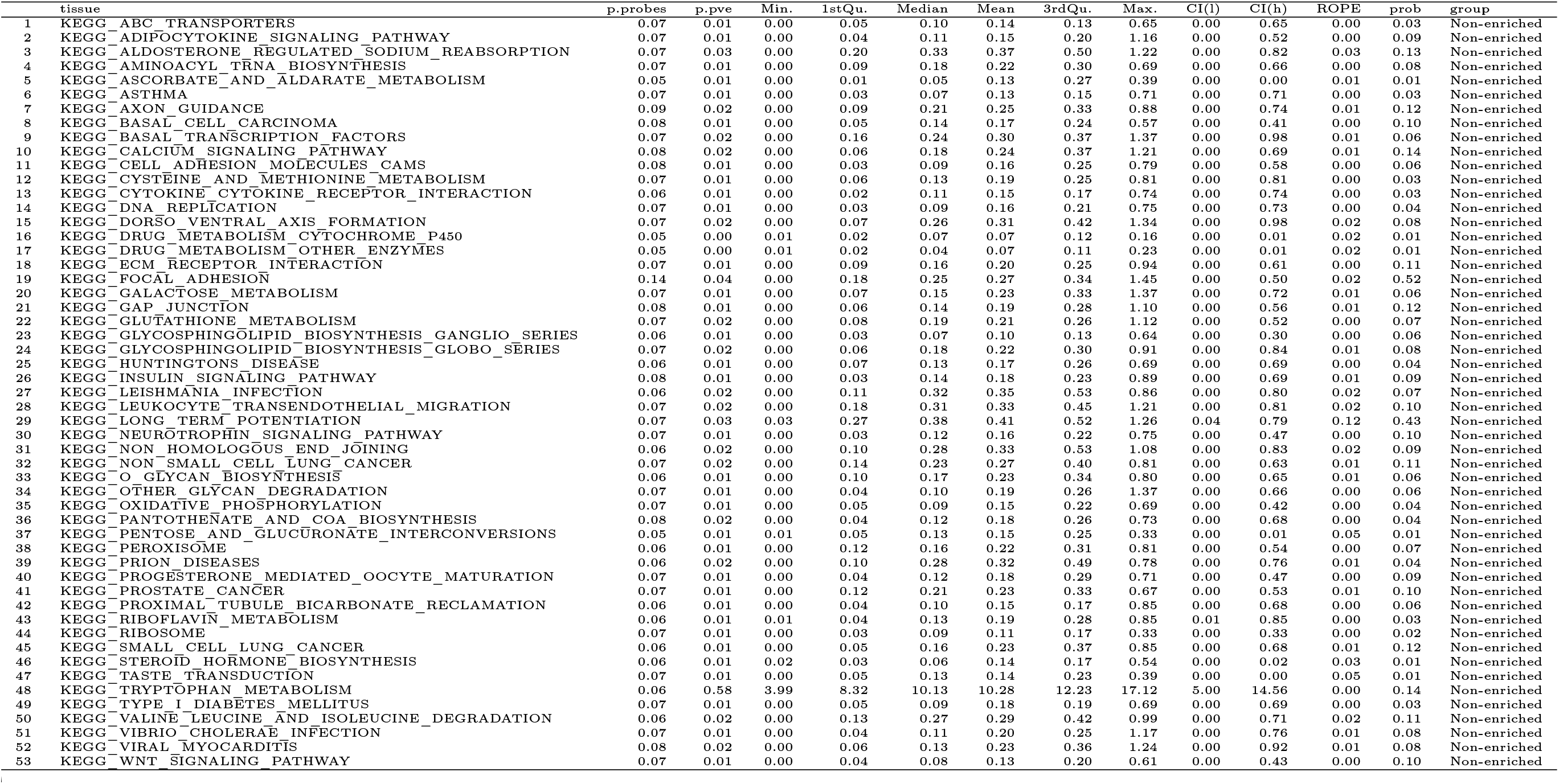
Enrichment summary of over-expressed genes in DEPICT for smoking

**Table S15.**
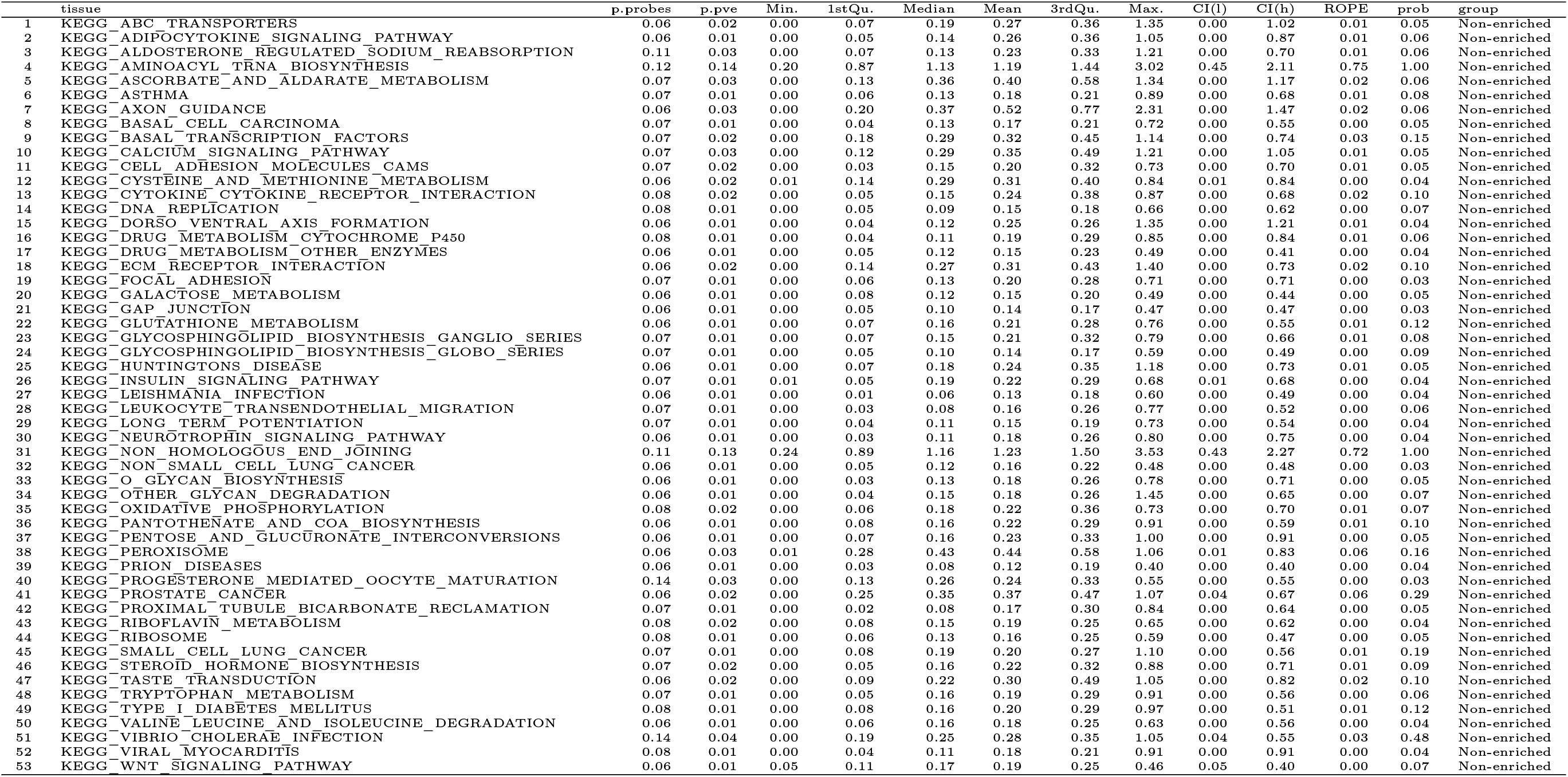
Enrichment summary of repressed genes in DEPICT for smoking

## References

1. Rudolf Jaenisch and Adrian Bird. Epigenetic regulation of gene expression: how the genome integrates intrinsic and environmental signals. Nature genetics, 33:245, 2003.

2. Yehudit Bergman and Howard Cedar. Dna methylation dynamics in health and disease. Nature structural & molecular biology, 20(20):274, 2013.

3. Tina Rönn and Charlotte Ling. Dna methylation as a diagnostic and therapeutic target in the battle against type 2 diabetes. Epigenomics, 7(7):451–460, 2015.

4. Xiaojing Yang, Fides Lay, Han Han, and Peter A Jones. Targeting dna methylation for epigenetic therapy. Trends in pharmacological sciences, 31(31):536–546, 2010.

5. Caroline L Relton and George Davey Smith. Epigenetic epidemiology of common complex disease: prospects for prediction, prevention, and treatment. PLoS medicine, 7(7):e1000356, 2010.

6. James M Flanagan. Epigenome-wide association studies (ewas): past, present, and future. In Cancer Epigenetics, pages 51–63. Springer, 2015.

7. Maarten van Iterson, Erik W van Zwet, and Bastiaan T Heijmans. Controlling bias and inflation in epigenome-and transcriptome-wide association studies using the empirical null distribution. Genome biology, 18(18):19, 2017.

8. Ewan Birney, George Davey Smith, and John M Greally. Epigenome-wide association studies and the interpretation of disease-omics. PLoS genetics, 12(12):e1006105, 2016.

9. Jeffrey T Leek and John D Storey. Capturing heterogeneity in gene expression studies by surrogate variable analysis. PLoS genetics, 3(3):e161, 2007.

10. Johann A Gagnon-Bartsch and Terence P Speed. Using control genes to correct for unwanted variation in microarray data. Biostatistics, 13(13):539–552, 2012.

11. Eugene Andres Houseman, John Molitor, and Carmen J. Marsit. Reference-free cell mixture adjustments in analysis of dna methylation data. Bioinformatics, 30(30):1431–1439, 2014.

12. James Zou, Christoph Lippert, David Heckerman, Martin Aryee, and Jennifer Listgarten. Epigenome-wide association studies without the need for cell-type composition. Nature methods, 11(11):309, 2014.

13. Elior Rahmani, Noah Zaitlen, Yael Baran, Celeste Eng, Donglei Hu, Joshua Galanter, Sam Oh, Sparse pca corrects for cell type heterogeneity in epigenome-wide association studies. Nature methods, 13(13):443, 2016.

14. Zhang Futao, Chen Wenhan, Zhu Zhihong, Zhang Qian, Deary Ian J., Wray Naomi R., Visscher Peter M., McRae Allan F., and Yang Jian. Osca: a tool for omic-data-based complex trait analysis.

15. Blair H Smith, Archie Campbell, Pamela Linksted, Bridie Fitzpatrick, Cathy Jackson, Shona M Kerr, Ian J Deary, Donald J MacIntyre, Harry Campbell, Mark McGilchrist, et al. Cohort profile: Generation scotland: Scottish family health study (gs: Sfhs). the study, its participants and their potential for genetic research on health and illness. International journal of epidemiology, 42(42):689–700, 2012.

16. D Habier, R.L Fernando, and J. C. M. Dekkers. The impact of genetic relationship information on genome-assisted breeding values. Genetics, 177:2389–2397, 2007.

17. Kevin Caye, Basile Jumentier, and Olivier Francois. Lfmm 2.0: Latent factor models for confounder adjustment in genome and epigenome-wide association studies. bioRxiv, 2018.

18. Jerome Friedman, Trevor Hastie, and Robert Tibshirani. Regularization paths for generalized linear models via coordinate descent. Journal of Statistical Software, 33(33):1–22, 2010.

19. Srikant Ambatipudi, Cyrille Cuenin, Hector Hernandez-Vargas, Akram Ghantous, Florence Le Calvez-Kelm, Rudolf Kaaks, Myrto Barrdahl, Heiner Boeing, Krasimira Aleksandrova, Antonia Trichopoulou, et al. Tobacco smoking-associated genome-wide dna methylation changes in the epic study. Epigenomics, 8(8):599–618, 2016.

20. Michael M Mendelson, Riccardo E Marioni, Roby Joehanes, Chunyu Liu, Åsa K Hedman, Stella Aslibekyan, Ellen W Demerath, Weihua Guan, Degui Zhi, Chen Yao, et al. Association of body mass index with dna methylation and gene expression in blood cells and relations to cardiometabolic disease: mendelian randomization approach. PLoS medicine, 14(14):e1002215, 2017.

21. John K Kruschke. Rejecting or accepting parameter values in bayesian estimation. Advances in Methods and Practices in Psychological Science, 1(1):270–280, 2018.

22. Fran Supek, Matko Bošnjak, Nives Škunca, and Tomislav Šmuc. Revigo summarizes and visualizes long lists of gene ontology terms. PloS one, 6(6):e21800, 2011.

23. Latarsha J Carithers, Kristin Ardlie, Mary Barcus, Philip A Branton, Angela Britton, Stephen A Buia, Carolyn C Compton, David S DeLuca, Joanne Peter-Demchok, Ellen T Gelfand, et al. A novel approach to high-quality postmortem tissue procurement: the gtex project. Biopreservation and biobanking, 13(13):311–319, 2015.

24. Tune H Pers, Juha M Karjalainen, Yingleong Chan, Harm-Jan Westra, Andrew R Wood, Jian Yang, Julian C Lui, Sailaja Vedantam, Stefan Gustafsson, Tonu Esko, et al. Biological interpretation of genome-wide association studies using predicted gene functions. Nature communications, 6:5890, 2015.

25. Sonia Shah, Marc J. Bonder, Riccardo E. Marioni, Zhihong Zhu, Allan F. McRae, Alexandra Zhernakova, Sarah E. Harris, Dave Liewald, Anjali K. Henders, Michael M. Mendelson, Chunyu Liu, Roby Joehanes, Liming Liang, Bastiaan T. Heijmans, Peter A.C. ‘t Hoen, Joyce van Meurs, Aaron Isaacs, Rick Jansen, Lude Franke, Dorret I. Boomsma, René Pool, Jenny van Dongen, Jouke J. Hottenga, Marleen M.J. van Greevenbroek, Coen D.A. Stehouwer, Carla J.H. van der Kallen, Casper G. Schalkwijk, Cisca Wijmenga, Sasha Zhernakova, Ettje F. Tigchelaar, P. Eline Slagboom, Marian Beekman, Joris Deelen, Diana van Heemst, Jan H. Veldink, Leonard H. van den Berg, Cornelia M. van Duijn, Bert A. Hofman, André G. Uitterlinden, P. Mila Jhamai, Michael Verbiest, H. Eka D. Suchiman, Marijn Verkerk, Ruud van der Breggen, Jeroen van Rooij, Nico Lakenberg, Hailiang Mei, Maarten van Iterson, Michiel van Galen, Jan Bot, Peter van ‘t Hof, Patrick Deelen, Irene Nooren, Matthijs Moed, Martijn Vermaat, Dasha V. Zhernakova, René Luijk, Marc Jan Bonder, Freerk van Dijk, Wibowo Arindrarto, Szymon M. Kielbasa, Morris A. Swertz, Erik W. van Zwet, Daniel Levy, Nicholas G. Martin, John M. Starr, Cisca Wijmenga, Naomi R. Wray, Jian Yang, Grant W. Montgomery, Lude Franke, Ian J. Deary, and Peter M. Visscher. Improving phenotypic prediction by combining genetic and epigenetic associations. The American Journal of Human Genetics, 97(97):75–85, Jul 2015.

26. Hans D Daetwyler, Beatriz Villanueva, and John A Wooliams. Accuracy of predicting the genetic risk of disease using a genome-wide approach. PLoS One, 3(3):e3395, 2008.

27. Daniel L. McCartney, Robert F. Hillary, Anna J. Stevenson, Stuart J. Ritchie, Rosie M. Walker, Qian Zhang, Stewart W. Morris, Mairead L. Bermingham, Archie Campbell, Alison D. Murray, Heather C. Whalley, Catharine R. Gale, David J. Porteous, Chris S. Haley, Allan F. McRae, Naomi R. Wray, Peter M. Visscher, Andrew M. McIntosh, Kathryn L. Evans, Ian J. Deary, and Riccardo E. Marioni. Epigenetic prediction of complex traits and death. Genome Biology, 19(19):136, Sep 2018.

28. John Geweke. Bayesian treatment of the independent student-t linear model. Journal of applied econometrics, 8(S1):S19–S40, 1993.

29. M Erbe, BJ Hayes, LK Matukumalli, S Goswami, PJ Bowman, CM Reich, BA Mason, and ME Goddard. Improving accuracy of genomic predictions within and between dairy cattle breeds with imputed high-density single nucleotide panels. Journal of dairy science, 95(95):4114–4129, 2012.

30. Dirk Eddelbuettel and Romain François. Rcpp: Seamless R and C++ integration. Journal of Statistical Software, 40(40):1–18, 2011.

31. Gaël Guennebaud, Benoît Jacob, et al. Eigen v3. http://eigen.tuxfamily.org, 2010.

32. Douglas Bates and Dirk Eddelbuettel. Fast and elegant numerical linear algebra using the RcppEigen package. Journal of Statistical Software, 52(52):1–24, 2013.

33. John Geweke et al. Evaluating the accuracy of sampling-based approaches to the calculation of posterior moments, volume 196. Federal Reserve Bank of Minneapolis, Research Department Minneapolis, MN, USA, 1991.

34. Andrew Gelman and Donald B Rubin. Inference from iterative simulation using multiple sequences. Statistical science, pages 457–472, 1992.

35. Xavier Fernández i Marín. ggmcmc: Analysis of MCMC samples and Bayesian inference. Journal of Statistical Software, 70(70):1–20, 2016.

36. Kasper Daniel Hansen. IlluminaHumanMethylation450kanno.ilmn12.hg19: Annotation for Illumina’s 450k methylation arrays, 2016. R package version 0.6.0.

37. Steffen Durinck, Paul T. Spellman, Ewan Birney, and Wolfgang Huber. Mapping identifiers for the integration of genomic datasets with the r/bioconductor package biomart. Nature Protocols, 4:1184–1191, 2009.

38. Marc Carlson. GO.db: A set of annotation maps describing the entire Gene Ontology, 2017. R package version 3.5.0.

39. Adele M Taylor, Alison Pattie, and Ian J Deary. Cohort profile update: The lothian birth cohorts of 1921 and 1936. International Journal of Epidemiology, 47(47):1042–1042r, 2018.

40. Andy Boyd, Jean Golding, John Macleod, Debbie A Lawlor, Abigail Fraser, John Henderson, Lynn Molloy, Andy Ness, Susan Ring, and George Davey Smith. Cohort profile: the ‘children of the 90s’—the index offspring of the avon longitudinal study of parents and children. International journal of epidemiology, 42(42):111–127, 2013.

41. Abigail Fraser, Corrie Macdonald-Wallis, Kate Tilling, Andy Boyd, Jean Golding, George Davey Smith, John Henderson, John Macleod, Lynn Molloy, Andy Ness, et al. Cohort profile: the avon longitudinal study of parents and children: Alspac mothers cohort. International journal of epidemiology, 42(42):97–110, 2012.

42. Caroline L Relton, Tom Gaunt, Wendy McArdle, Karen Ho, Aparna Duggirala, Hashem Shihab, Geoff Woodward, Oliver Lyttleton, David M Evans, Wolf Reik, et al. Data resource profile: accessible resource for integrated epigenomic studies (aries). International journal of epidemiology, 44(44):1181–1190, 2015.

43. JL Min, G Hemani, G Davey Smith, C Relton, M Suderman, and John Hancock. Meffil: efficient normalization and analysis of very large dna methylation datasets. Bioinformatics, 2018.

44. Jean-Philippe Fortin, Aurélie Labbe, Mathieu Lemire, Brent W Zanke, Thomas J Hudson, Elana J Fertig, Celia MT Greenwood, and Kasper D Hansen. Functional normalization of 450k methylation array data improves replication in large cancer studies. Genome biology, 15(15):503, 2014.

